# Reproductive experience drives changes in behavior and physiology in male California mice (*Peromyscus californicus*)

**DOI:** 10.1101/2024.10.08.617085

**Authors:** Maria E. Colt, Priyanka Agarwal, David Kolb, Erica R. Glasper, Heidi S. Fisher

**Affiliations:** Department of Biology, University of Maryland, College Park, MD 20742, USA; Department of Psychology, University of Maryland, College Park, MD 20742, USA; Department of Neuroscience, The Ohio State University College of Medicine, Columbus, OH 43210, USA; Institute for Behavioral Medicine Research, The Ohio State University College of Medicine, Columbus, OH, 43210, USA; The Jackson Laboratory, Bar Harbor, ME 04609, USA

**Keywords:** anxiety-like behavior, California mouse, parental care, paternal care, recognition memory, reproductive investment, sperm production

## Abstract

Paternal experience improves memory and reduces anxiety-like behavior in males, but it is unclear whether these changes are due to mating, siring offspring, or caregiving behavior. Likewise, paternal males have larger testes, a measure of sperm production, but again, the effects of siring and caregiving experience are difficult to disentangle. We examined behavioral and physiological outcomes in three groups of male mice: 1) virgins, 2) males paired with sterile females (‘non-fathers’), and 3) experienced fathers (Experiment 1). Compared to virgins and non-fathers, experienced fathers exhibited increased recognition memory (novel object recognition) and decreased anxiety-like behavior (elevated plus maze). Virgin males, however, had smaller testes and fewer sperm compared to non-fathers. We then compared the same traits in three additional groups of male mice: 1) non-fathers, 2) non-fathers with experience caring for unrelated pups (‘pup-sensitized non-fathers’), and 3) first-time fathers, to determine whether the behavioral and physiological observations in Experiment 1 were due to siring offspring or caregiving, and how rapidly these changes occur (Experiment 2). Recognition memory and anxiety-like behavior did not differ among these three groups, suggesting that caring for a single litter does not recapitulate the behavioral changes observed in experienced fathers (Experiment 1). Despite equal mating opportunity, we observed larger testes in first-time fathers compared to non-fathers, suggesting that investment in sperm production may be more plastic than behavioral changes. Finally, we compared pup interactions in pup-sensitized non-fathers and first-time fathers. While pup-sensitized non-fathers were slower to approach pups than first-time fathers, they spent more time grooming pups, whereas first-time fathers invested more time in nest building, suggesting different caregiving behaviors in pup-sensitized males and biological fathers. Taken together, our study revealed that mating, siring, and caregiving experience contributes to changes in memory, anxiety-like behavior, and reproductive investment in males of a biparental species.

## Introduction

Across taxa, the transition to parenthood is often associated with an array of behavioral changes that improve the fitness of the parent, as well as the offspring. This is particularly well-studied in rodents (Lonstein and De Vries, 2000). Rodent mothers, for example, exhibit improved cognition and memory as well as reduced anxiety-like behavior (Leuner *et al*., 2010a; Saltzman & Ziegler, 2014; Leuner & Sabihi, 2016; Rogers & Bales, 2019), and these outcomes are often mirrored in fathers of naturally biparental species (Saltzman & Ziegler, 2014; Glasper *et al*., 2019; Been *et al*., 2021). In California mice, *Peromyscus californicus*, fathers demonstrate enhanced spatial memory on dry land maze tests compared to virgins and pup-exposed virgins (Franssen *et al*., 2011) and decreased anxiety-like behavior on elevated plus-maze tests compared to non-fathers (i.e., males paired with sterile females: Glasper *et al*., 2016; Hyer *et al*., 2016) during the mid-postpartum period. As offspring become more mobile, paternal care shifts from passive interactions with pups to those that are more active (Bester-Meredith *et al*., 1999; Marler *et al*., 2003). These shifts are accompanied by enhanced cognition and reduced anxiety-like behavior in fathers which may lead to an improved ability to locate pups (Lee *et al*., 1999) and decreased avoidance that promotes caregiving (Wartella *et al*., 2003). Despite this knowledge, the degree to which mating experience, reproductive experience (i.e., siring offspring), exposure to offspring, and caregiving experience influences these behavioral outcomes in fathers remains unclear since non-fathers are rarely tested with pups (e.g., Bardi *et al*., 2011; Glasper *et al*., 2011; Chauke *et al*., 2012; Glasper *et al*., 2016; Hyer *et al*., 2016; Hyer *et al*., 2017). In this study, we disentangle these separate experiences and assess their effect on indicators of memory and anxiety-like behavior in California mouse males.

In the biparental California mouse, both the mother and father care extensively for the offspring (Gubernick & Alberts, 1987). Mated pairs form a strong, life-long bond, resulting in social monogamy, but these mice are also genetically monogamous, meaning that only bonded pairs produce offspring (Dudley, 1974; Ribble, 1991). With high levels of paternal certainty, California mouse fathers invest significantly in raising their young (Whittingham *et al*. 1992; Xia, 1992; Kokko & Jennions, 2008) and engage in all maternal behaviors besides nursing including grooming, nest building, pup retrieval, and huddling (Dewsbury, 1985; Gubernick & Alberts, 1987; Gubernick & Nelson, 1989; Gubernick & Nordby, 1993). Experimental removal of the paternal male from the nest decreases pup survival, suggesting that, although energetically costly (West & Capellini, 2016), paternal investment is essential for offspring development (Cantoni & Brown, 1997; Gubernick & Teferi, 2000; Glasper *et al*., 2018). Therefore, in addition to the changes in memory and anxiety-like behavior in males, fatherhood is associated with increased energetic costs from providing care (Zhao *et al*., 2017); however, it is unclear if the demands of caregiving lead to a reduction in other energetically costly traits.

Life history theory predicts differential allocation in reproductive traits between promiscuous and monogamous species because monogamous individuals often form long-term pair bonds and, consequently, experience less competition for mates and more certainty of future mating opportunities (Parker, 1984). Indeed, monogamous *Peromyscus* species exhibit less investment in reproductive potential exhibiting lower sperm count, smaller relative testes size, and decreased seminiferous tubule density, compared to promiscuous species (Weber & Fisher, 2023). Furthermore, within California mouse males, virgins have smaller testes size, a robust indicator of sperm production (Parker & Pizzari, 2010), compared to non-fathers that have been paired with sterile females and first-time fathers (Harris & Saltzman, 2013; Zhao *et al*., 2017), yet the impact on sperm count is under-reported. Here, we test the prediction that males of this monogamous, biparental species reduce investment in sperm production after becoming fathers and increase investment in offspring care by examining the effects of mating, siring, and caregiving on average testis weight and sperm count.

Unlike most avian species, in which social monogamy and biparental care is common (Lack, 1968), and fish, in which greater than 50% of families that provide parental care display male-only care (Goldberg *et al*., 2020), paternal care is only observed in roughly 3–5% of mammalian species (Kleiman, 1977; Rogers and Bales, 2019). Historically, mammalian studies investigating paternal care have used traditional laboratory rodents (i.e., *Rattus*, *Mus*) that do not show spontaneous paternal behavior (Kohl & Dulac, 2018); however, the use of naturally biparental mammals has aided in the study of paternal care (e.g., California mice [Gubernick & Alberts, 1987], oldfield mice [*P. polionotus*; Margulis, 1998], prairie voles [*Microtus ochrogaster*; Thomas & Birney, 1979], mandarin voles [*Lasiopodomys mandarinus*; Smorkatcheva, 2003], Djungarian hamsters [*Phodopus sungorus*; Wynne-Edwards, 1998], Mongolian gerbils [*Meriones unguiculatus*; Ostermeyer & Elwood, 1984], common marmosets [*Callithrix jacchus*; Dixson & George, 1982], titi monkeys [*Plecturocebus donacophilus*; Fragaszy *et al*., 1982], and cotton top tamarins [*Saguinus oedipus*; Ziegler *et al*., 1996]). To disentangle the trigger for the behavioral and physiological changes previously observed in paternal males, we leveraged the California mouse system and established experimental groups of males with varying degrees of mating, siring, and caregiving experience consisting of virgin males, mated non-fathers with no exposure to pups, non-fathers that have been sensitized to pups and exhibited paternal care behaviors, first time fathers, and experienced fathers that have sired two litters.

In the first phase of this study (Experiment 1), we estimated one-trial object recognition memory, anxiety-like behavior, average testis weight, and sperm count in virgins, non-fathers, and experienced fathers. We observed that experienced fathers spent more time exploring the novel object during the novel object recognition test compared to virgins and spent more time on the open arms of the elevated plus maze compared to virgins and non-fathers. Additionally, virgins had smaller testes and less sperm than non-fathers but did not differ from experienced fathers. We therefore designed Experiment 2 to compare similar traits in two additional groups, non-fathers who had experience caring for pups, and first-time fathers, plus a second set of non-fathers without pup exposure as an interexperiment control, which together allowed us to disentangle whether changes observed in Experiment 1 were due to reproductive or caregiving experience, and how rapidly these changes occur. Additionally, we quantified caregiving behaviors in pup-sensitized non-fathers and first-time fathers in Experiment 2 to determine if any of these phenotypes were correlated with interaction with pups and to determine to what extent these two groups differed in pup interactions.

## Materials and Methods

### Animals

We obtained sexually mature, gonadally-intact male and female California mice and age-matched females that were sterilized by tubal ligation from the Peromyscus Genetic Stock Center at the University of South Carolina. We provided the mice *ad libitum* access to food and water and housed them at 22 °C on a 16L:8D cycle in accordance with guidelines established by the Institutional Animal Care and Use Committee at the University of [anonymized for review] (protocols R-OCT-17-41 and R-JUN-21-40).

In Experiment 1, we randomly assigned virgin male California mice to one of three experimental groups: virgins, non-fathers, and experienced fathers. We housed *virgins* (N = 16) with another virgin male for the duration of the study, thus virgins did not mate nor reproduce (Figure 1) but had social exposure. We housed *non-fathers* (N = 8) with sterile, tubally-ligated females, which allows for normal estrous cycling and mating behavior, thus non-fathers had the opportunity to mate but not reproduce and had no exposure to pups. We housed *experienced fathers* (N = 15) with a reproductive female and began data collection while they were rearing their second litter, thus fathers were able to mate, reproduce, and provide parental care (Figure 1A).

**Figure 1.**
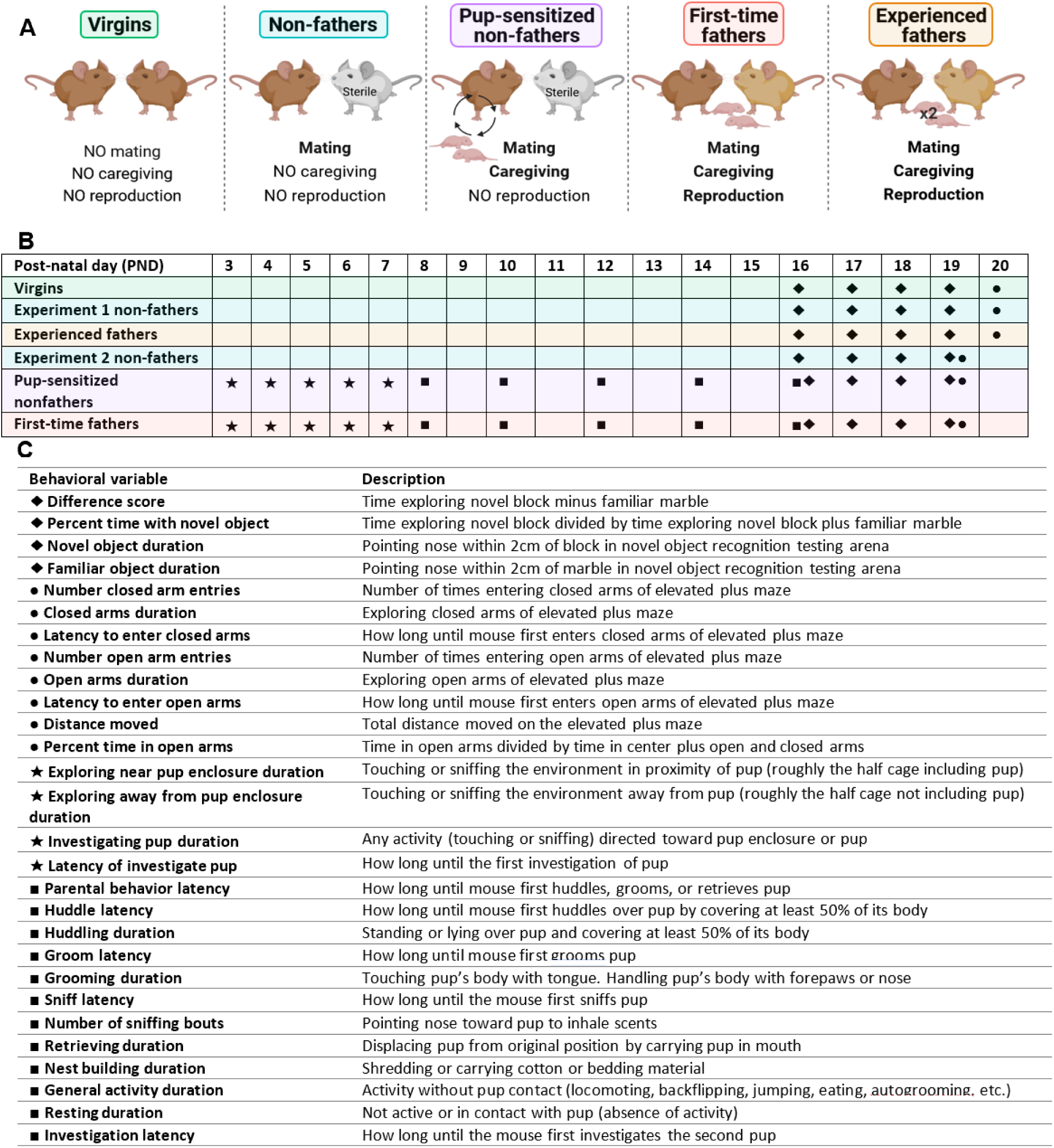
Experimental design. **(A)** Experimental treatment groups. Virgin males were socially housed with another virgin male for the duration of the study, thus virgins did not mate nor reproduce. Non-fathers were housed with a sterile, tubal-ligated females which allows for normal estrous cycling and mating behavior, thus non-fathers had the opportunity to mate but not reproduce and had no exposure to pups. Pup-sensitized non-fathers were housed with sterile females, however, these males interacted with unrelated pups to test the effect of caregiving in the absence of reproduction. First-time fathers were housed with a reproductive female and their first litter offspring, whereas experienced fathers were housed with a reproductive female and their second litter, thus fathers were able to mate, reproduce, and provide parental care. **(B)** Timeline of behavioral tests in Experiment 1 and Experiment 2. Shading is associated with treatment group and empty squares denote no testing; ★, Pup-exposure assay; ■, Caregiving assay; ◆, Novel object recognition test; ●, Elevated plus maze test. **(C)** Description of scored observations for behavioral testing.

In Experiment 2, we randomly assigned virgin male California mice to one of three experimental groups: non-fathers, pup-sensitized non-fathers, and first-time fathers. As in Experiment 1, we housed *non-fathers* (N = 10) with sterile females. We also housed *pup-sensitized non-fathers* (N = 9) with sterile females, however, these males interacted with unrelated pups to test the effects of caregiving in the absence of reproduction (see ‘Pup-exposure assay’ and ‘Caregiving assay’ below). We housed *first-time fathers* (N = 17) with a gonadally-intact female and their first litter offspring. Since California mouse females will remate (Valentino *et al*., 2021), we paired females with a second experimental male two weeks after separation from the first male, and therefore considered female experience in our statistical analyses in Experiment 2 (Figure 1A).

### Ethical Note

We conducted a power analysis before beginning our experiments to determine the appropriate number of male mice needed, ensuring we achieved sufficient statistical power while minimizing the number of animals used. To minimize stress on the mice, we reduced unnecessary handling and ensured that only one observer was present in the room throughout the experiment. During the behavioral assays, we closely monitored the male mice for any signs of aggression toward the pups or distress. We did not observe any instances of aggression toward the pups. After each assay, we carefully inspected both the males and pups for injuries and found none. At the end of the study, we humanely euthanized all male mice and pups according to approved protocols. We followed ethical guidelines throughout the study, with all procedures approved by the Institutional Animal Care and Use Committee at the University of [anonymized for review] (protocols R-OCT-17-41 and R-JUN-21-40).

### Behavioral Tests

#### Novel object recognition test: Experiments 1 and 2

To estimate recognition memory in all males, we performed novel object recognition tests under red-light illumination during the dark phase of the light cycle from postnatal day (PND) 16 to 19 for first-time fathers and experienced fathers following methods of Agarwal *et al*. (2020). We tested age-matched virgins, non-fathers, and pup-sensitized non-fathers at the same time as the first-time fathers and experienced fathers (Figure 1). Briefly, on day 1 of testing, we placed the male in a holding cage and transferred it to the behavioral testing room to habituate to the room for one hour, then returned the male to its home-cage. On days 2 and 3, we transferred the male to the behavioral testing room in a holding cage and placed it in the open field arena (43 cm x 43 cm x 30 cm) for five minutes, then returned the male to its home-cage. On day 4, we transferred the male to the behavioral testing room in a holding cage and placed it in the open field for five minutes where two identical marbles were secured to the surface (i.e., acquisition phase), then placed the male back in the holding cage for 15 minutes.

Lastly, we returned the male to the open field for five minutes where a third identical marble and a one-inch wooden block were now secured to the surface (i.e., recognition phase). We counterbalanced the location of the block and marble between mice to ensure there was not a preference for a particular side of the open field. To remove potential scent cues, we cleaned the open field arena and marbles after each phase with 70% ethanol or Bio-Clean (Stanbio Laboratory, Boerne, TX, USA), and used a new wooden block for each recognition phase. We used a random number generator to determine the order individuals were tested each day. We recorded novel object recognition tests from above the testing arena using a ceiling-mounted camera (Noldus, Leesburg, VA, USA). We determined time spent exploring (i.e., nose pointed within 2 cm of the object) the familiar (i.e., marble) and novel (i.e., wooden block) objects using EthoVision® XT 14 behavioral tracking software (Noldus, Leesburg, VA, USA). We excluded males that spent less than 20 seconds exploring either object (N = 5 Virgins, 2 Non-fathers [Experiment 1], 4 Experienced fathers, and 4 Non-fathers [Experiment 2]). We calculated difference scores (i.e., time exploring novel block minus familiar marble) and percent time exploring the novel object (i.e., time exploring novel block divided by time exploring novel block plus familiar marble).

#### Elevated plus maze test: Experiments 1 and 2

To estimate anxiety-like behavior in all males, we conducted elevated plus maze tests under red-light illumination during the dark phase of the light cycle on PND 20 in Experiment 1 and PND 19 in Experiment 2 for first-time fathers and experienced fathers following methods of Glasper *et al*., (2016), Hyer *et al*., (2016), and Hyer *et al*., (2017). We tested age-matched virgins, non-fathers, and pup-sensitized non-fathers at the same time as the first-time fathers and experienced fathers (Figure 1). Specifically, we transferred the male to the behavioral testing room in a holding cage (a random number generator was used to determine the order), placed the male in the center of the elevated plus maze facing an open arm, and recorded its behavior for five minutes. The elevated plus maze was 75 cm above the ground and the arms were 112 cm by 10 cm. We recorded elevated plus maze behavior from above using a ceiling-mounted camera (Noldus, Leesburg, VA, USA). We measured the latency to enter each arm, total time spent in arms, number of arm entries, and total distance traveled using EthoVision® XT 14 behavioral tracking software (Noldus, Leesburg, VA, USA). We excluded males that failed to move for more than 40% of the test (N = 1 Virgin and 1 Experienced father) and those that fell off the maze more than twice (N = 2 Virgins, 3 Non-fathers [Experiment 1], 2 Experienced fathers, 1 Non-father [Experiment 2], and 1 First-time father). We calculated percent time in open arms (i.e., time in open arms divided by time in open and closed arms).

#### Pup-exposure assay: Experiment 2

To expose males in the ‘pup-sensitized non-fathers’ group to unrelated pups and to score pup-exploration behavior in pup-sensitized non-fathers and first-time fathers, we conducted pup-exposure assays under red-light illumination during the dark phase of the light cycle at approximately 13:00 hours daily from PND 3 to PND 7 in first-time fathers. We age-matched each pup-sensitized non-father with a first-time father and performed the pup-exposure assay on the pup-sensitized non-fathers at the same time as their ‘yoked’ fathers (Figure 1; see photo in Figure S1A). We used a random number generator to determine the order individuals were tested each day. From above the cage, we recorded pup-exposure assays using remotely controlled NoIR cameras (Raspberry Pi Foundation, Cambridge, United Kingdom; Weber & Fisher, 2019). Briefly, we first removed the female and any offspring from the male’s cage, then exposed the male to one pup secured within a mesh enclosure for 10 minutes; first-time fathers were exposed to one of their own pups and pup-sensitized non-fathers to their assigned unrelated pup.

We then scored the following male behaviors during the pup-exposure assay using the Behavioral Observation Research Interactive Software (BORIS, version 7.13; Friard and Gamba, 2016): duration of exploration near enclosure (i.e., touching or sniffing the environment in the half of the cage including the pup), exploration away from enclosure (i.e., touching or sniffing the environment in the half of the cage excluding the pup), resting, pup investigation (i.e., touching or sniffing directed toward the pup), and recorded latency to pup investigation (Lambert *et al*., 2013). All videos were scored by a primary observer blind to individual identification and verified by a randomly assigned second observer. We compared the primary observer’s scores to the second observer’s scores and if found to be different, both observers would re-score the video until their results were concordant. To demonstrate agreement between coders and confidence in our results, we performed intra-class correlations for each behavior between the scores of four independent coders on a subset of videos and ensured they were above the 0.75 threshold (Table S1). We removed resting from the pup-exposure assay analysis because the intra-class correlation for this behavior was below the 0.75 threshold. To confirm consistency in scoring, the primary observer re-scored five videos and we compared the old scores to the new scores using intra-class correlations for each behavior and ensured they were above the 0.75 threshold (Table S1). Our reported analyses are based on the primary observer’s scores.

#### Caregiving assay: Experiment 2

To quantify caregiving behavior in pup-sensitized non-fathers and first-time fathers, we conducted caregiving assays under red-light illumination during the dark phase of the light cycle at approximately 13:00 hours following methods of Bendesky *et al*. (2017) every other day from PND 8 to PND 16 in first-time fathers. We tested pup-sensitized non-fathers at the same time as their ‘yoked’ fathers (Figure 1; see photo in Figure S1B), used a random number generator to determine the order individuals were tested, and recorded caregiving assays from above the cage using remotely controlled NoIR cameras (Raspberry Pi Foundation, Cambridge, United Kingdom; Weber & Fisher, 2019). We first removed the female and any offspring from the male’s cage and placed a 2.1 g cotton nestlet in the cage to assess nest quality and building behavior. At minute 30, we placed one of the first-time father’s pups in the corner of the cage farthest away from the male. For pup-sensitized non-fathers, we placed one unrelated pup from the same father as in the pup-exposure assay in the corner farthest from the male. Using BORIS, we measured the duration and latency to groom, sniff, retrieve, and huddle over the pup, as well as the duration of general activity (i.e., activity without contact with pup) and nest building until minute 50 (see detailed scoring methods within ‘Pup-exposure assay’ above and inter-rater reliability results in Table S2). At minute 50, we removed the first pup, placed a second pup in the corner of the cage for two minutes, and measured the latency to investigate and retrieve the pup. We then measure nest quality using the scoring system described in Bendesky *et al*., (2017).

### Physiology

#### Reproductive traits: Experiments 1 and 2

We estimated average testis weight and sperm count following methods of Weber & Fisher (2023) in all males. Briefly, following the completion of the behavioral tests, we anesthetized each male with isoflurane, recorded its body mass, and euthanized the mouse via rapid decapitation. Next, we removed both testes and the left caudal epididymis. To calculate average testis weight, we weighed both testes from each mouse and calculated the average. To estimate sperm count, we made three incisions in the epididymis and incubated the tissue at 37°C for 1 h with 300 rpm agitation in 1000 μl of Modified Human Tubal Fluid (Irvine Scientific, Santa Ana, CA, USA) supplemented with 5 mg/ml of Probumin bovine serum albumin (Millipore Sigma, Darmstadt, Germany). We inverted the tube of suspended sperm 3X to homogenize, pipetted 95 μl into another tube containing 5 μl of formalin, pipetted gently to mix, and aliquoted 10 μl onto a hemocytometer (Paul Marienfeld GmbH & Co. KG, Lauda-Königshofen, Germany). We then imaged five hemocytometer grids at 250x magnification using an AxioCam 105c camera on an AxioPlan microscope (Carl Zeiss AG, Jena, Germany), and used ImageJ (version 1.53a; Schindelin *et al*., 2012) to estimate sperm count following the World Health Organization guidelines (WHO, 2010).

### Statistical Analyses

We conducted all statistical analyses using R Statistical Software (version 4.2.0, R Core Team 2022). To examine if behaviors on the novel object recognition and elevated plus maze tests differed by experience, we first performed separate univariate linear models (LM) for each behavior using the R ‘lm’ function (see univariate LM results in Tables S3-6). We then performed LMs with fixed effects included to test the effect of experience and age on novel object recognition and elevated plus maze behaviors. We included male age as a fixed effect because it has been suggested that paternal motivation may be age-dependent (Nguyen *et al*., 2020). In Experiment 2, we also included female mating history (i.e., number of males to which the female had been paired) in each LM because it has been shown that maternal behaviors in rodents are altered by previous reproductive experience (Nasu *et al*., 2022; Dunlap *et al*., 2020; Rymer & Pillay, 2012; Cohen-Salmon, 1987) and that California mouse fathers will spend more time with pups when the mother is not with them (Dudley, 1974). For each behavior, we used the “MuMIn” R package (Burnham & Anderson, 2002; Bartoń, 2022) dredge function to generate LMs using all combinations of covariates and determined the best fitting model for each behavior by the selecting the model with the lowest corrected Akaike information criterion (AICc).

For the novel object recognition test in Experiment 1, the best fitting models for difference score (i.e., time exploring novel block minus familiar marble) and time with the novel and familiar objects included no covariates. However, the best fitting model for percent time with the novel object included experience as a predictor. The best fitting models for all Experiment 2 novel object recognition test behaviors (i.e., difference score, percent time exploring the novel object, and time with the block and marble) included no covariates.

For the elevated plus maze test in Experiment 1, the best fitting models for the number of open arm entries and distance moved included no covariates. However, the best fitting model for the number of closed arm entries included age as a predictor, and best fitting models for time spent in the closed and open arms and percent time in the open arms included experience as a predictor. We removed female mating history as a covariate from the Experiment 2 elevated plus maze analysis because only one behavior (i.e., distance moved) had a model that included female mating history as a predictor with a delta value less than 2 and this model had relatively low weight. For the elevated plus maze test in Experiment 2, the best fitting models for all behaviors included no covariates.

To examine if behaviors during the pup-exposure and caregiving assays differed among the pup-sensitized non-fathers and first-time fathers, we completed separate linear mixed effects models (LMM) for each behavior using the “lme4” package in R (Luke, 2016). We considered male experience group and age as fixed effects and male ID as a random effect to account for repeated measures. We did not include female mating and reproduction history as a fixed effects because we found that male behavior did not differ significantly when paired with a female with previous mating and reproduction experience (paired t-tests: *P* > alpha when adjusted for multiple comparisons). For each behavior, we used the “MuMIn” R package (Burnham & Anderson, 2002; Bartoń, 2022) dredge function to generate LMMs using all combinations of covariates and determined the best fitting model for each behavior by selecting the model with the lowest corrected AICc (see full results in Tables S9-10).

For the pup-exposure assay, the best fitting models for the duration of pup investigation and exploration near and away from the pup enclosure, as well as the number of bouts of pup investigation and exploration away from the enclosure, and the latency to investigate the pup included experience group as a fixed effect. The best fitting model for the number of bouts of exploration near the pup enclosure included experience group and age as fixed effects.

For the caregiving assay, the best fitting model for sniff latency included no fixed effects. The best fitting models for groom and investigate latency, and number of sniffing bouts and retrieving duration included experience as a fixed effect. Finally, best fitting models for the duration of huddling, grooming, nest building, and general activity included experience and male age as fixed effects. Because the random factor subject ID did not contribute to variation in huddle latency, we reverted to a linear model. The best fitting model for huddle latency included experience as a covariate.

To determine if nest quality differed between nests of pup-sensitized non-fathers and first-time fathers, we performed Student’s T-tests between scores for each PND.

To examine if reproductive traits differed by experience group, we conducted separate univariate LMs for average testis weight and sperm count (see univariate LM results in Tables S7-8). We then performed LMs with fixed effects included to test the effect of experience, age, and body weight on average testis weight and sperm count since age and overall body weight can be positively associated with sperm count (reviewed in Hook & Fisher, 2020). We determined the best fitting model for each trait by selecting the lowest AICc value. However, in Experiment 1, the two models with the lowest AICc for average testis weight were similarly weighted, so we report here the second-best fitting model which included experience and body weight as effects. The best fitting model for sperm count included experience as a predictor. In Experiment 2, the best fitting model for average testis weight included experience and age as predictors, whereas for sperm count the best-fitting model included no covariates.

## Results

### Experiment 1

#### Novel object recognition

When males were allowed to explore the arena during novel object recognition testing, we observed that experienced fathers spent a greater percentage of time exploring the novel object, compared to virgins but not non-fathers; virgins and non-fathers explored the novel object similarly (Figure 2A; Table 1). Moreover, males from the experience groups did not differ in difference score or time with novel and familiar objects (Figure S2; Table 1).

**Figure 2.**
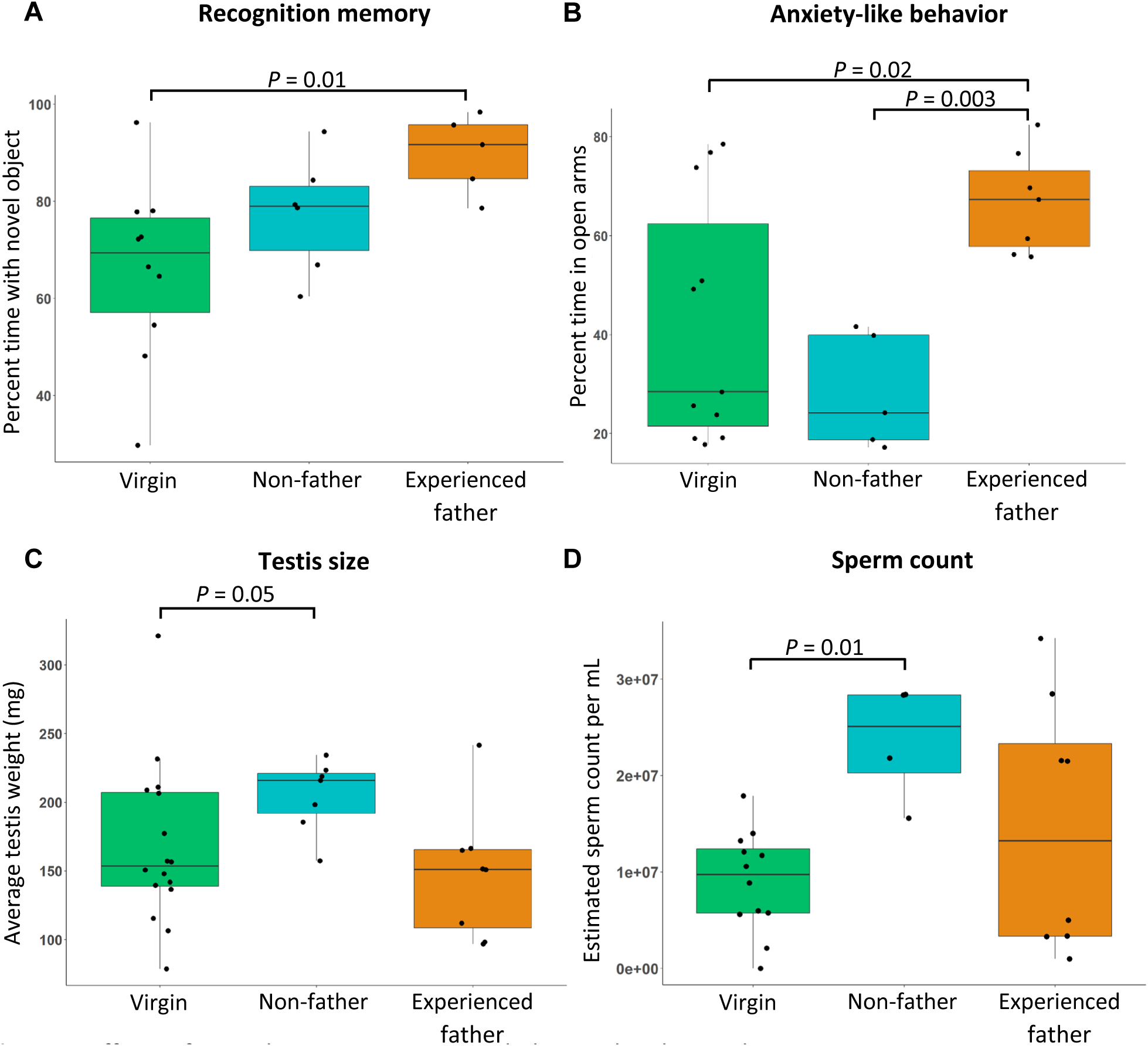
Effects of reproductive experience on behavioral and reproductive traits in Experiment 1 males. **(A)** Percent time with novel object during the novel object recognition test, a positive indicator of recognition memory, **(B)** percent time in open arms during the elevated plus maze test, a negative indicator of anxiety-like behavior, **(C)** average testis weight and **(D)** sperm count, both positive indicators of sperm production, of virgins (green), non-fathers (teal), and experienced fathers (orange). Points represent individuals, boxes represent the first and third quartiles, center lines represent medians, and whiskers extend to 1.5 times the inter-quartile ranges. P-values < 0.05 are indicated.

**Table 1.** Results from best fitting linear models comparing the effect of predictors on novel object recognition and elevated plus test outcomes, and reproductive traits in Experiment 1.

| Response | Predictors (P-value pairwise comparison) | Estimated mean | P value | Adjusted R<sup>2</sup> |
| --- | --- | --- | --- | --- |
| Novel object recognition test |  |  |  |  |
| Difference score | Intercept | 55.32 |  |  |
| Percent time with novel object | Experience: Virgin (v. Non-father) | 66.09 | 0.16 | 0.25 |
|  | Experience: Non-father (v. Experienced father) | 77.35 | 0.19 |  |
|  | Experience: Experienced father (v. Virgin) | 89.82 | 0.01 |  |
| Time with novel object | Intercept | 75.11 |  |  |
| Time with familiar object | Intercept | 19.80 |  |  |
| Elevated plus maze test |  |  |  |  |
| Number closed arm entries | Age | 0.10 | 0.05 | 0.13 |
| Time in closed arms | Experience: Virgin (v. Non-father) | 158.65 | 0.16 | 0.34 |
|  | Experience: Non-father (v. Experienced father) | 199.28 | 0.002 |  |
|  | Experience: Experienced father (v. Virgin) | 91.54 | 0.02 |  |
| Number open arm entries | Intercept | 47.48 |  |  |
| Time in open arms | Experience: Virgin (v. Non-father) | 115.55 | 0.23 | 0.32 |
|  | Experience: Non-father (v. Experienced father) | 80.38 | 0.003 |  |
|  | Experience: Experienced father (v. Virgin) | 183.85 | 0.02 |  |
| Distance moved | Intercept | 4527.40 |  |  |
| Percent time in open arms | Experience: Virgin (v. Non-father) | 42.13 | 0.20 | 0.33 |
|  | Experience: Non-father (v. Experienced father) | 28.35 | 0.003 |  |
|  | Experience: Experienced father (v. Virgin) | 66.80 | 0.02 |  |
| Reproductive traits |  |  |  |  |
| Average testis weight | Experience: Virgin (v. Non-father) | 44.59 | 0.05 | 0.18 |
|  | Experience: Non-father (v. Experienced father) | 90.6 | 0.08 |  |
|  | Experience: Experienced father (v. Virgin) | 45.29 | 0.98 |  |
|  | Body weight | 0.003 | 0.05 |  |
| Sperm count | Experience: Virgin (v. Non-father) | 8993006 | 0.01 | 0.22 |
|  | Experience: Non-father (v. Experienced father) | 23539928 | 0.12 |  |
|  | Experience: Experienced father (v. Virgin) | 14812499 | 0.16 |  |

Note: When intercept-only, null model is the best fit model, only the estimate of the intercept is reported. The experimental group in parentheses is the group being compared to get P-value. P-values < 0.05 are bolded.

#### Elevated plus maze

When males were allowed to navigate the open and closed arms of the elevated plus maze, we observed that experienced fathers spent a greater percentage of time in the open arms as well as more total time in the open arms and less time in the closed arms compared to virgins and non-fathers, whereas virgins and non-fathers explored similarly (Figure 2B; Figure S3; Table 1). Males from the experience groups did not differ in the number of arm entries or distance moved on the elevated plus maze (Figures S3-4; Table 1). Furthermore, we found that male age was positively associated with the number of closed arm entries (Figure S5; Table 1).

#### Reproductive traits

We found that virgins had lower average testis weights (Figure 2C) and sperm counts (Figure 2D) compared to non-fathers, whereas experienced fathers did not differ from virgins or non-fathers (Table 1). Additionally, we observed a positive association between body weight and average testis weight (Figure S6; Table 1) and a positive correlation between average testis weight and sperm count (cor = 0.47, *P* = 0.02; Figure S7).

### Experiment 2

#### Novel object recognition

We observed that non-fathers, pup-sensitized non-fathers, and first-time fathers explored the novel object recognition testing arena similarly (Figure 3A; Figure S8; Table 2). Additionally, we found no effect of female mating history and male age on novel object recognition (Table 2).

**Figure 3.**
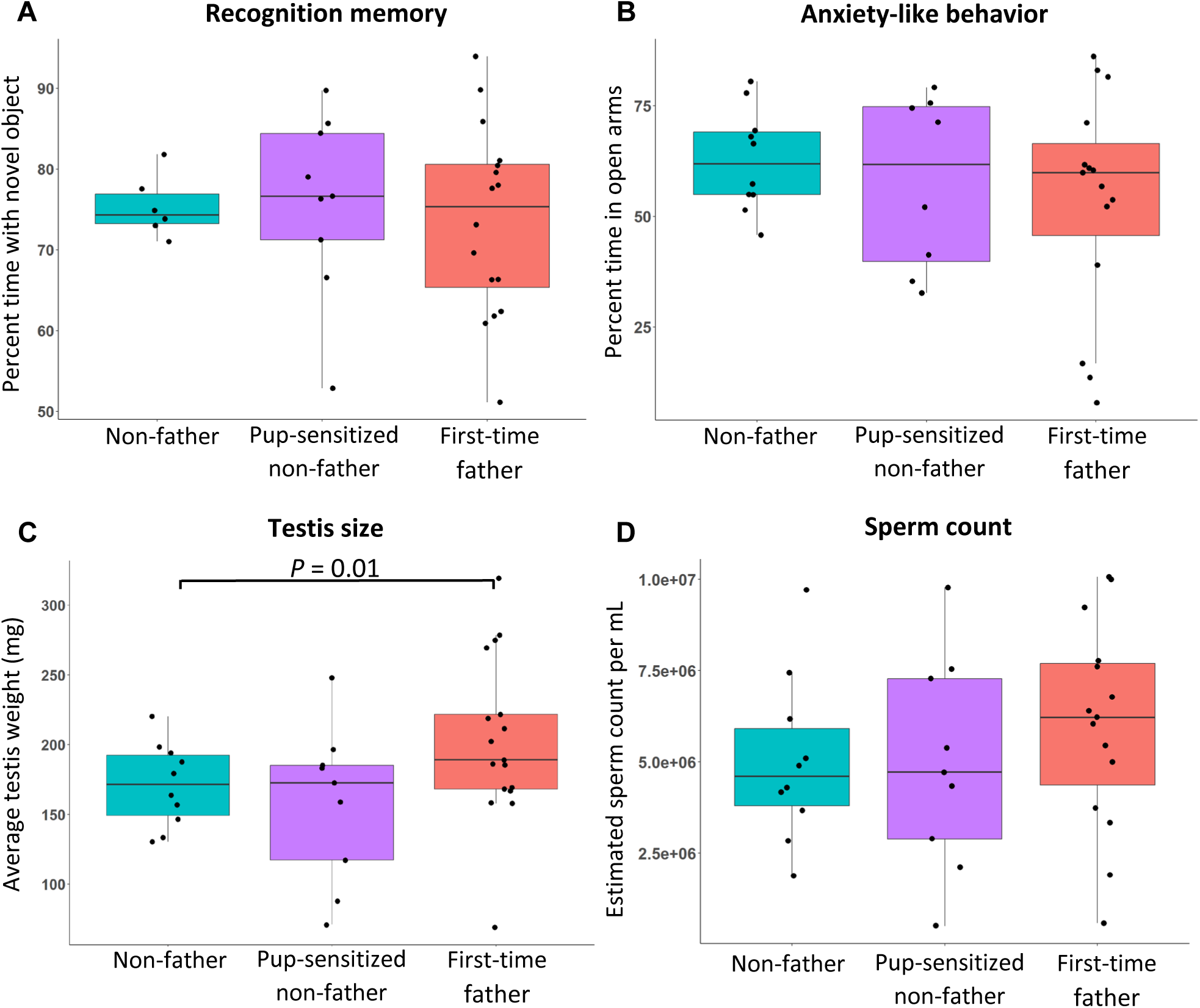
Effects of reproductive experience on behavioral and reproductive traits in Experiment 2 males. **(A)** Percent time with novel object during the novel object recognition test, a positive indicator of recognition memory, **(B)** time in open arms during the elevated plus maze test, a negative indicator of anxiety-like behavior, **(C)** average testis weight and **(D)** sperm count, both positive indicators of sperm production, of non-fathers (teal), pup-sensitized non-fathers (purple), and first-time fathers (red). Points represent individuals, boxes represent the first and third quartiles, center lines represent medians, and whiskers extend to 1.5 times the inter-quartile ranges. P-values < 0.05 are indicated.

**Table 2.** Results from best fitting linear models comparing the effect of predictors on novel object recognition and elevated plus test outcomes, and reproductive traits in Experiment 2.

| Response | Predictors (P-value pairwise comparison) | Estimated mean | P value | Adjusted R<sup>2</sup> |
| --- | --- | --- | --- | --- |
| **Novel object recognition test** |  |  |  |  |
| Difference score | Intercept | 47.36 |  |  |
| Percent time with novel object | Intercept | 74.61 |  |  |
| Time with novel object | Intercept | 71.04 |  |  |
| Time with familiar object | Intercept | 23.68 |  |  |
| **Elevated plus maze test** |  |  |  |  |
| Number closed arm entries | Intercept | 18.46 |  |  |
| Time in closed arms | Intercept | 90.70 |  |  |
| Latency to enter closed arms | Intercept | 40.77 |  |  |
| Number open arm entries | Intercept | 27.33 |  |  |
| Time in open arms | Intercept | 175.03 |  |  |
| Latency to enter open arms | Intercept | 1.82 |  |  |
| Distance moved | Intercept | 4726.97 |  |  |
| Percent time in open arms | Intercept | 65.40 |  |  |
| **Reproductive traits** |  |  |  |  |
| Average testis weight | Experience: Non-father (v. First-time father) | -47.87 | **0.01** | 0.18 |
|  | Experience: Pup-sensitized non-father (v. Non-father) | -19.83 | 0.34 |  |
|  | Experience: First-time father (v. Pup-sensitized non-father) | 17.52 | 0.08 |  |
|  | Age | 1.11 | **0.03** |  |
| Sperm count | Intercept | 5438290 |  |  |

Note: When intercept-only, null model is the best fit model, only the estimate of the intercept is reported. Experience in parentheses is the group being compared to get P-value. P-values < 0.05 are bolded.

#### Elevated plus maze

We observed that non-fathers, pup-sensitized non-fathers, and first-time fathers explored the elevated plus maze similarly (Figure 3B; Figures S9-11; Table 2). We observed no effect of male age on elevated plus maze behaviors (Table 2).

#### Reproductive traits

We found that non-fathers had lower average testis weights, compared to first-time fathers, but pup-sensitized non-fathers did not differ from non-fathers or first-time fathers (Figure 3C; Table 2). Furthermore, we found that non-fathers, pup-sensitized non-fathers, and first-time fathers did not differ in sperm count (Figure 3D; Table 2). We also found a positive association between age and average testis weight (Figure S12; Table 2) and a positive correlation between average testis weight and sperm count (cor = 0.34, *P* = 0.52; Figure S13).

#### Pup exposure assay

Pup-sensitized non-fathers and first-time fathers did not differ in any behaviors measured when introduced to a pup within an enclosure in the pup-exposure assay including exploration near pup enclosure duration, number of bouts of exploration near pup enclosure, exploration away from pup enclosure duration, number of bouts of exploration away from pup enclosure, investigating pup duration, number of investigation of pup bouts, and latency to investigate pup (Figures S14-15; Table S9).

#### Caregiving assay

When males were allowed to directly interact with the pups during the caregiving assay, pup-sensitized non-fathers took longer to investigate pups compared to first-time fathers (Figure 4A; Table 3). Additionally, we observed that pup-sensitized non-fathers spent more time grooming pups (Figure 4B), but less time building a nest (Figure 4D) compared to first-time fathers (Table 3). Our models also revealed a positive association between the male age and grooming duration (Figure S16; Table 3). Conversely, we found a negative association between male age and huddling (Figure S17) and nest building duration (Figure S18; Table 3; additional Figures S19-24). Lastly, using a Student’s T-test, we determined that first-time fathers built higher quality nests compared to pup-sensitized non-fathers on PND 10 and 12 (Figures 4E and 4F).

**Figure 4.**
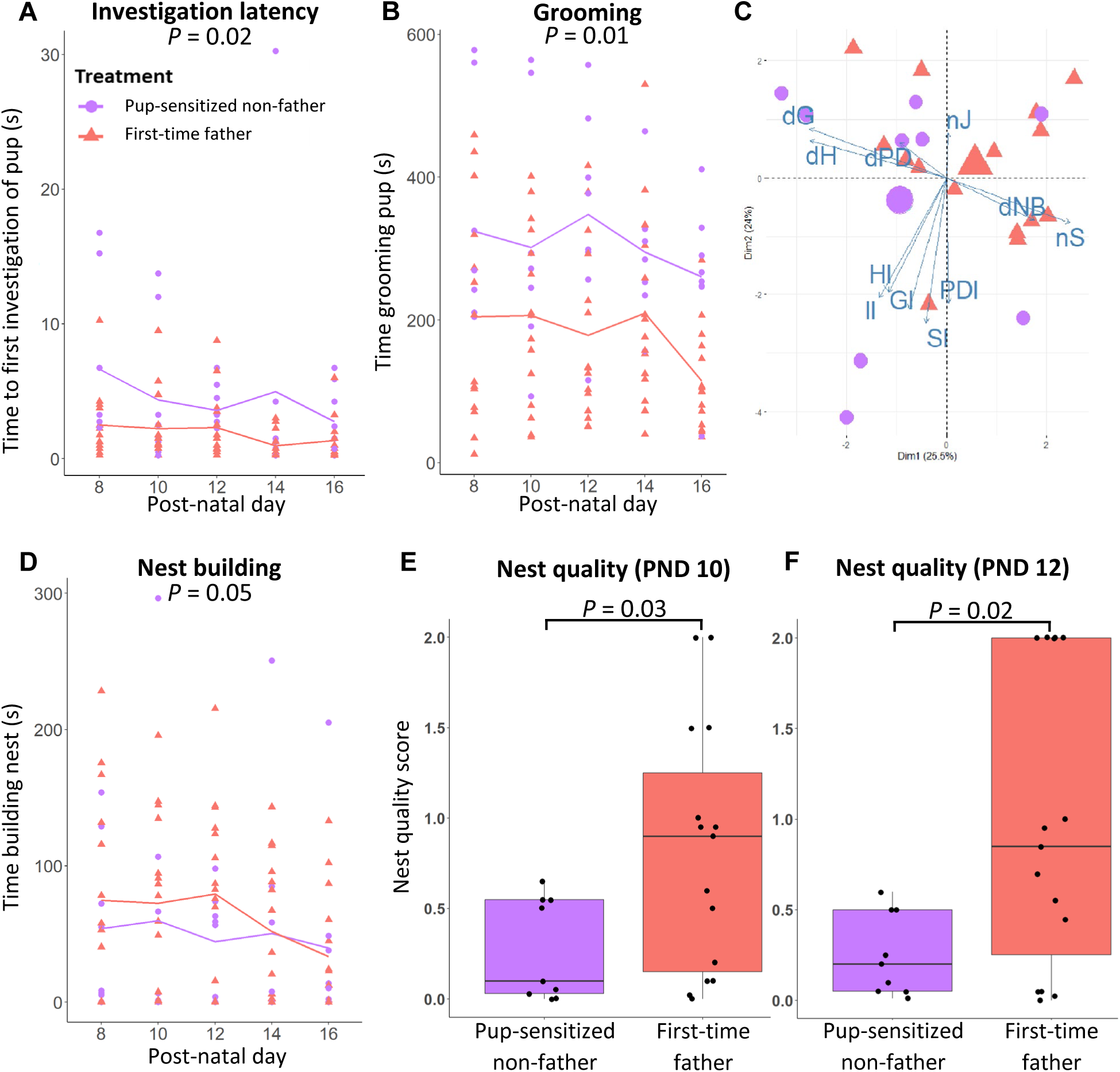
Paternal care behaviors in pup-sensitized non-fathers (purple) and first-time fathers (shown in red). (A) Latency to investigate pups during caregiving assay, where points represent individuals and lines represent treatment means. **(B)** Duration of pup grooming among the same males during caregiving assay. **(C)** Principal component analysis biplot of pup-exposure assay and caregiving assay behaviors. Purple triangles represent pup-sensitized non-fathers and red circles represent first-time fathers. The large triangle and circle represent the mean of male experience groups, and the blue arrows represent vectors for pup-exposure and caregiving assay behaviors. Vectors for duration of grooming (dG), huddling (dH), nest building (dNB), pup-directed activity (dPD), number of jumps/backflips (nJ), sniffing bouts (nS) are labeled. **(D)** Nest building duration in pup-sensitized non-fathers and first-time fathers during caregiving assay, and nest quality scores between pup-sensitized non-fathers and first-time fathers on **(E)** PND 10 and **(F)** PND 12.

**Table 3.** Results of best fitting linear mixed effects models for behaviors observed in caregiving assay.

| Response | Fixed effects | Estimate | SE | t-value | P value | Random effect | Variance | SD |
| --- | --- | --- | --- | --- | --- | --- | --- | --- |
| Huddle latency | Intercept | 5.56 | 0.63 | 8.83 | <0.001 |  |  |  |
|  | Experience | 1.04 | 1.03 | 1.01 | 0.32 |  |  |  |
| Huddling duration | Intercept | 1175.14 | 378.97 | 3.10 | 0.01 | Subject | 19863 | 140.9 |
|  | Experience | 8.79 | 71.77 | 0.12 | 0.91 | Residual | 26482 | 162.7 |
|  | Age | -4.93 | 2.31 | -2.14 | **0.05** |  |  |  |
| Groom latency | Intercept | 3.57 | 0.35 | 10.356 | <0.001 | Subject | 0.79 | 0.89 |
|  | Experience | 0.68 | 0.56 | 1.21 | 0.24 | Residual | 4.94 | 2.22 |
| Grooming duration | Intercept | -285.68 | 219.16 | -1.3 | 0.20 | Subject | 6402 | 80.01 |
|  | Experience | 129.57 | 41.24 | 3.14 | **0.01** | Residual | 9449 | 97.21 |
|  | Age | 2.85 | 1.33 | 2.14 | **0.04** |  |  |  |
| Sniff latency | Intercept | 1.77 |  |  |  |  |  |  |
| Number of sniffing bouts | Intercept | 18.79 | 2.22 | 8.46 | <0.001 | Subject | 36.75 | 6.06 |
|  | Experience | -4.65 | 3.63 | -1.28 | 0.21 | Residual | 185.75 | 13.63 |
| Retrieving duration | Intercept | 1.52 | 2.45 | 0.62 | 0.54 | Subject | 77.07 | 8.78 |
|  | Experience | 3.79 | 4.01 | 0.95 | 0.35 | Residual | 66.13 | 8.13 |
| Nest building duration | Intercept | 392.14 | 112.36 | 3.49 | 0.002 | Subject | 1618 | 40.23 |
|  | Experience | -43.20 | 21.02 | -2.06 | **0.05** | Residual | 2654 | 51.52 |
|  | Age | -2.00 | 0.68 | -2.93 | **0.01** |  |  |  |
| General activity duration | Intercept | -33.02 | 315.78 | -0.11 | 0.92 | Subject | 12584 | 112.2 |
|  | Experience | -80.01 | 58.93 | -1.36 | 0.20 | Residual | 21526 | 146.7 |
|  | Age | 2.07 | 1.92 | 1.08 | 0.29 |  |  |  |
| Investigate latency | Intercept | 1.85 | 0.62 | 2.98 | 0.01 | Subject | 3.41 | 1.85 |
|  | Experience | 2.57 | 1.02 | 2.54 | **0.02** | Residual | 11.90 | 3.45 |

Note: When intercept-only, null model is the best fit model, only the estimate of the intercept is reported. 'Experience' compares pup-sensitized non-fathers and first-time fathers, and P-values < 0.05 are bolded.

## Discussion

In Experiment 1, we observed that in the biparental California mouse, males that had cared for two litters (i.e., experienced fathers) spent more time exploring the novel object during the novel object recognition test compared to virgin males housed with other males and more time in the open arms of the elevated plus maze compared to virgins and males with mating experience but no offspring (i.e., sexually experienced non-fathers). These data suggest that paternal experience, but not mating, is associated with increased recognition memory and decreased anxiety-like behavior. In contrast, we found that virgins exhibited smaller testes and produced less sperm than non-fathers but did not differ from experienced fathers, suggesting that fathers may reduce investment in sperm production and shift toward caregiving. However, it was unclear from this first experiment if the behavioral changes we observed in experienced fathers were a result of reproductive experience (i.e., siring) or caregiving experience, and how quickly these phenotypes emerged. Therefore, in Experiment 2, we compared these same behavioral and reproductive traits in a second cohort of non-fathers as a cross-experiment control, as well as in pup-sensitized non-fathers and first-time fathers that had sired and cared for a single litter. Here, we observed that the three groups of males did not differ in traits associated with recognition memory or anxiety-like behavior, yet we did find that non-fathers with no exposure to pups had smaller testes compared to first-time fathers but did not differ from pup-sensitized non-fathers. This second experiment also allowed us to test whether the previously measured phenotypes were correlated with the type or amount of pup-interaction behaviors in pup-sensitized non-fathers and first-time fathers, which we found minimal evidence of, and how the two groups of males differed in how they interacted with pups. We observed that pup-sensitized non-fathers were slower to approach pups than first-time fathers, but eventually spent more time grooming pups, whereas fathers spent more time nest building. Although we cannot exclude the possibility that exposure to females at different reproductive stages may impact our results (Leuner *et al*., 2010b; Glasper *et al*., 2013), our findings suggest that the transition to fatherhood in California mouse males is associated with changes in behavior towards pups, recognition memory, and anxiety-like behavior, and may reflect changes in sperm production.

Across taxonomic groups, social interactions between individuals, including parents that care for offspring, are linked to improved cognition and emotional regulation (Saltzman & Ziegler, 2014; Horrell *et al*., 2021; Weziak-Bialowolska *et al*., 2022; Dimonte *et al*., 2023; Lanooij *et al*., 2023). Using a range of male mice that differ in the extent of their mating, siring, and caregiving experience, we were able to disentangle the influence of these experiences on indicators of recognition memory and anxiety-like behavior. We predicted that the males that interacted with pups (i.e., pup-sensitized non-fathers, first-time fathers, and experienced fathers) would exhibit improved memory compared to virgins and non-fathers because previous work showed that, beginning on PND 12, first-time California mouse fathers demonstrated enhanced spatial memory on a dry land maze test compared to virgins and pup-exposed virgins (Franssen *et al*., 2011). However, Glasper and colleagues (2011) observed that fathers did not exhibit improved recognition memory compared to non-fathers when tested at pup weaning, therefore paternal experience may differentially affect varying aspects of learning and memory and may vary with the amount of interaction with pups.

Here, we demonstrate that during a time of peak father-offspring interaction (i.e., PND 19), multiparous fathers display increased recognition memory compared to virgins but not non-fathers. In addition, we predicted that males that interacted with pups would exhibit decreased anxiety-like behavior compared to virgins and non-fathers, and indeed, we observed that experienced fathers displayed lower levels of anxiety-like behavior compared to virgins and non-fathers. This finding is consistent with previous studies that reported that on PND 16 and 19, California mouse fathers exhibited decreased anxiety-like behavior on elevated plus-maze tests compared to non-fathers (Glasper *et al*., 2016; Hyer *et al*., 2016), as well as expectant fathers and isolated virgins on PND 3 and 4 (Chauke *et al*., 2012). Together, these data provide additional evidence that pup-interaction is associated with an improved emotional state in this species. These behavioral changes in California mouse fathers may be particularly important in the context of caregiving since it is necessary for rodent fathers to efficiently recognize and locate their mate and offspring while defending their nest from intruders (McCabe & Blanchard, 1950; Cum *et al*., 2024). Moreover, it is at PND 19 when California mouse pups stray from the nest and fathers must locate and retrieve them (Bester-Meredith *et al*., 1999), therefore, it has been proposed that an attenuation of anxiety-like behaviors may be required for males to perform pup retrieval (Hyer *et al*., 2016). Unlike in California mouse fathers, findings in humans are mixed. The transition to fatherhood has been associated with increased stress, anxiety, and depression (Fletcher *et al*., 2006; Singley & Edwards, 2015), whereas, in other studies, fathers have a decreased likelihood of diagnosed depression (Pulkki-Råback *et al*., 2012), improved overall psychological well-being (Goldman, 2012), and report better job performance when they interact more with their children (Ladge *et al*., 2015). Furthermore, a study of another biparental rodent, the prairie vole, demonstrates that fatherhood increases anxiety- and depressive-like behaviors, and reduces social recognition memory on PND 6-10 (Lieberwirth *et al*., 2013). Taken together, these data highlight the complex relationships between paternal care and behavioral outcomes, underscoring the need to better understand broad patterns of paternal experience-induced behavioral changes.

Interestingly, in Experiment 2, when we compared memory and anxiety-like behaviors in non-fathers, pup-sensitized non-fathers, and first-time fathers, we observed no differences among groups. This suggests that more time caring for pups may be necessary to observe changes in recognition memory in California mouse males, since experienced fathers that had cared for two litters exhibited increased memory in Experiment 1. This finding is further supported by a study in maternal rats in which multiparous females exhibit improved spatial memory in a dry land maze compared to nulliparous females (Love *et al*., 2005). However, all Experiment 2 males, including non-fathers, had similarly low levels of anxiety-like behavior which were comparable to those of experienced fathers in Experiment 1. We cannot exclude the possibility that this finding resulted from increased handling of the mice (Walf and Frye, 2007; Sensini *et al*., 2020) during testing since pup-sensitized non-fathers and first-time fathers were handled during the pup-exposure and caregiving assays. We also handled the non-fathers in Experiment 2 equally to control for this effect, but all males from Experiment 1 were handled less; therefore, future studies might benefit from minimal animal handling when anxiety-like behaviors are being assessed.

In this study, we tested the hypothesis that caregiving experience in the absence of reproduction (i.e., pup-sensitized non-fathers) would improve memory and decrease anxiety-like behaviors, as seen in fathers (Franssen *et al*., 2011; Glasper *et al*., 2016; Hyer *et al*., 2016). While we did not find evidence to support this prediction, we did observe that when non-fathers interact with pups, their caregiving behaviors differ from biological fathers. During the caregiving assay, pup-sensitized non-fathers were initially slower to approach pups, but eventually spent more time grooming pups, while first-time fathers spent more time nest building. This result is further highlighted in the principal component analysis in which grooming and huddling behaviors are colinear in the opposite direction of nest building and sniffing in the PCA space and the pup-sensitized non-father group mean is closest to grooming/huddling, whereas the first-time father group mean is closest to nest building/sniffing (Figure 4C). Additionally, observations from this study at PNDs 14 and 16, the time when pups begin to first leave the nest, suggest that pup-sensitized non-fathers prioritized grooming and huddling pups, whereas first-time fathers repeatedly checked on their pups by sniffing while exploring the cage and building a nest. This observed pattern may signal different ultimate explanations responsible for these behaviors, for example, nest building in fathers may be necessary for early pup survival (Lewarch & Hoekstra, 2018), whereas grooming in pup-sensitized non-fathers may be critical for the maintenance of pup condition (Spruijt *et al*., 1992). Our results are consistent with previous findings in which *P. polionotus* mothers that had greater exposure to pups exhibited improved nest building behavior compared to inexperienced mothers (Margulis *et al*., 2005), and, similarly, in house mice and rats the amount of maternal experience influences the latency to care for pups (Cohen & Bridges, 1981; Koch & Ehret, 1989). Moreover, naïve male rodents can be sensitized to behave parentally, including house mice (Ehret *et al*., 1986), rats (Rosenblatt, 1967), and golden hamsters (Swanson & Campbell, 1979). These species, however, do not exhibit caregiving behaviors in the wild; therefore, their paternal behaviors and those of wild biparental species may not always be comparable. Contrary to our findings, however, Horrell and colleagues reported that virgin California mouse males that had been paired with another male and sensitized to pups over five days displayed the same paternal behaviors when compared to first-time fathers (Horrell *et al*., 2017). The caregiving behaviors we observed in pup-sensitized non-father “babysitters” and first-time fathers also align with caregiving dynamics seen in various other species, such as prairie voles in which paternal and alloparental care by siblings is common (Thomas & Birney, 1979), several primate and canid species in which females often act as babysitters, as well as alloparental avian communities that rely on “helpers at the nest” (reviewed in Riedman, 1982 and Mocha *et al*., 2023), suggesting a broader context for understanding these behaviors. Importantly, our results provide evidence that California mouse males will engage with unrelated pups and provide care, even if they have never reproduced themselves, yet the caregiving behaviors they display differ from those of fathers.

Parental care varies in intensity between species and can be energetically costly (Gubernick & Klopfer, 1981; Clutton-Brock, 1991). Similarly, sperm production is also costly (Hayward & Gillooly, 2011), and thus, males can experience a tradeoff in reproductive investment between ensuring offspring survival through parental care and maximizing reproductive potential via sperm production (Stearns, 1989; Stiver & Alonzo, 2009; Lynch, 2016; Requena & Alonzo, 2017). To examine if there is a tradeoff in these two forms of reproductive investment, we assessed testes size and sperm count in virgins, non-fathers, and experienced fathers and found limited evidence to support this prediction. We found that virgins had smaller testes and less sperm than paired non-fathers which is likely associated with lack of access to females and mating experience (Drori & Folman, 1967). Moreover, although not statistically significant, we found that experienced fathers tend to have smaller testes and less sperm than paired non-fathers, potentially suggesting that when fathers have pups, they reduce reproductive investment and shift resources toward providing care. Patterns observed in earlier studies on California mice demonstrated that non-fathers and first-time fathers have larger testes than virgins (Harris and Saltzman, 2013; Zhao *et al*., 2017), which is a robust indicator of sperm production (Parker and Pizzari, 2010). Interestingly, in each of these studies, and including the results we present here, virgins have the smallest testes, followed by fathers and non-fathers paired with sterile females, but non-father and father testes sizes were not significantly different. Yet, when we compared a second group of non-fathers with first-time fathers in Experiment 2, we found that first-time fathers have larger testes than non-fathers, which is counter to the prediction of a tradeoff between reproductive potential and offspring survival. This result could suggest that more time with pups is necessary to see the reduction in testes size, as evidenced in the experienced fathers. Overall, our results suggest that California mouse males increase reproductive potential when there is an opportunity to mate, but that caring for multiple litters may eventually result in reduced reproductive investment in biparental monogamous species like the California mouse.

In summary, our findings shed new light on the complex effects that mating, exposure to pups, siring offspring, and caregiving experience have on several behavioral and reproductive phenotypes in the male California mouse. We demonstrate that mating experience, fatherhood, and experience caring for pups are associated with changes in memory, anxiety-like behavior, reproductive investment, and the type of caregiving behaviors performed, and that even when the offspring are not their own, pup-sensitized California mouse males also exhibit caregiving. Therefore, these results not only offer valuable insights into the behavioral changes shaped by life experiences, but also underscore the relationship between parental behavior and sperm production in a monogamous biparental species.

## Author Contributions

M.E.C., H.S.F., and E.R.G. conceived of the study, designed experiments, and interpreted results; M.E.C. and P.A. collected the data, M.E.C. and D.K. carried out the statistical analyses; M.E.C. and H.S.F. wrote the manuscript, M.E.C., H.S.F., and E.R.G. edited the manuscript; all authors gave final approval for publication and agree to be held accountable for the work presented.

## Data Availability

The raw data files supporting the findings of this study and the scripts used for data analysis are available on Dryad at https://datadryad.org/stash/share/aTzBpTPekrnQhkhi4u3FB5mlbSA0dSJzPkvXfvdFxPA.

