## Supplement for "Reproductive experience drives changes in behavior and physiology in male California mice (*Peromyscus californicus*)"

**Supplemental Figures**

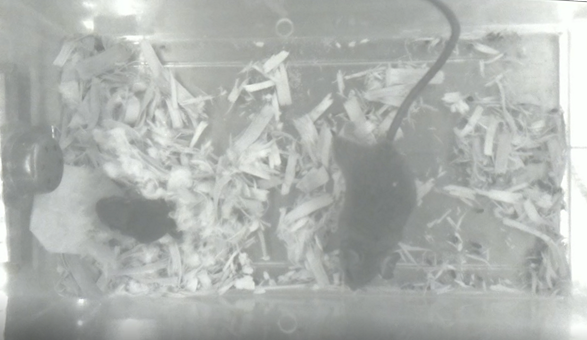

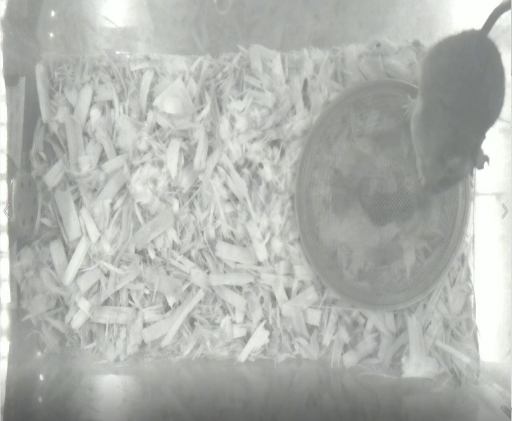

**Pup outside mesh enclosure**

**Pup within mesh enclosure**

**A**

**B**

**Figure S1.** Image of male during **(A)** pup-exposure assay and **(B)** caregiving assay.

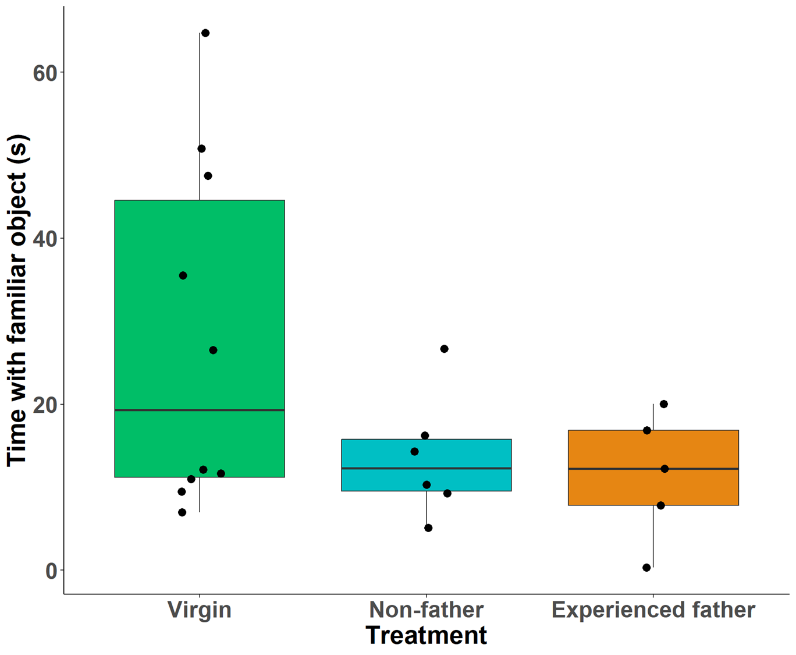

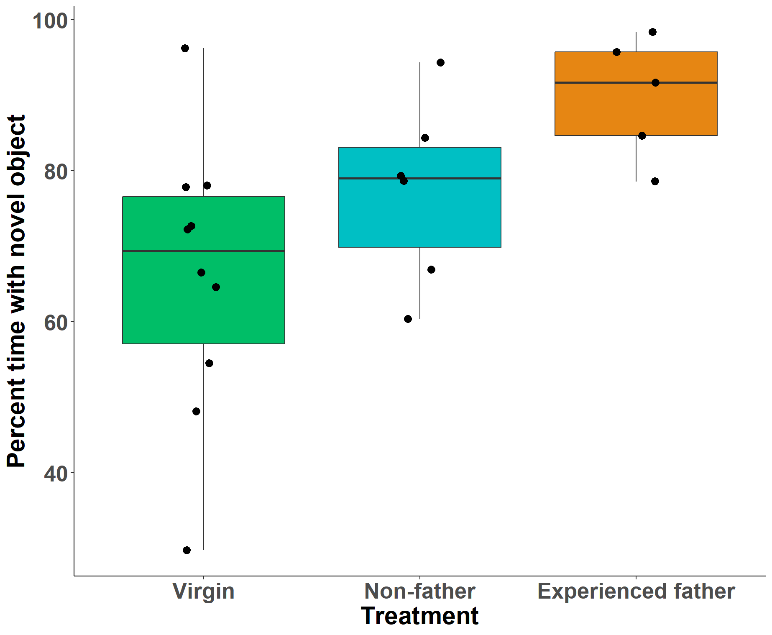

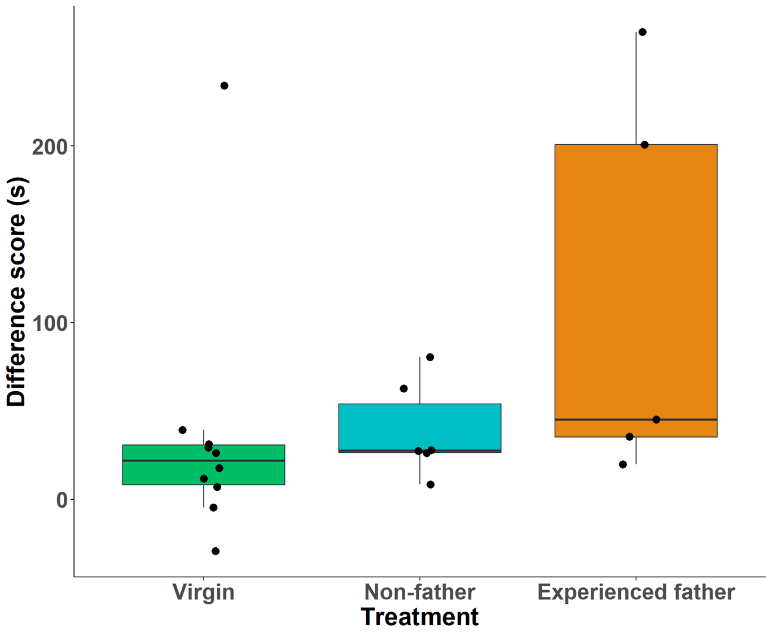

*P* = 0.01

**D**

**B**

**C**

**A**

*P* = 0.00896

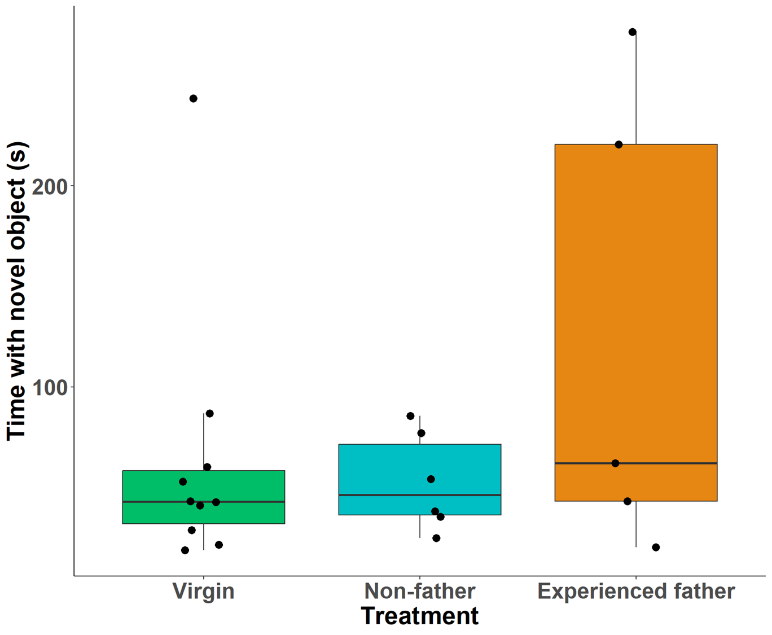
**Figure S2.** Novel object recognition **(A)** difference score, **(B)** percent time exploring novel object, and exploration of **(C)** novel and **(D)** familiar object in Experiment 1.

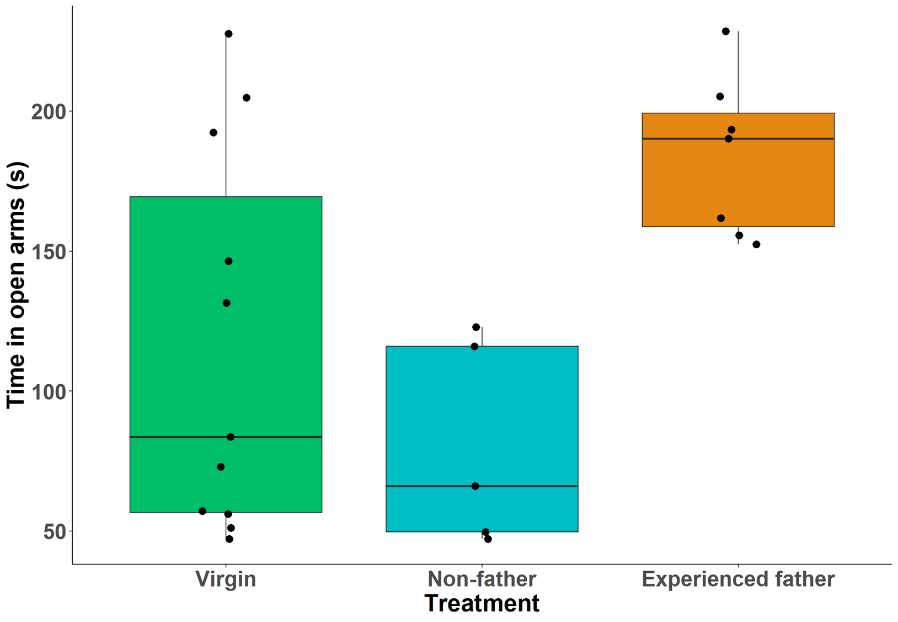

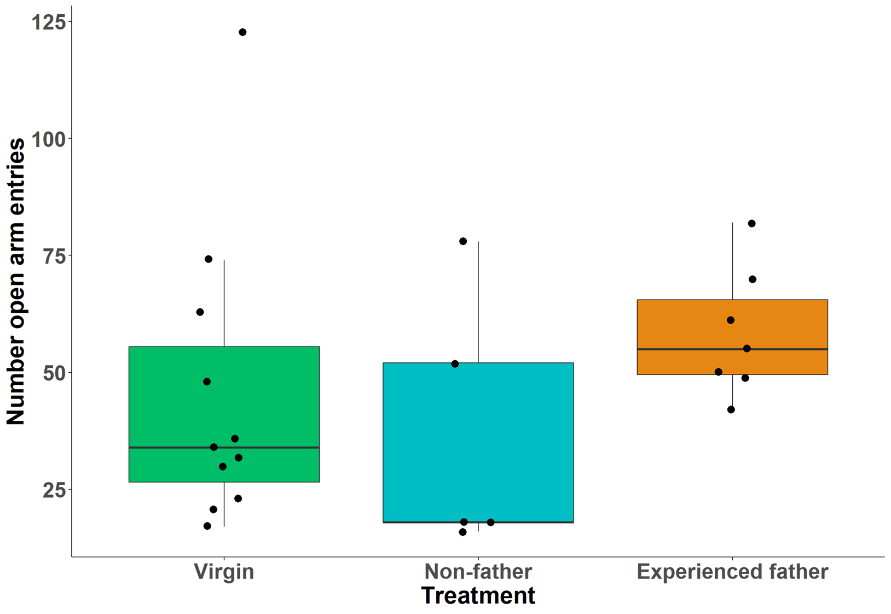

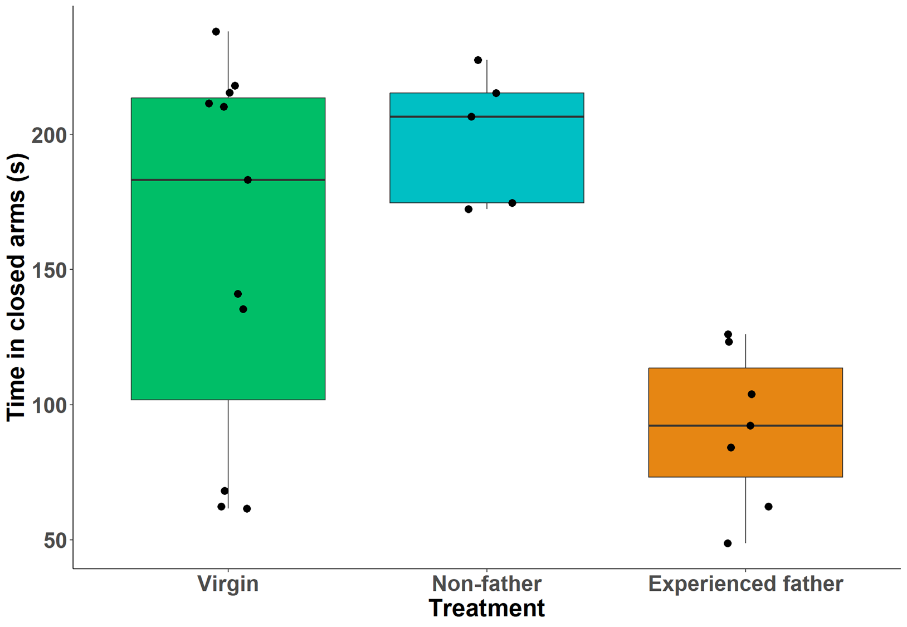

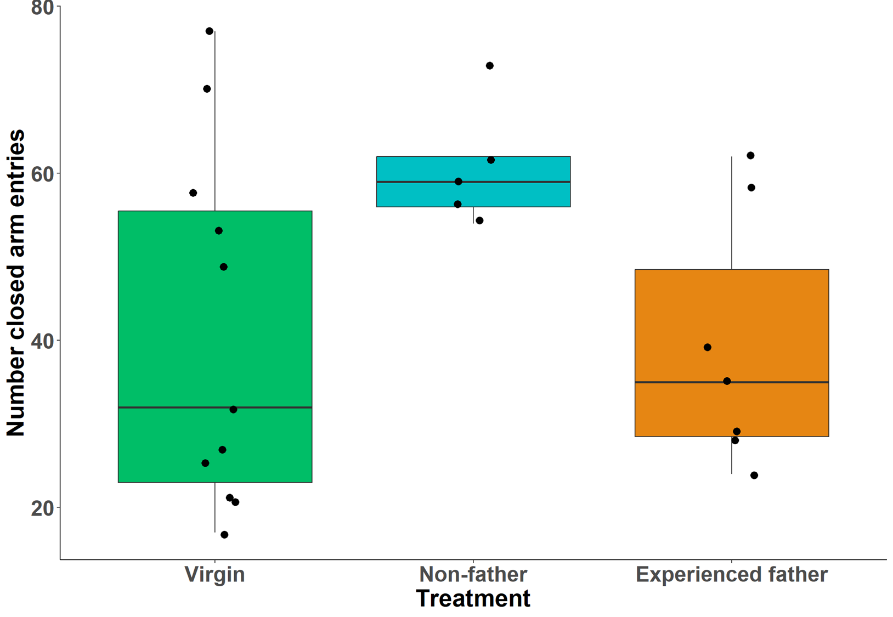
**Figure S3.** Elevated plus maze **(A)** number of closed arm entries and **(B)** open arm entries, and time in **(C)** closed arms and **(D)** open arms in Experiment 1.

*P* = 0.02

*P* = 0.002

*P* = 0.02

*P* = 0.003

**D**

**B**

**C**

**A**

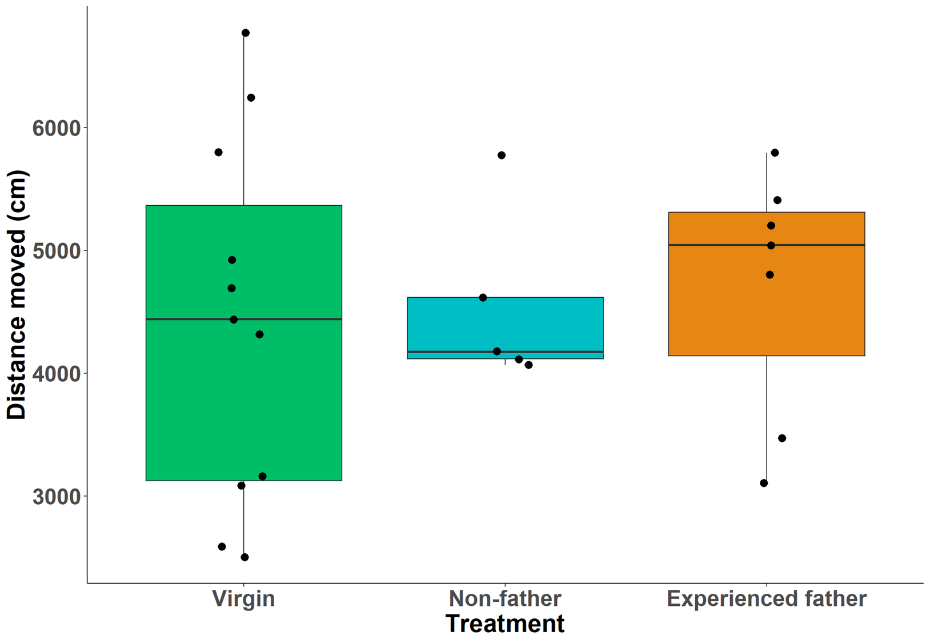
**Figure S4.** Distance moved on elevated plus maze in Experiment 1.

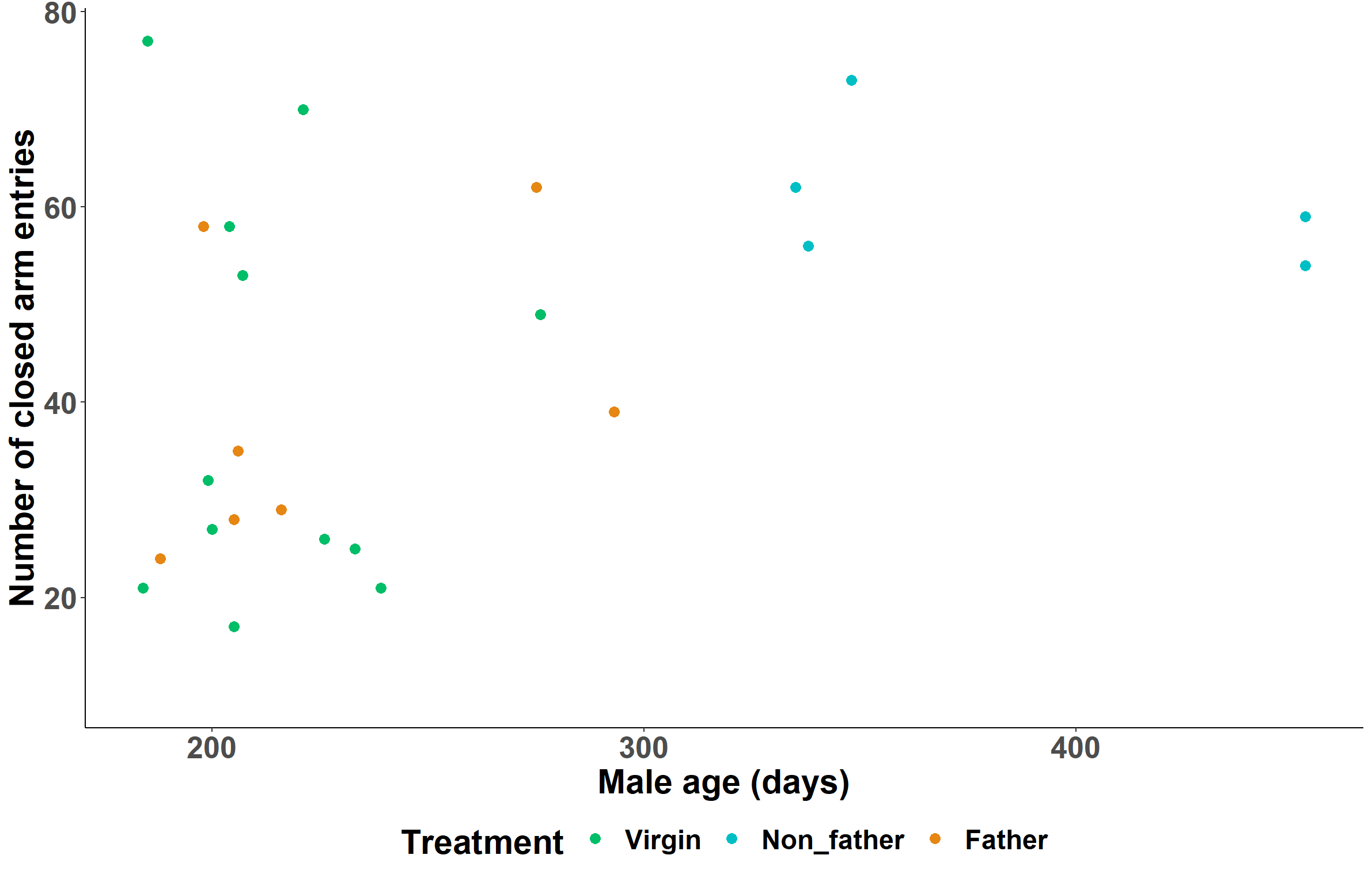

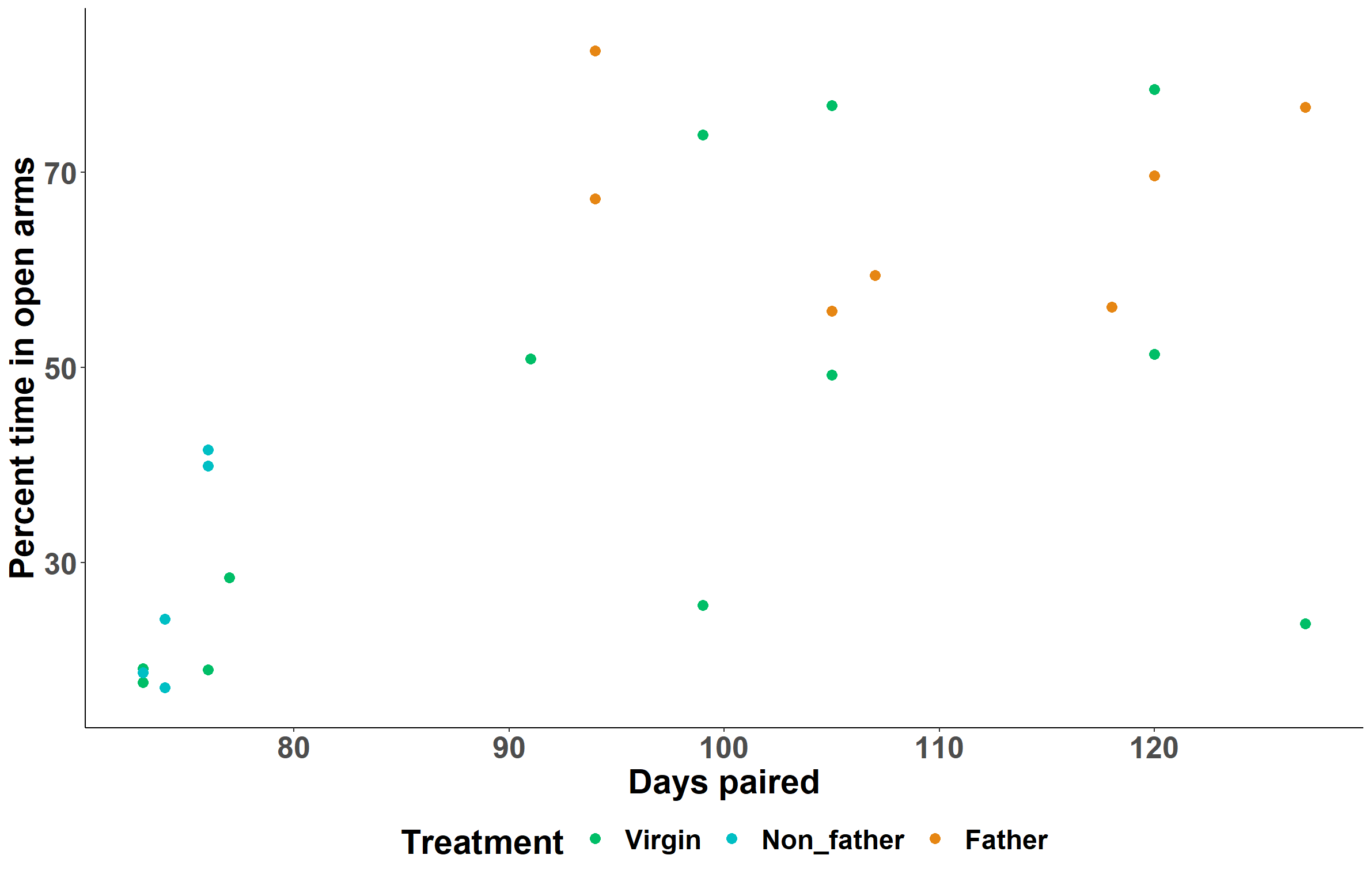

**Non-father**

**Experienced father**

**Virgin**

**Figure S5.** Effect of male age on the number of closed arm entries in Experiment 1.

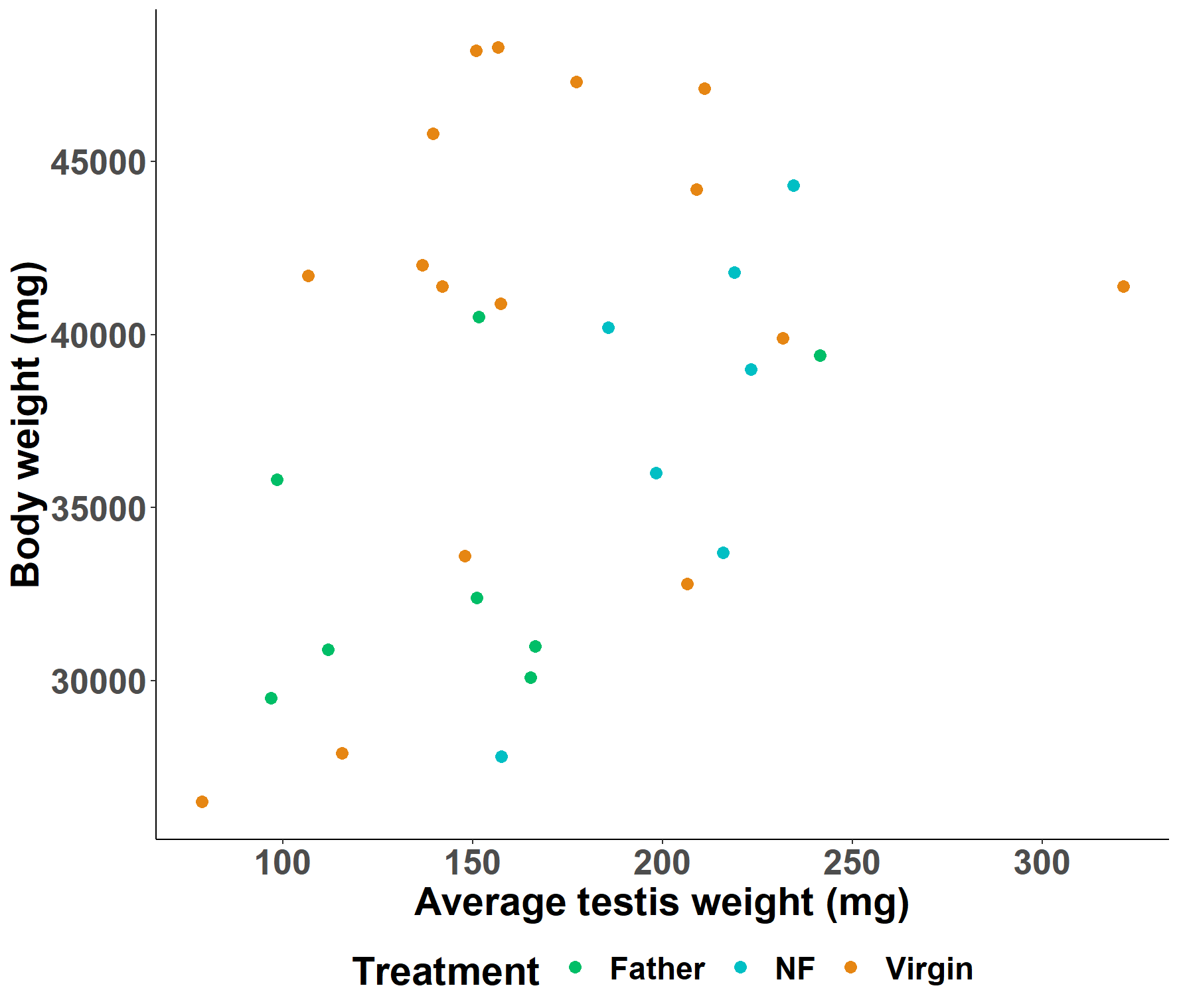

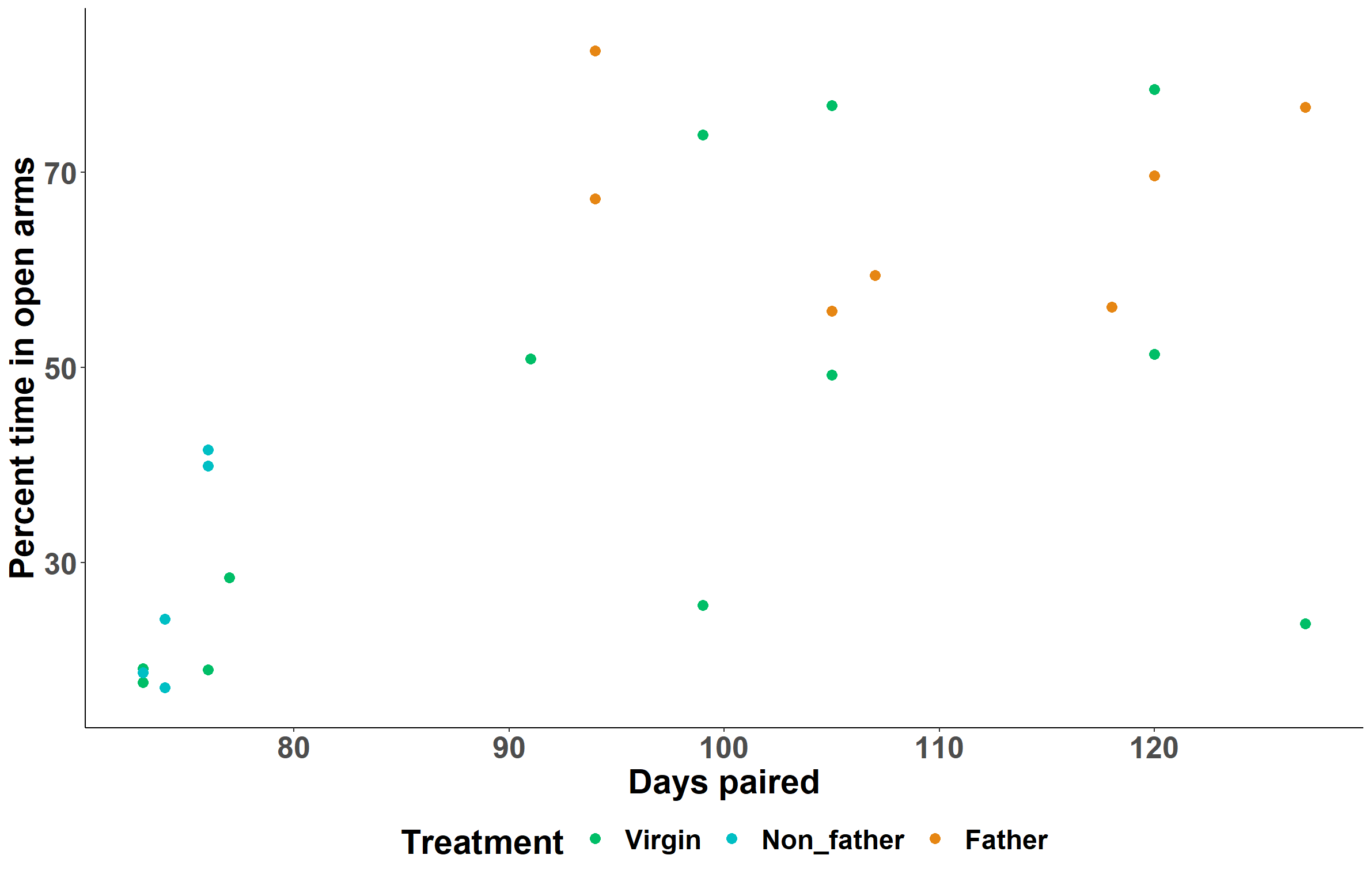

**Experienced father**

**Non-father**

**Virgin**

**Figure S6.** Effect of body weight on average testis weight in Experiment 1.

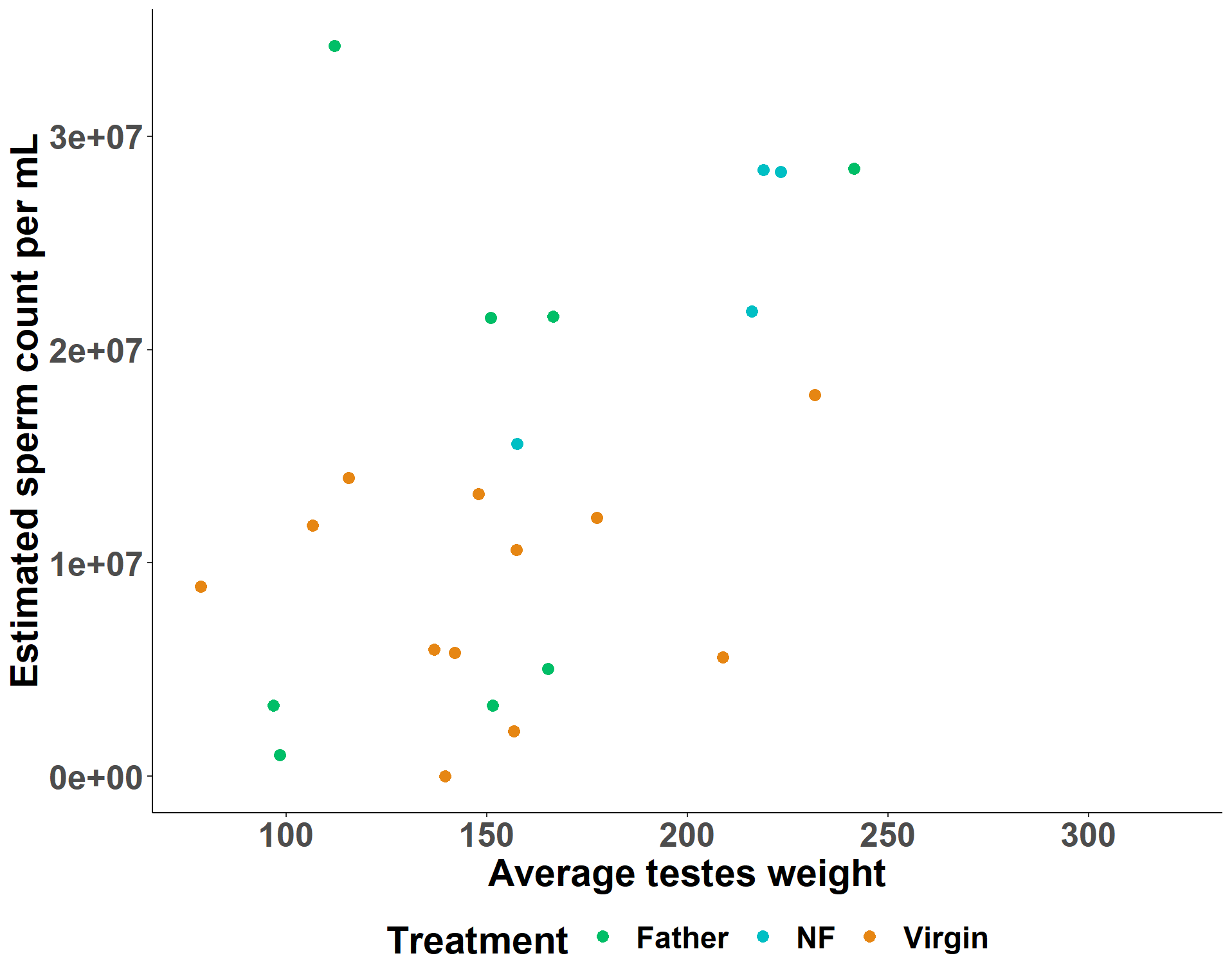

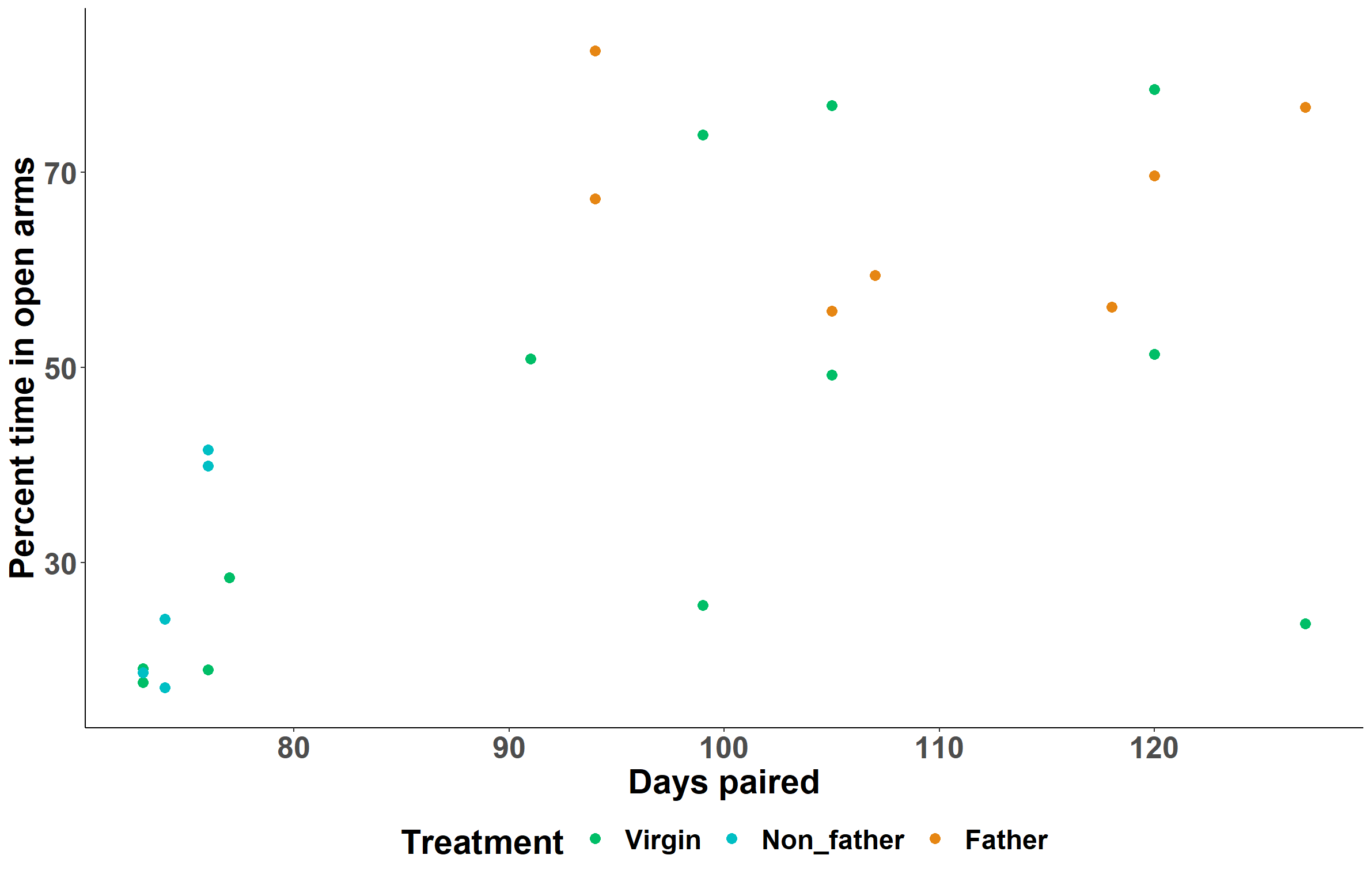

**Experienced father**

**Non-father**

**Virgin**

**Figure S7.** Effect of estimated sperm count on average testis weight in Experiment 1.

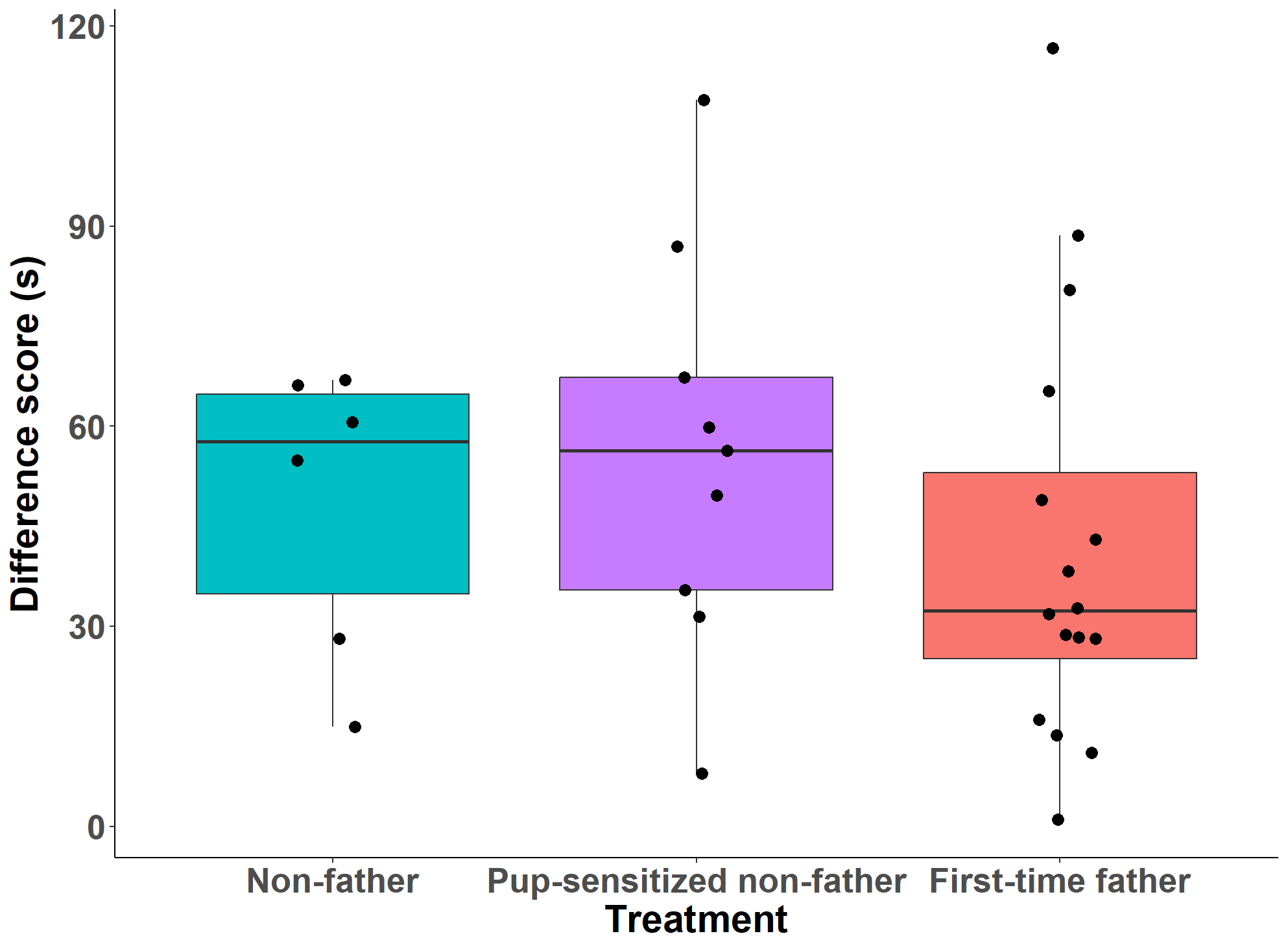

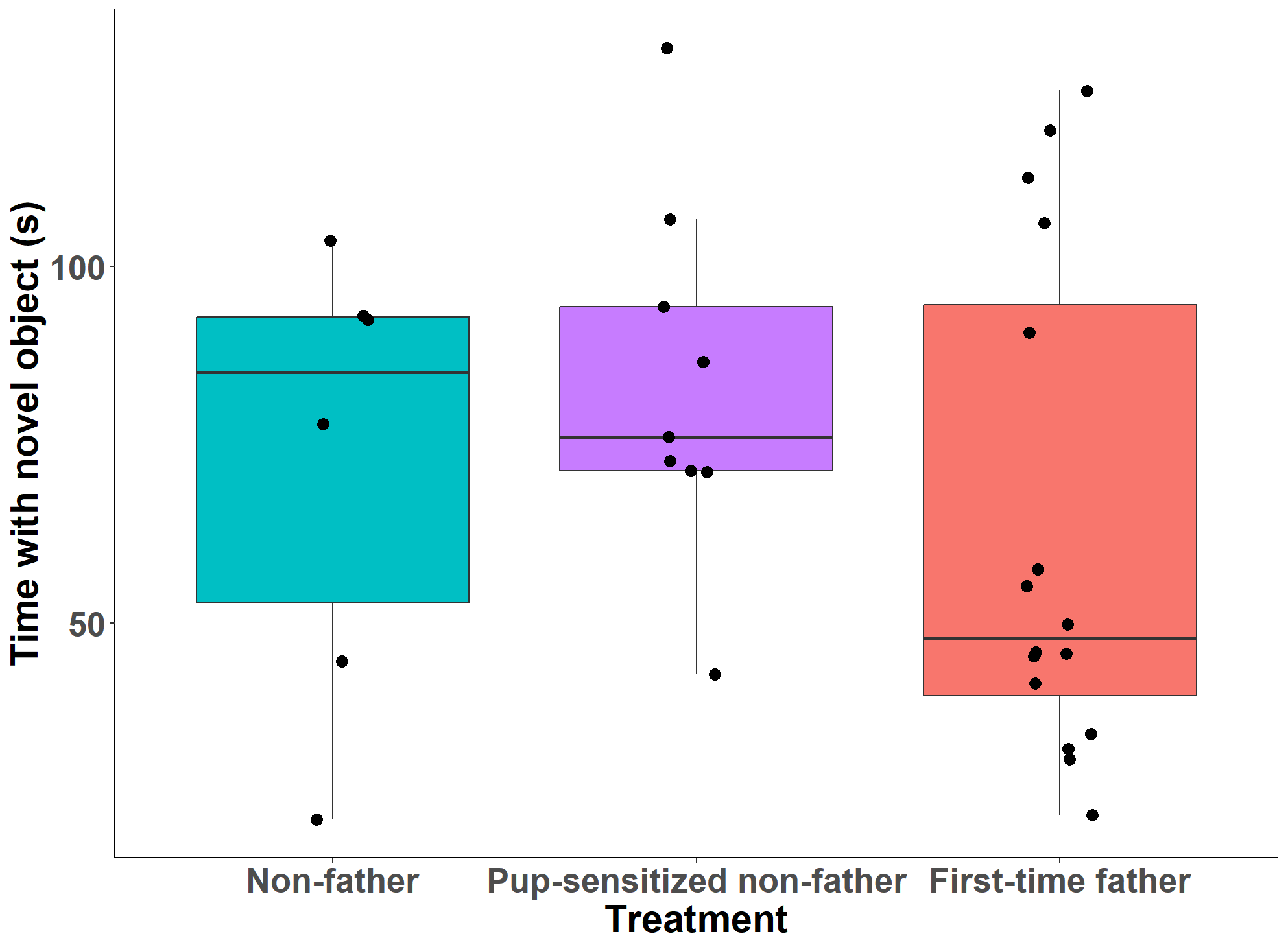

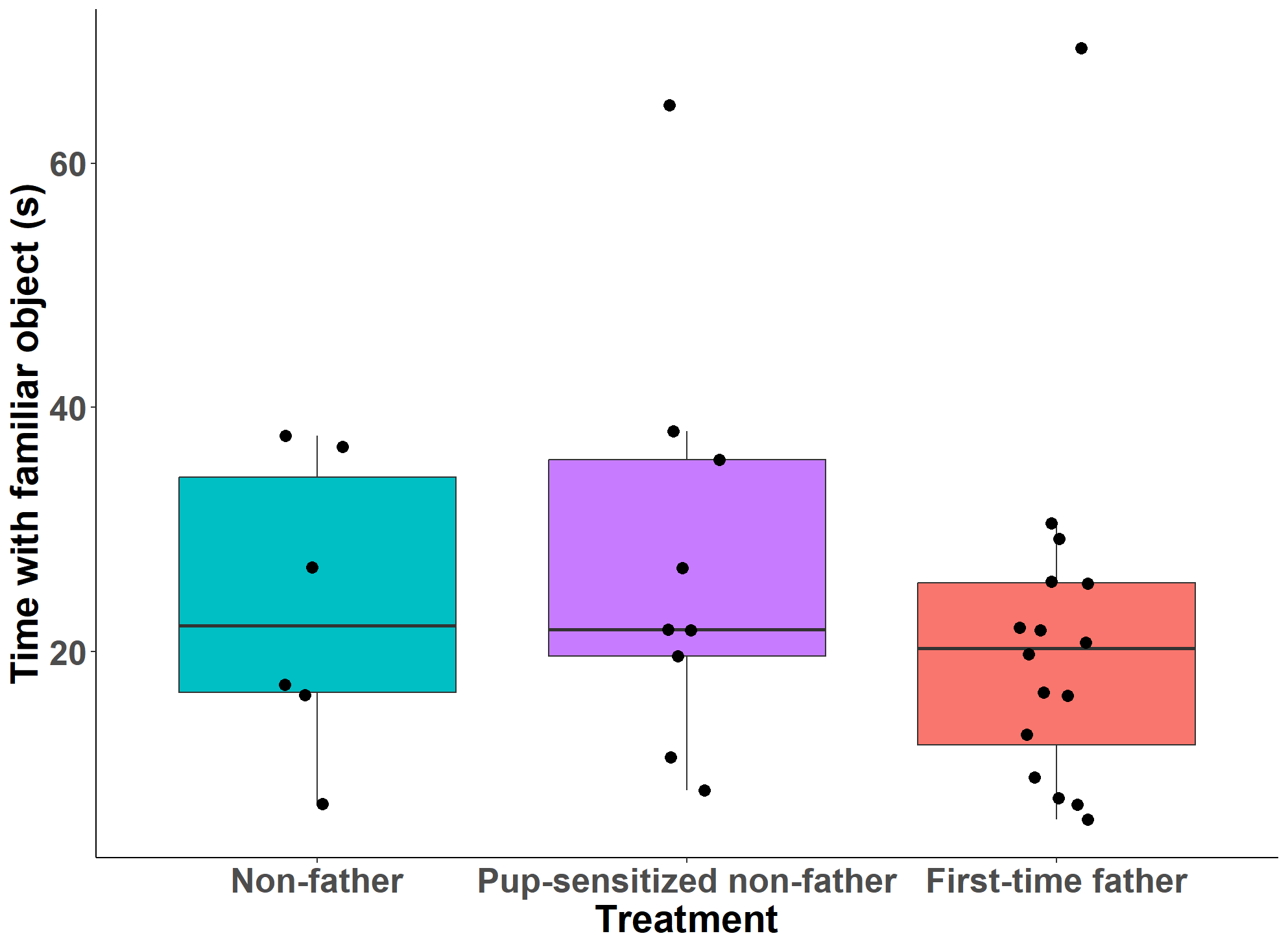

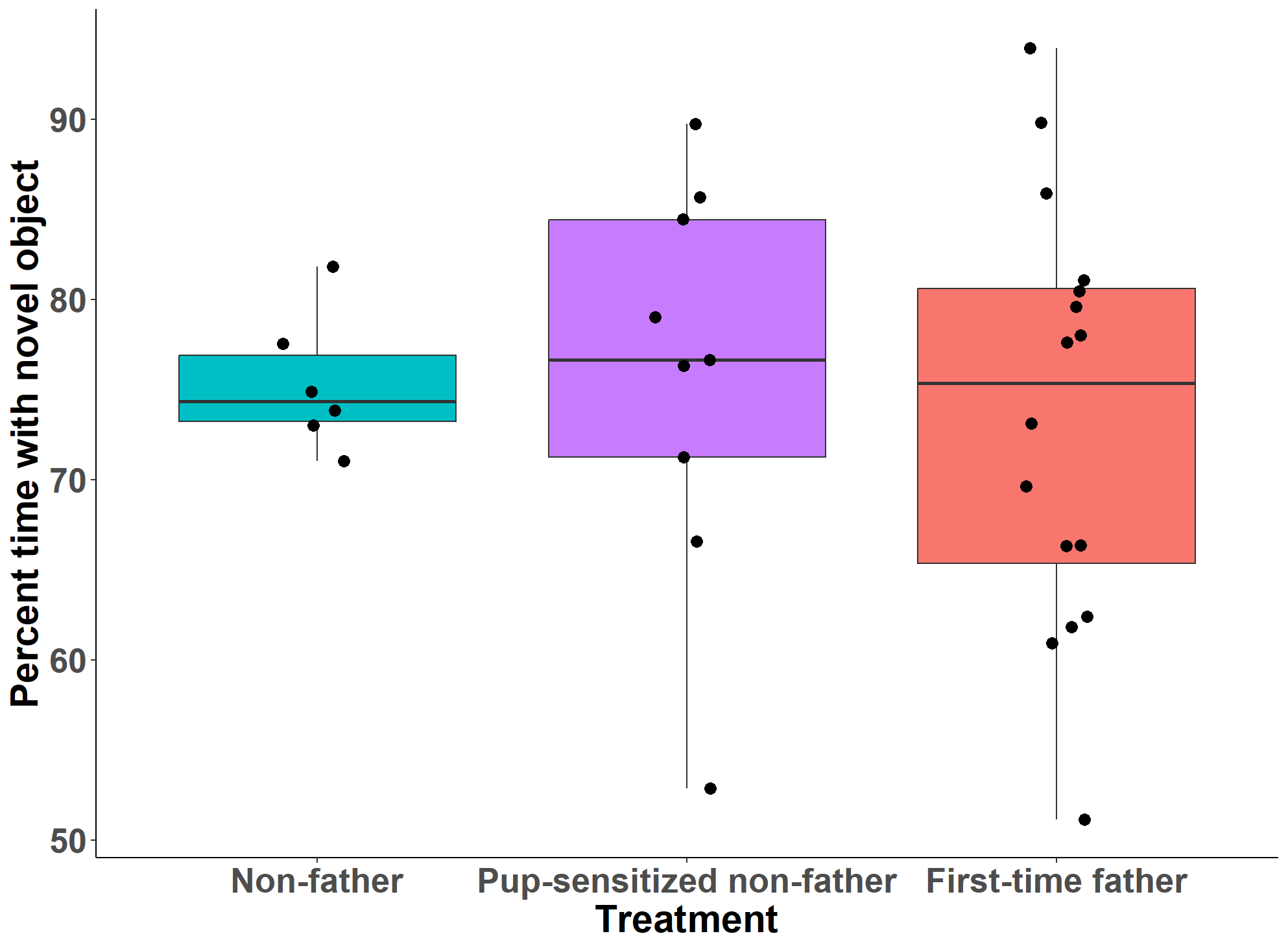
**Figure S8.** Novel object recognition test **(A)** difference score, **(B)** percent time exploring novel object, and exploration of **(C)** familiar and **(D)** novel objects in Experiment 2.

**B**

**D**

**C**

**A**

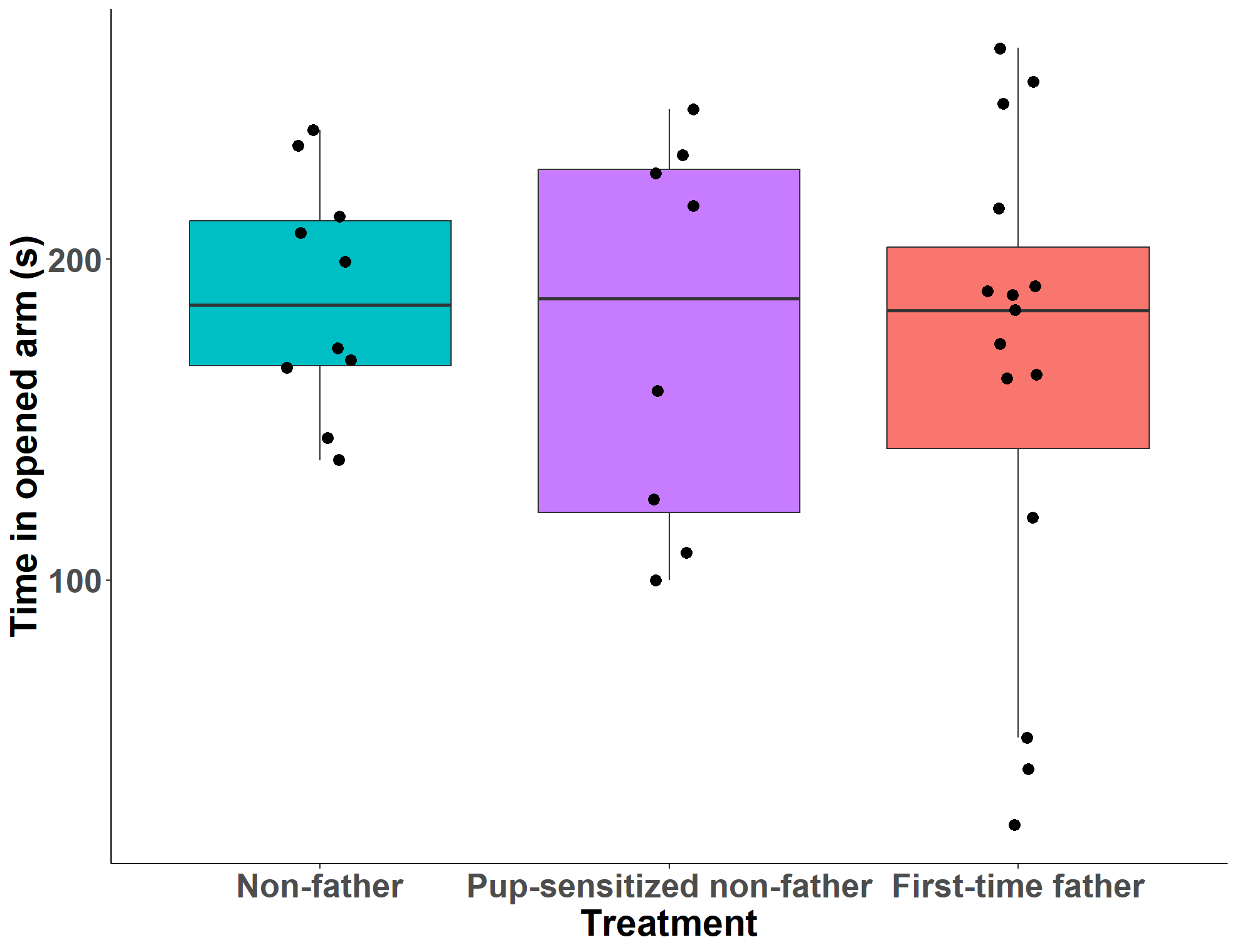

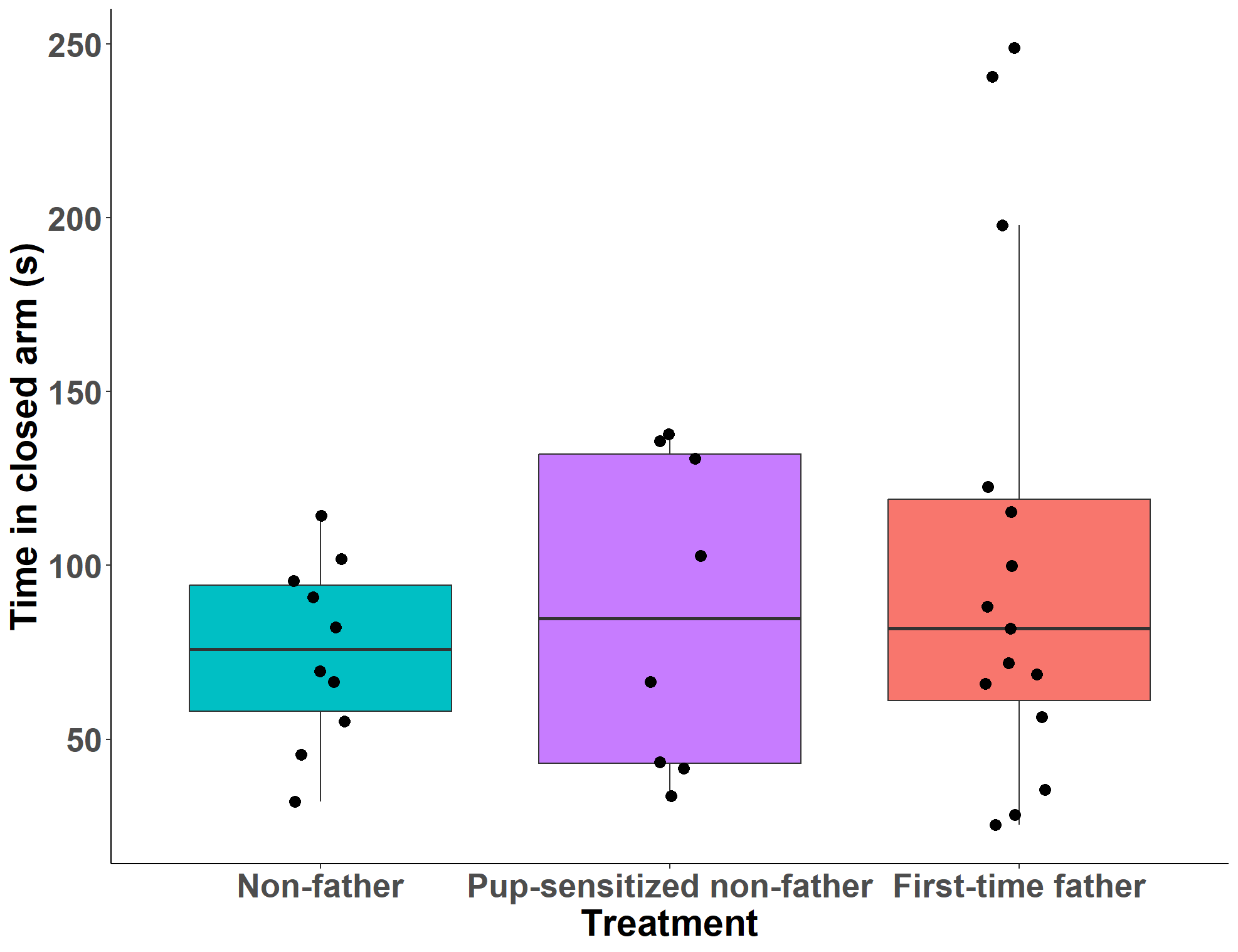

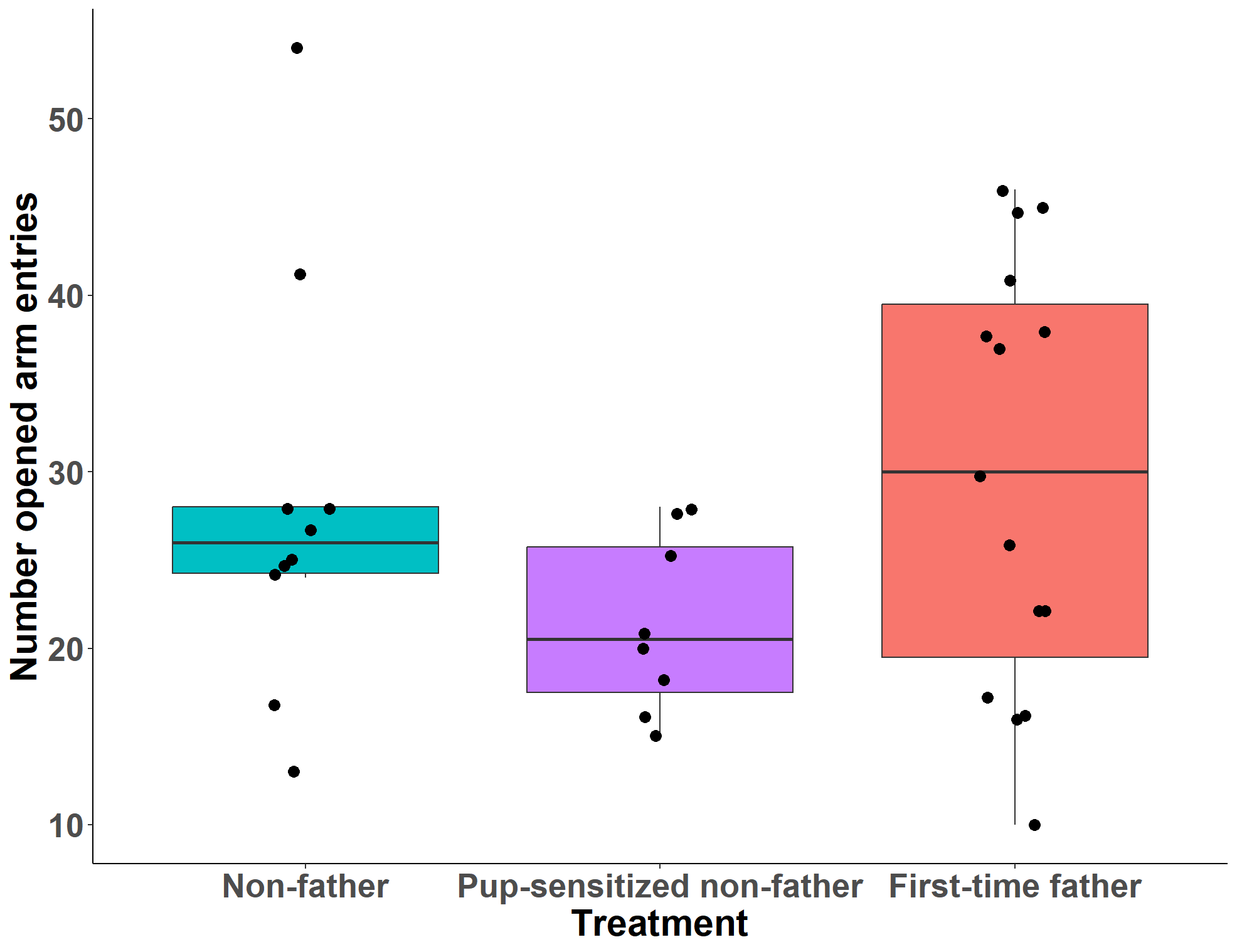

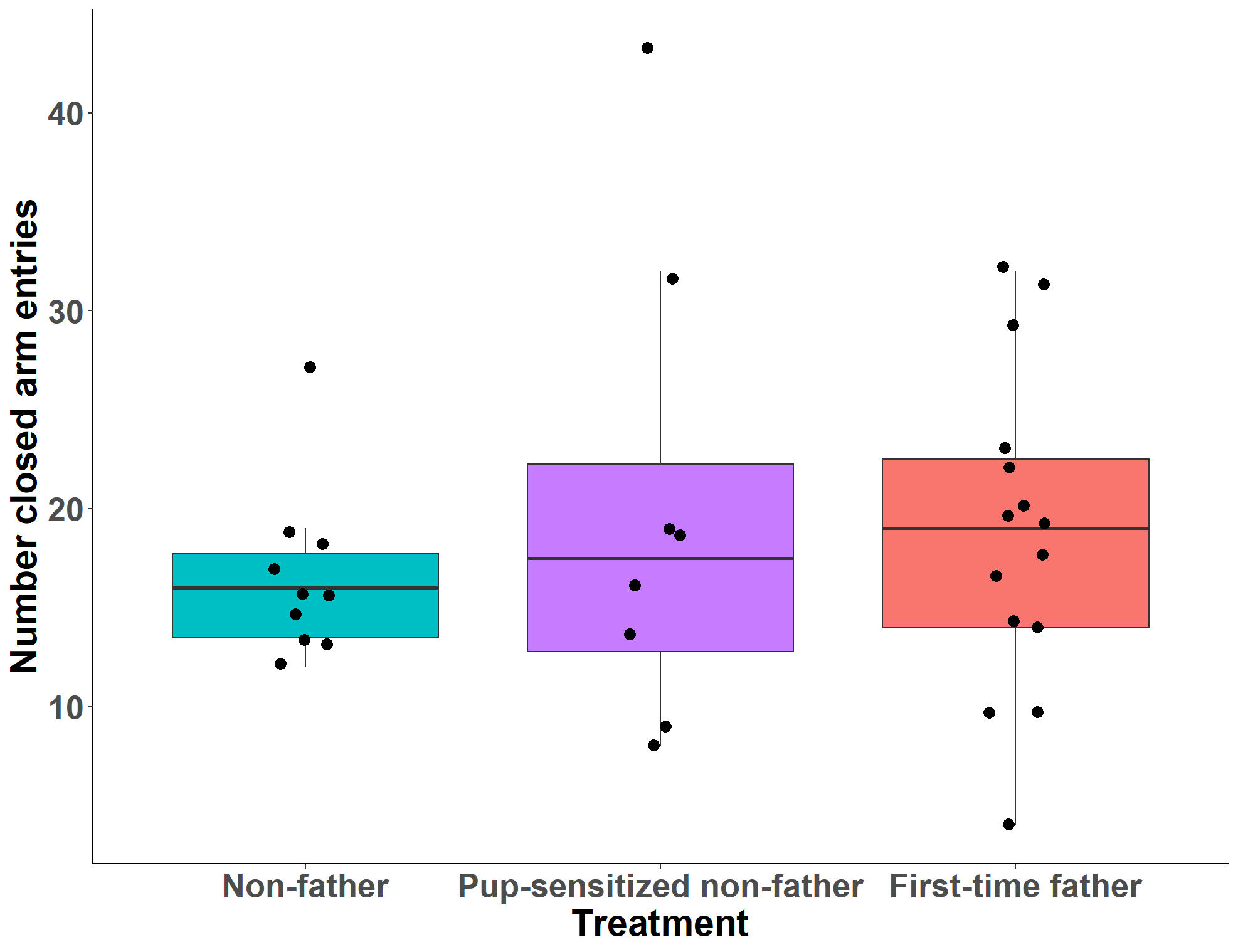
**Figure S9.** Elevated plus maze number and duration of (**A, B**) closed arm entries and (**C, D**) open arm entries in Experiment 2.

**D**

**C**

**B**

**A**

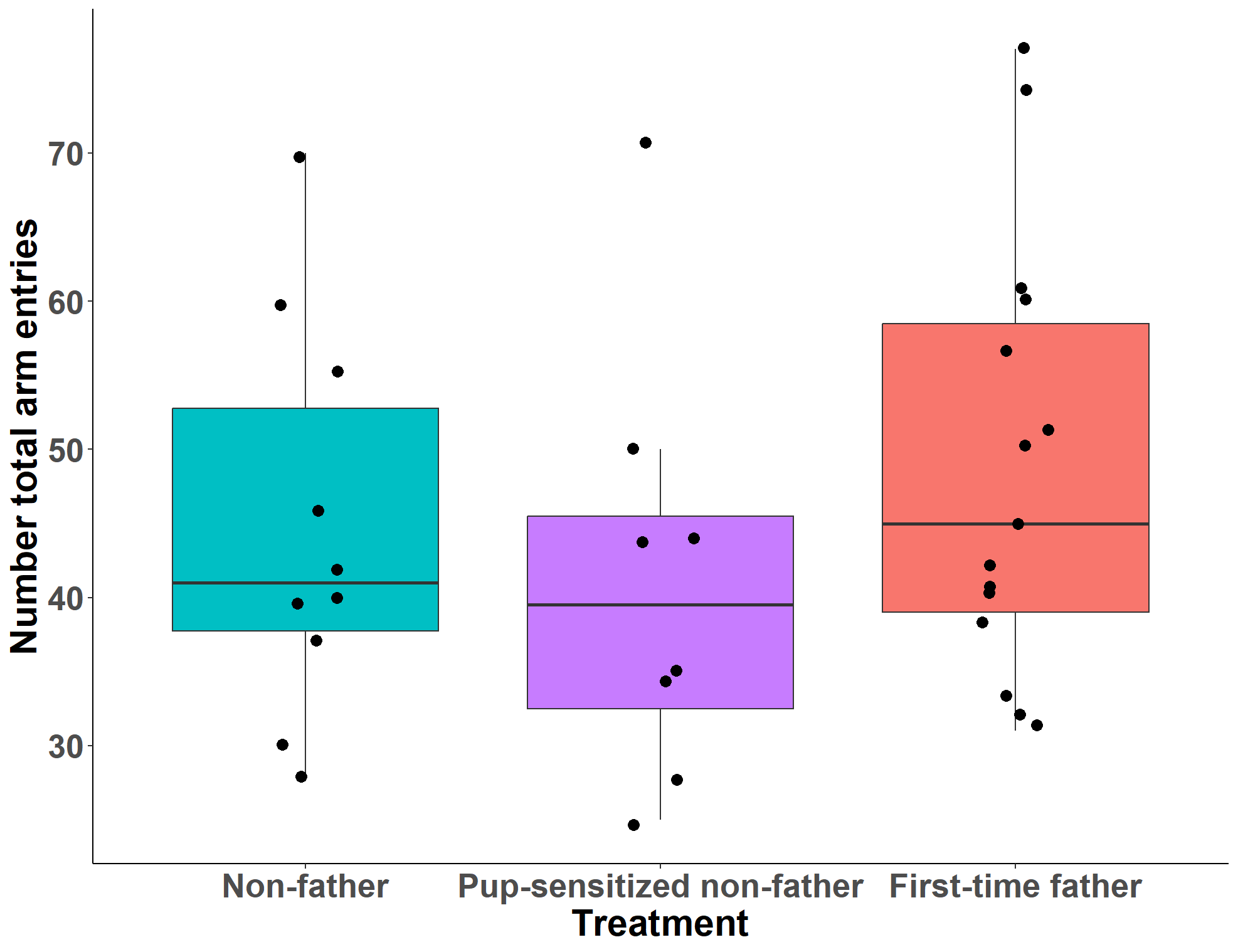

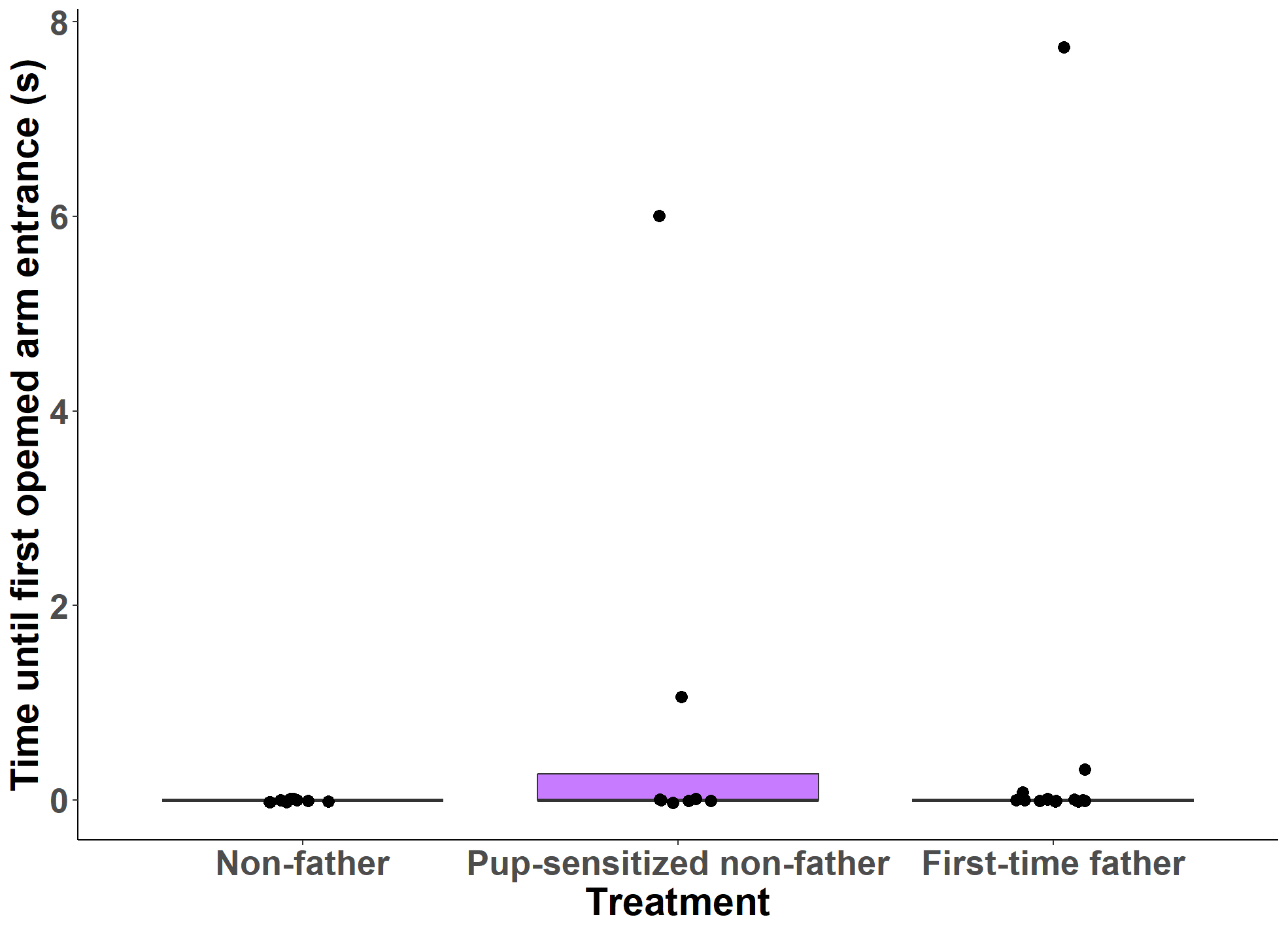

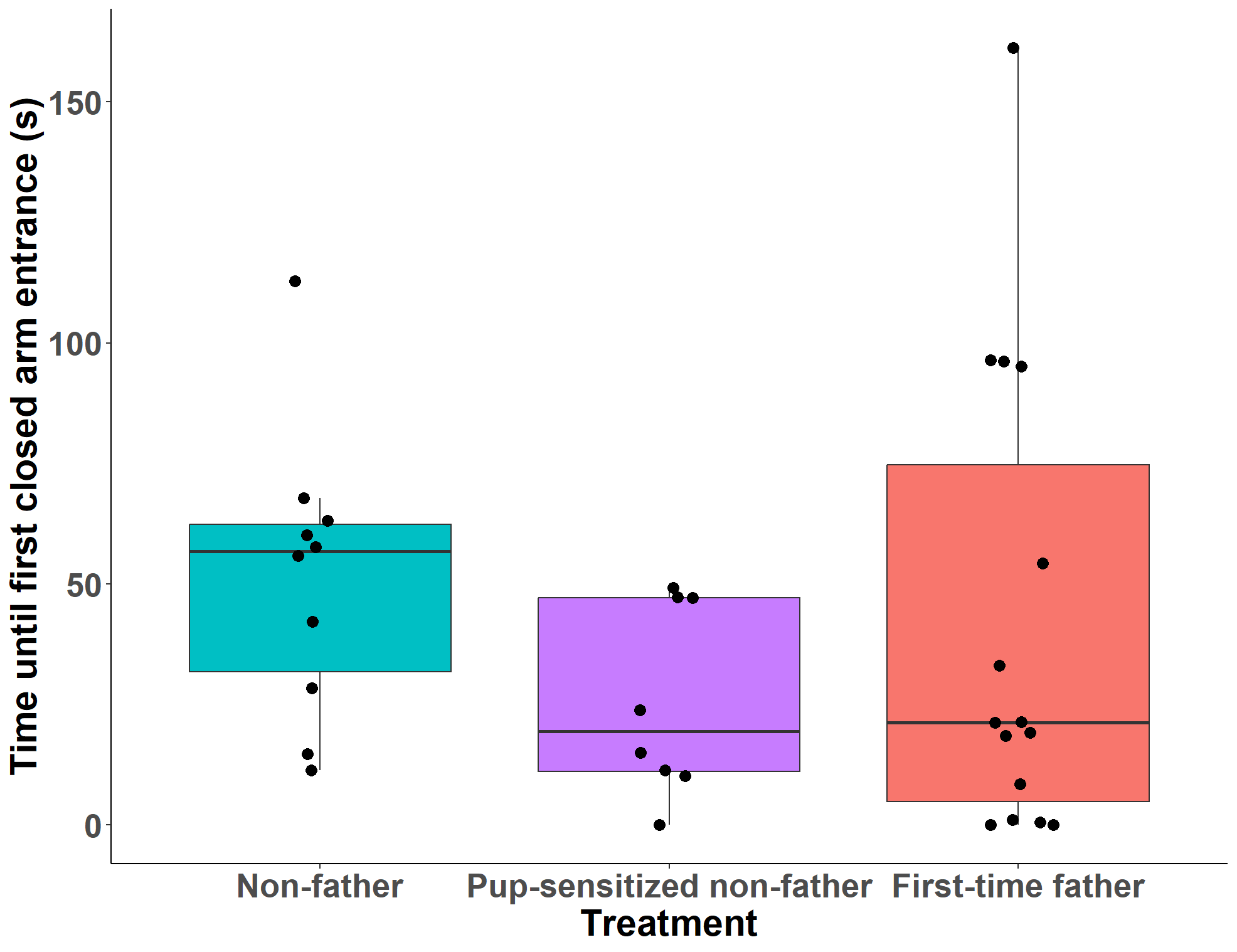
**Figure S10.** Elevated plus maze latency to enter **(A)** closed and **(B)** open arms and **(C)** total number of closed and open arm entries in Experiment 2.

**C**

**B**

**A**

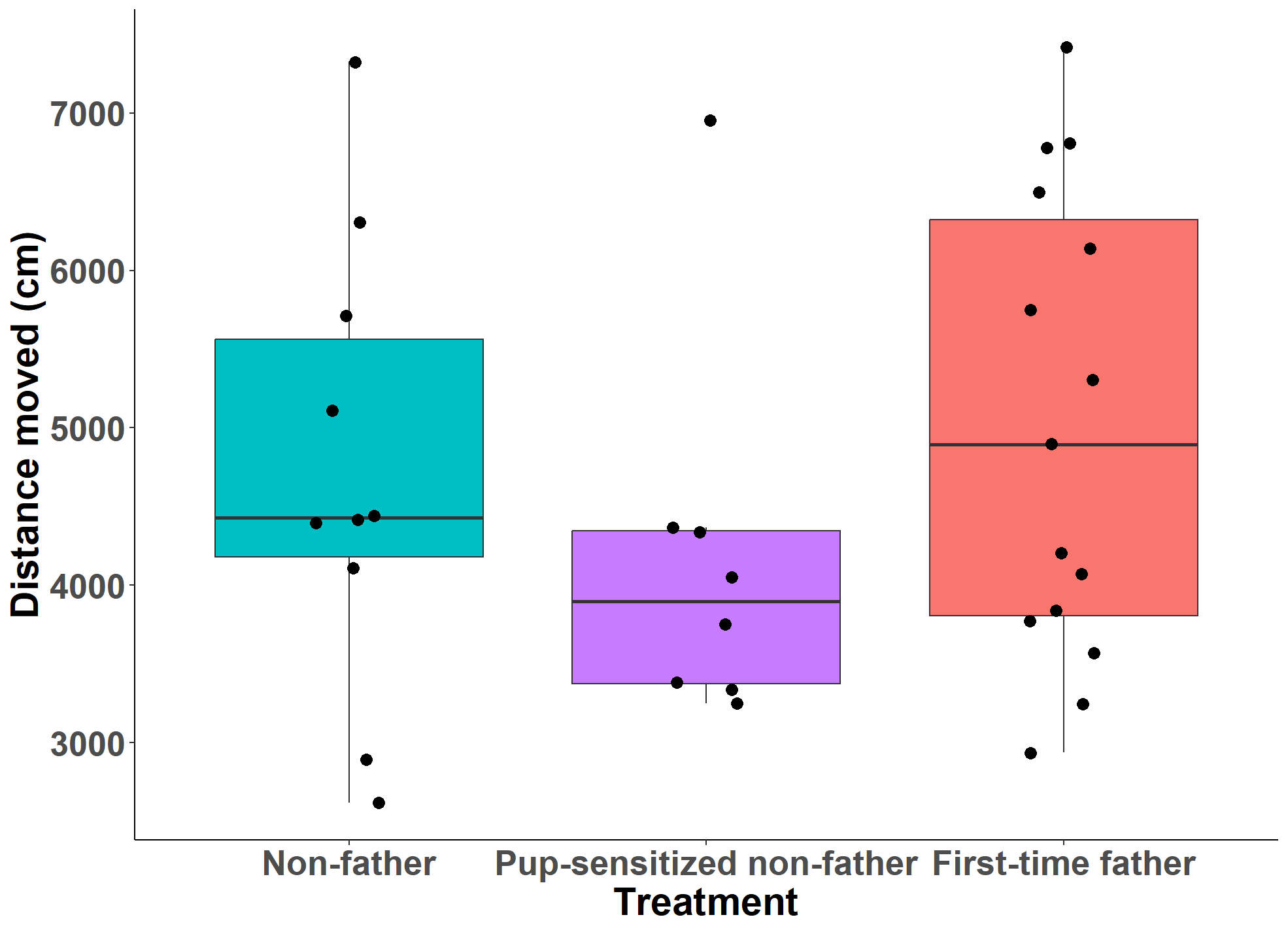

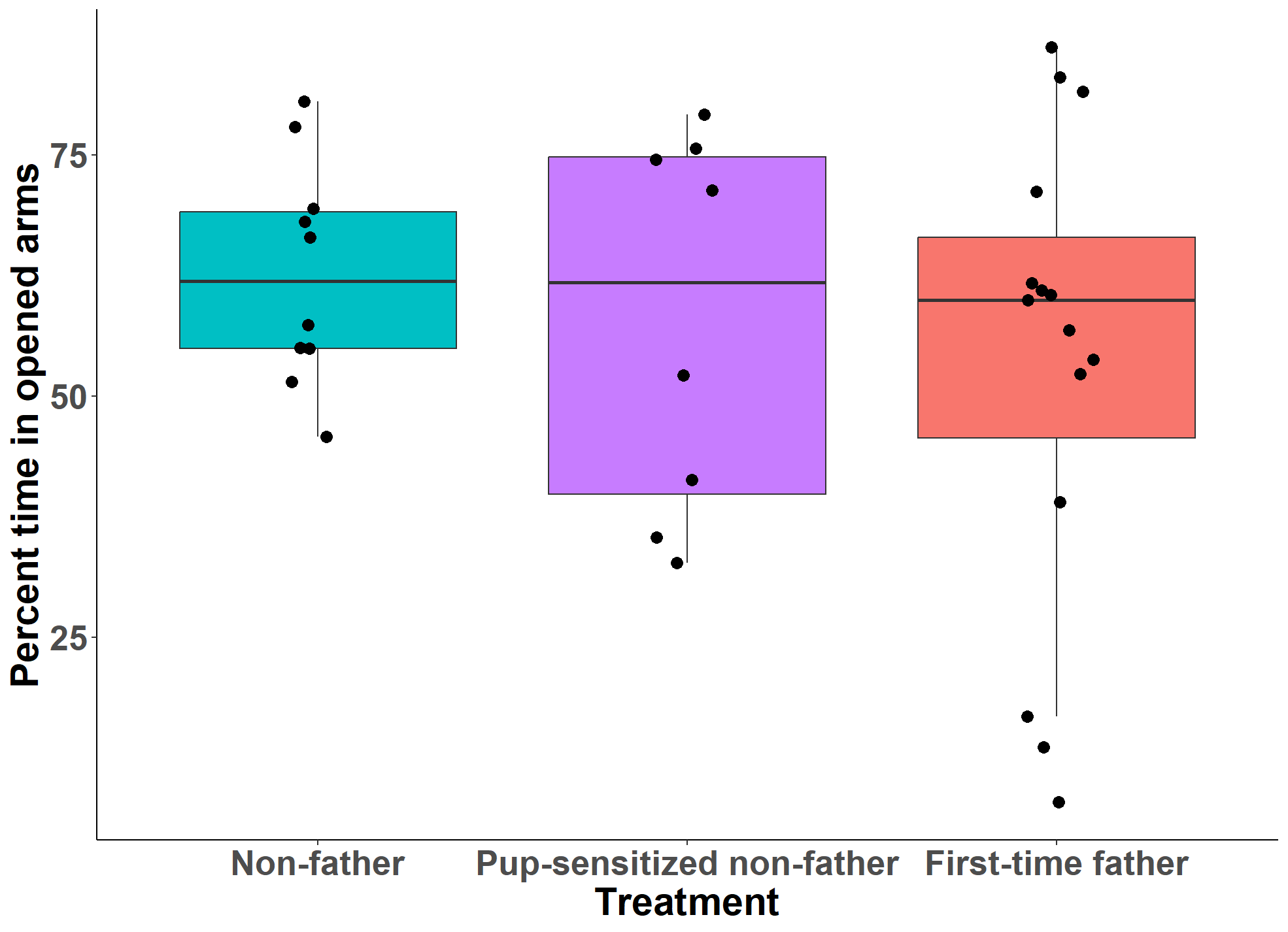
**Figure S11.** Elevated plus maze **(A)** percent time in open arms and **(B)** distance moved in Experiment 2.

**B**

**A**

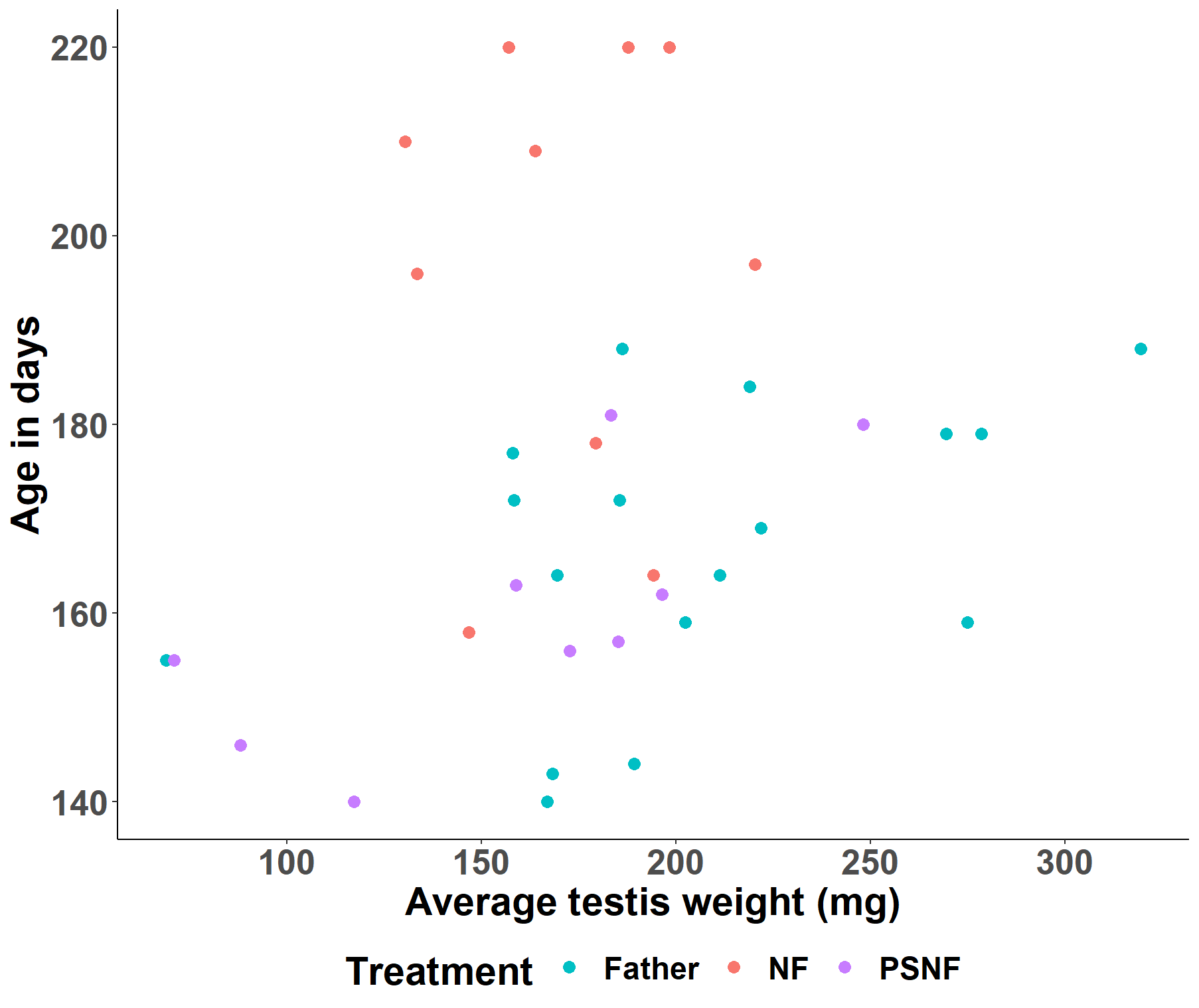

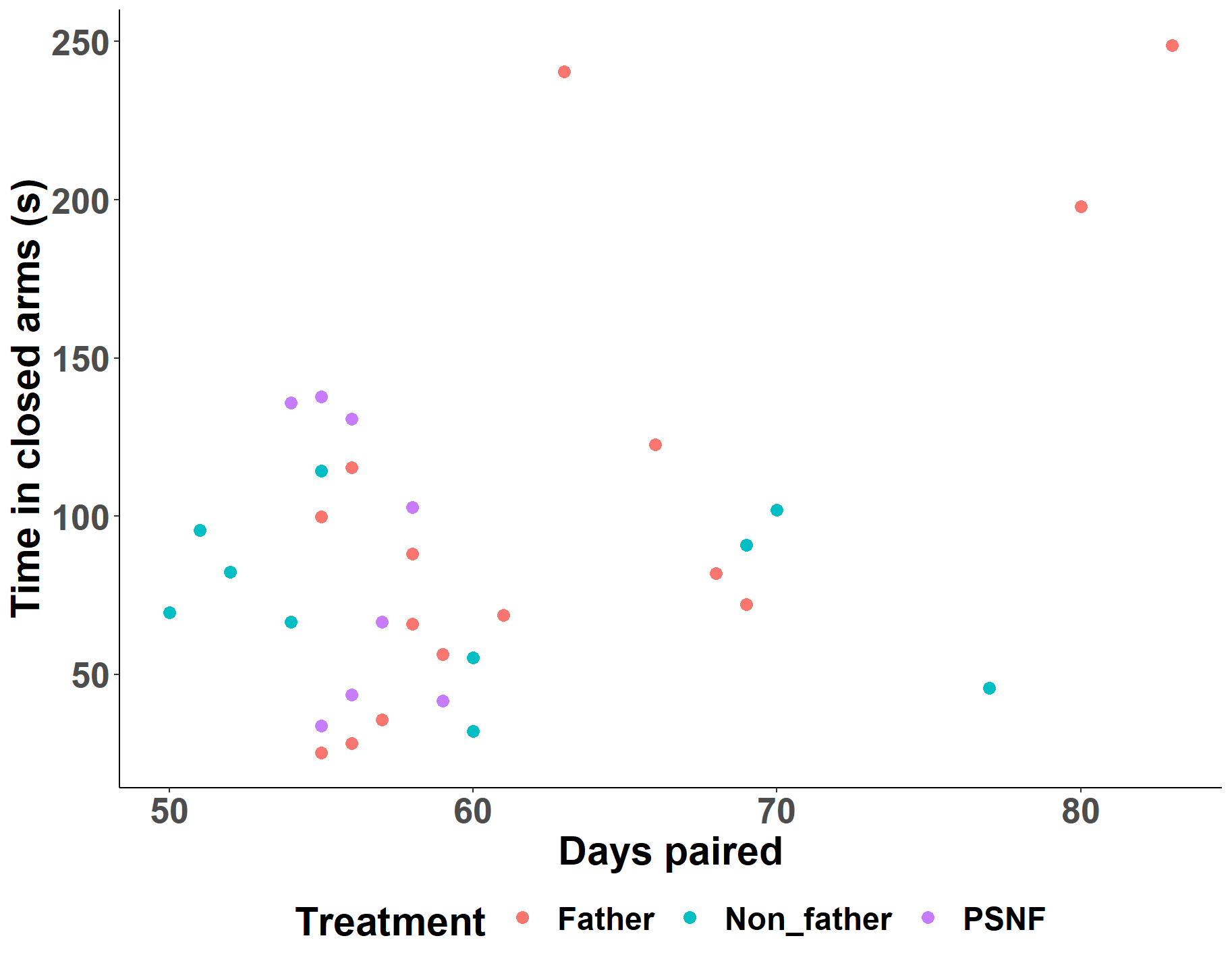

**Non-father**

**Pup-sensitized non-father**

**First-time father**

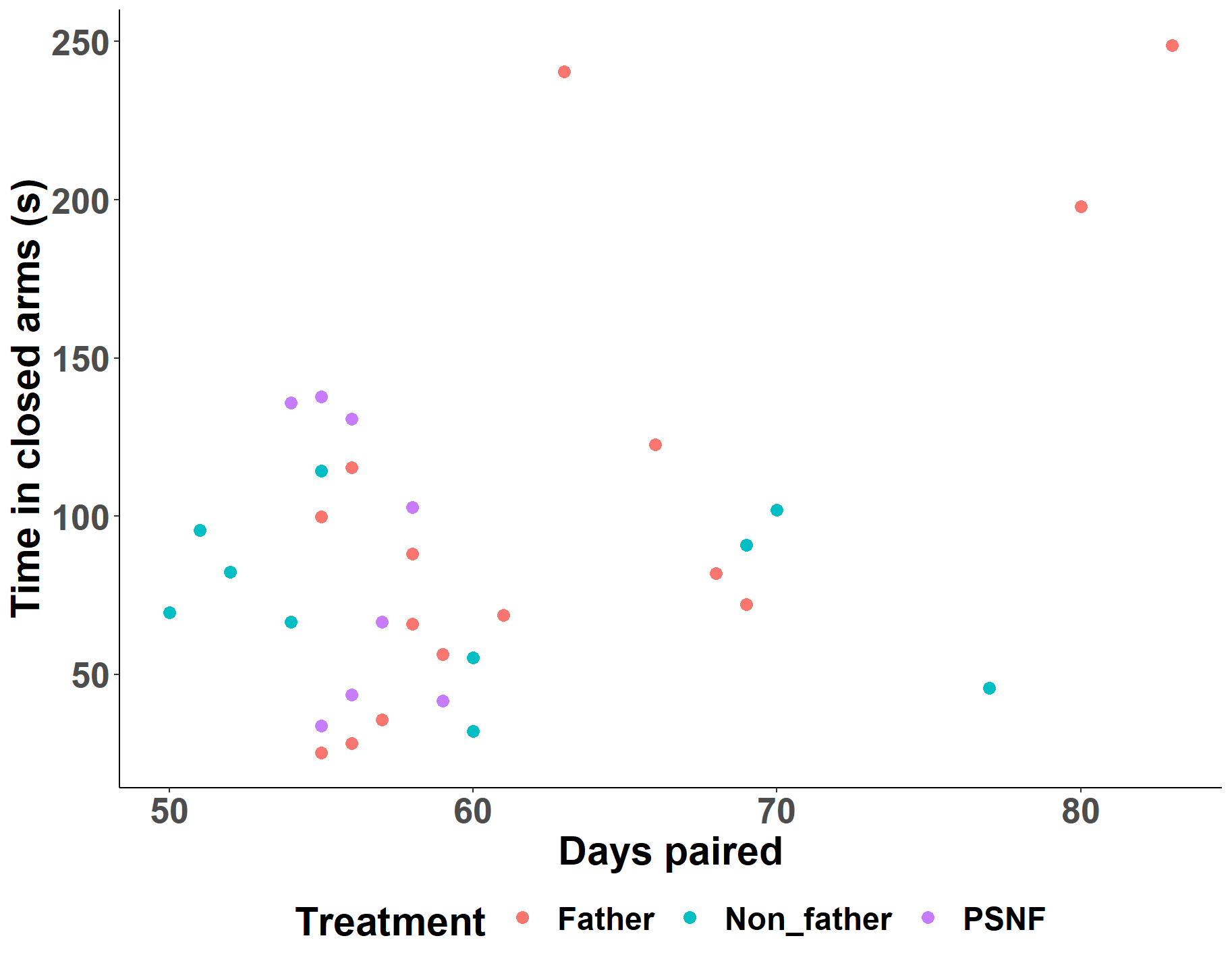

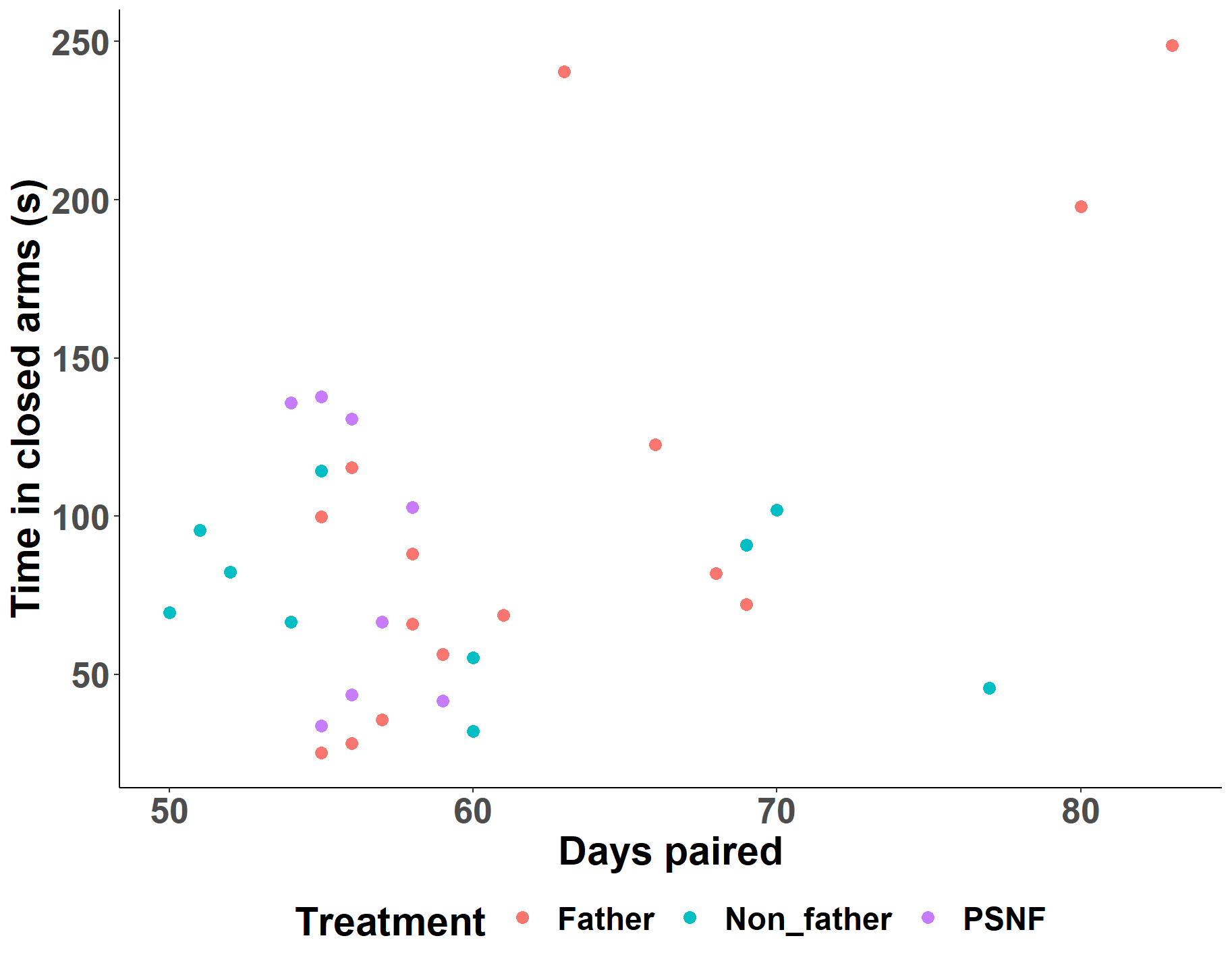

**Figure S12.** Effect of age on average testis weight in Experiment 2 males.

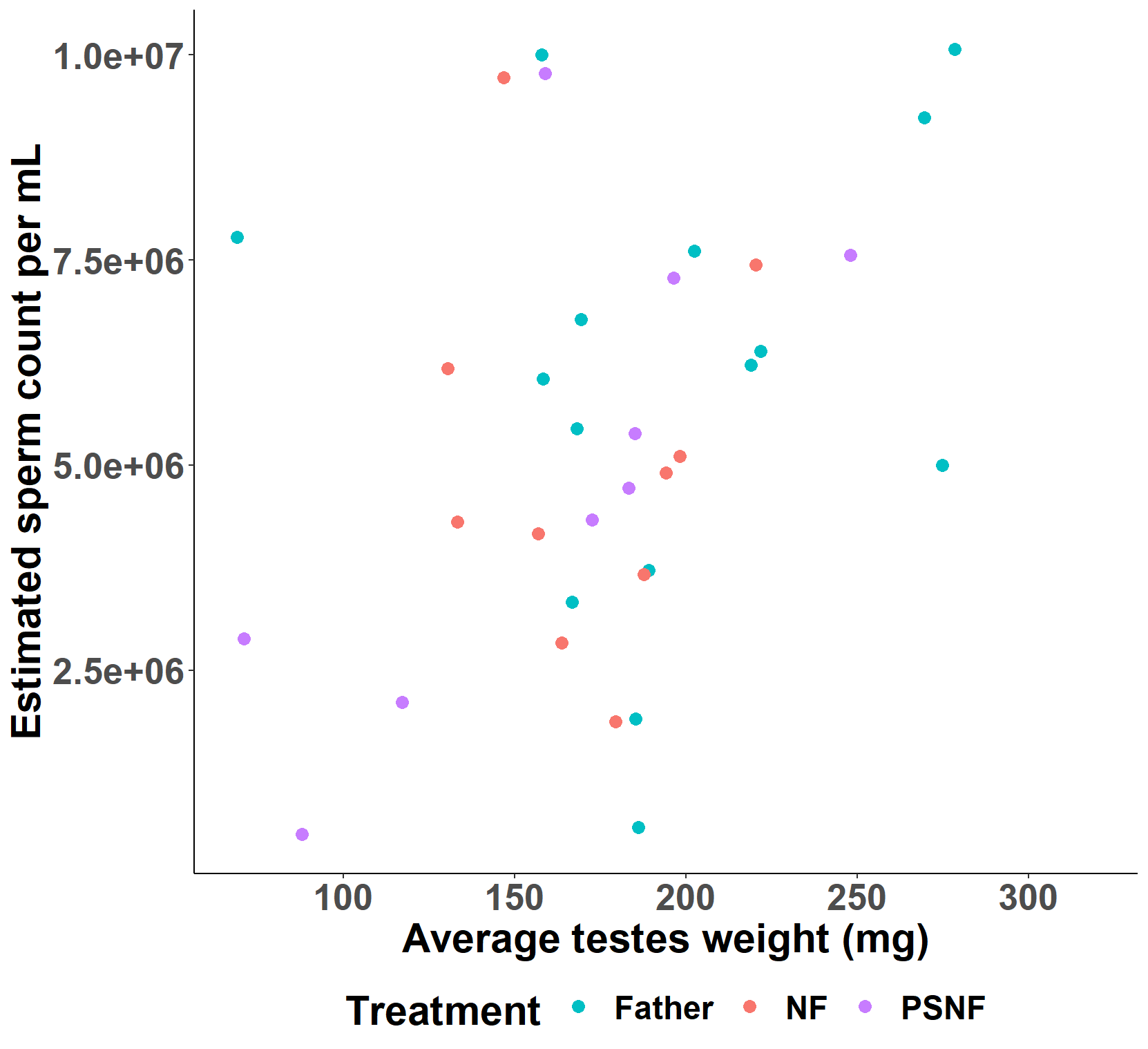

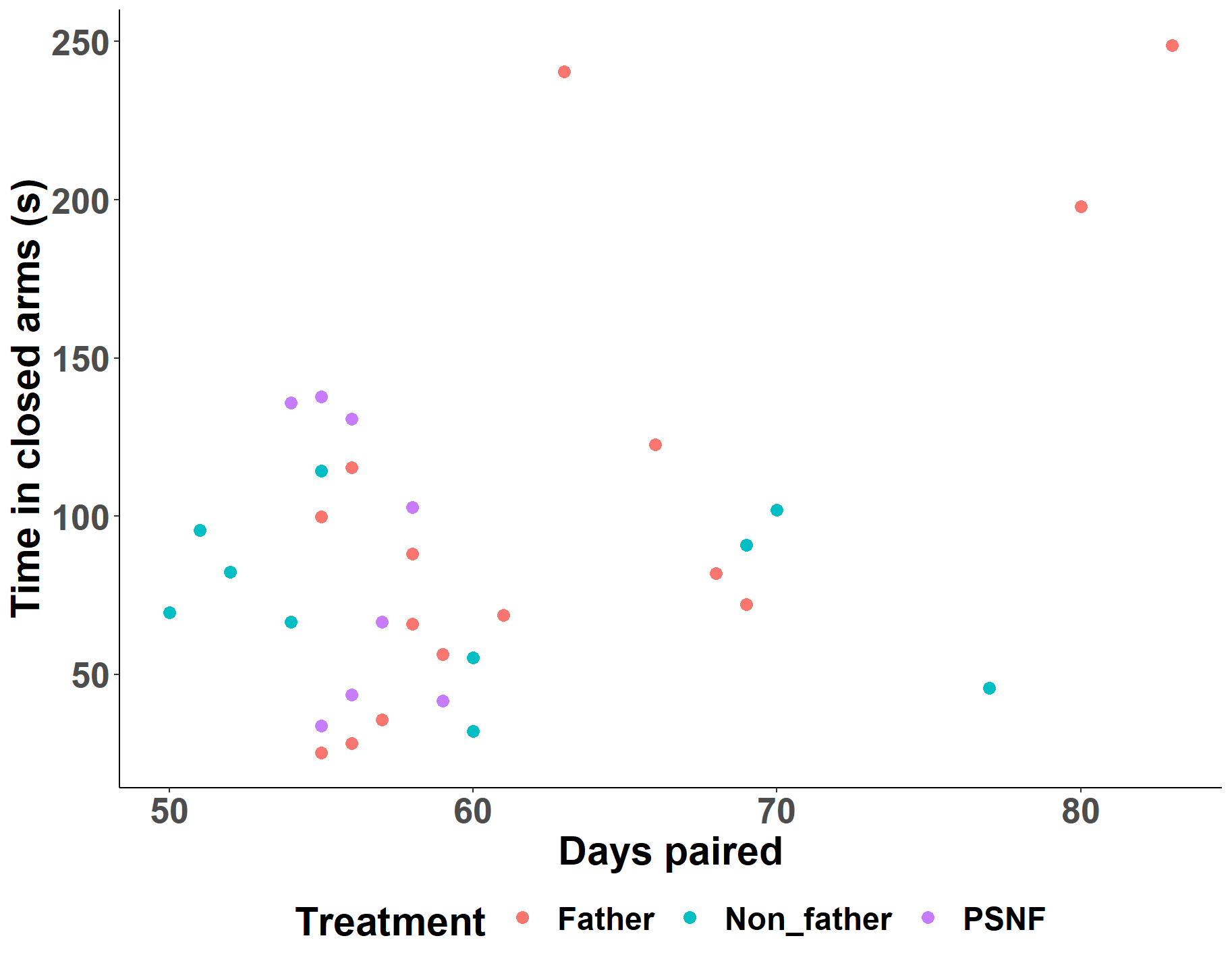

**Non-father**

**Pup-sensitized non-father**

**First-time father**

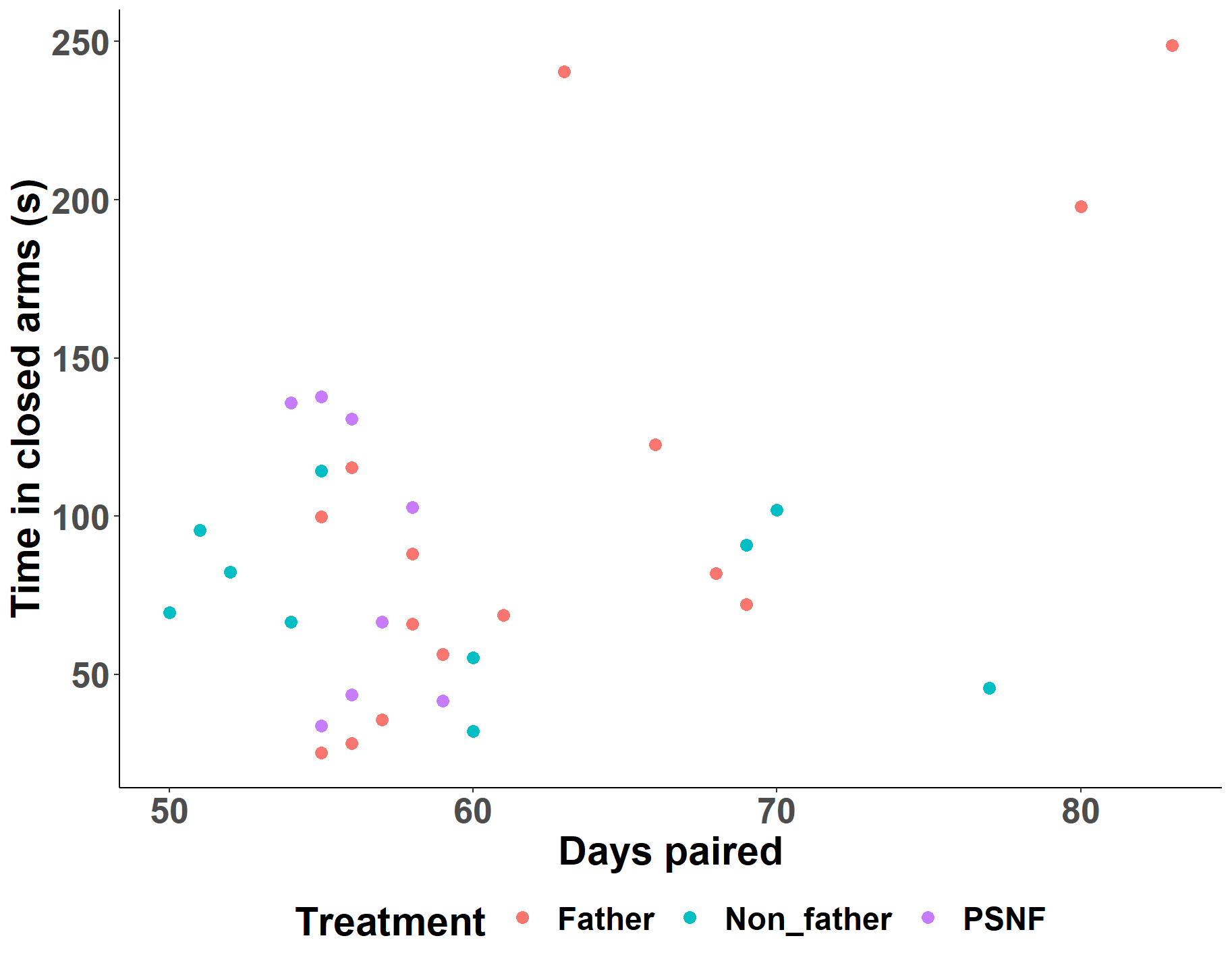

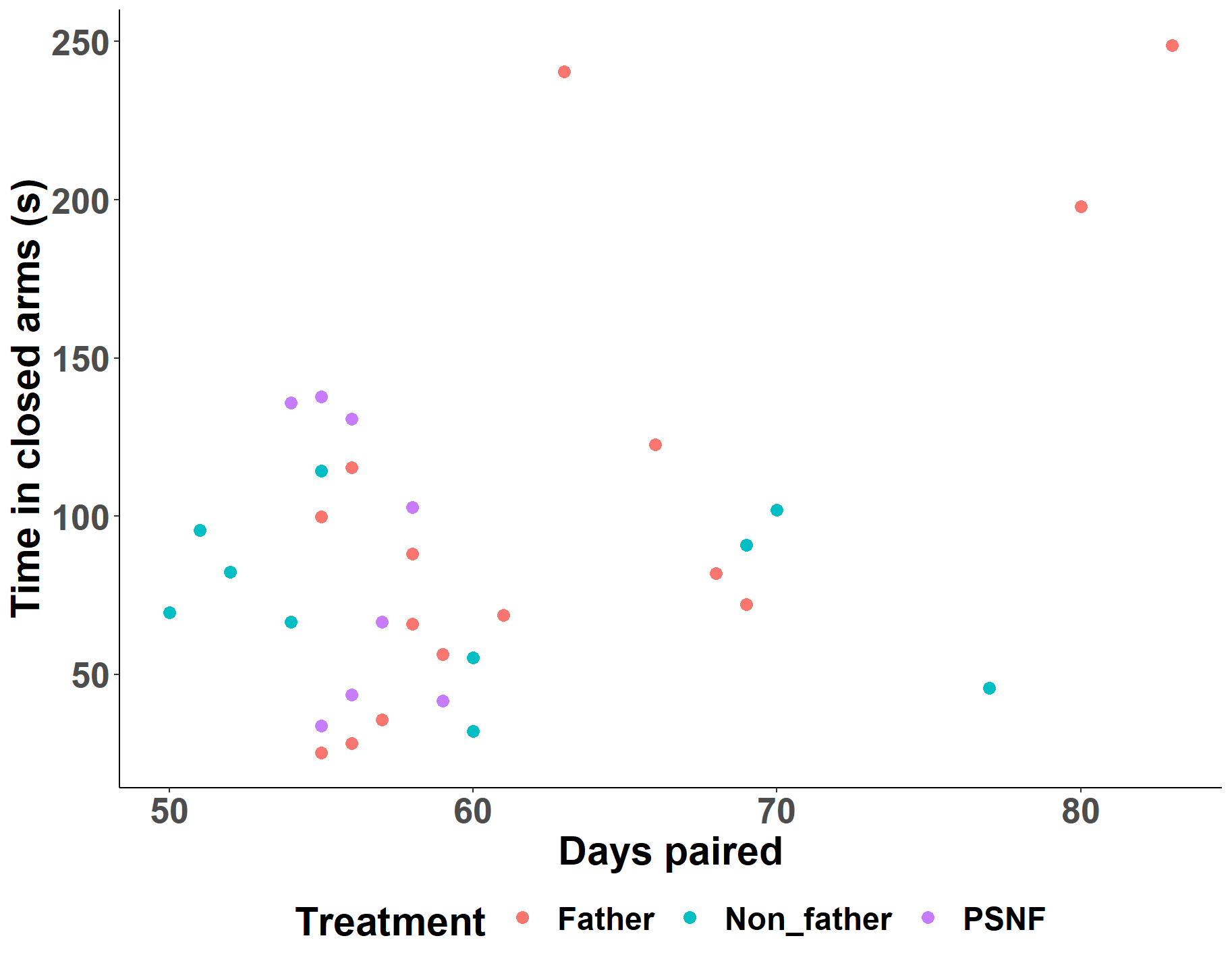

**Figure S13.** Effect of estimated sperm count on average testis weight in Experiment 2 males.

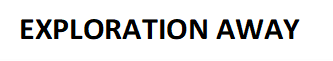

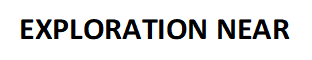

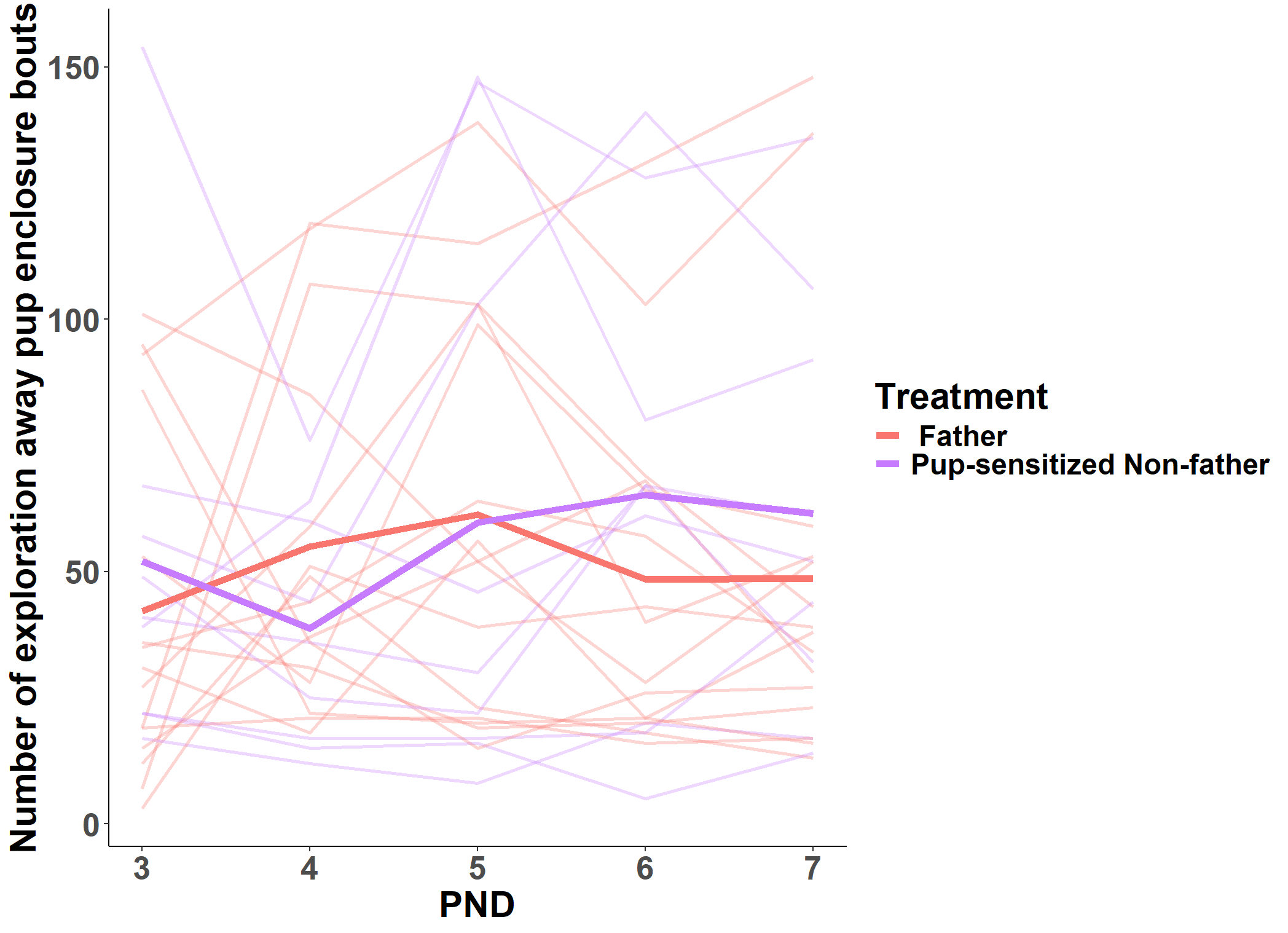

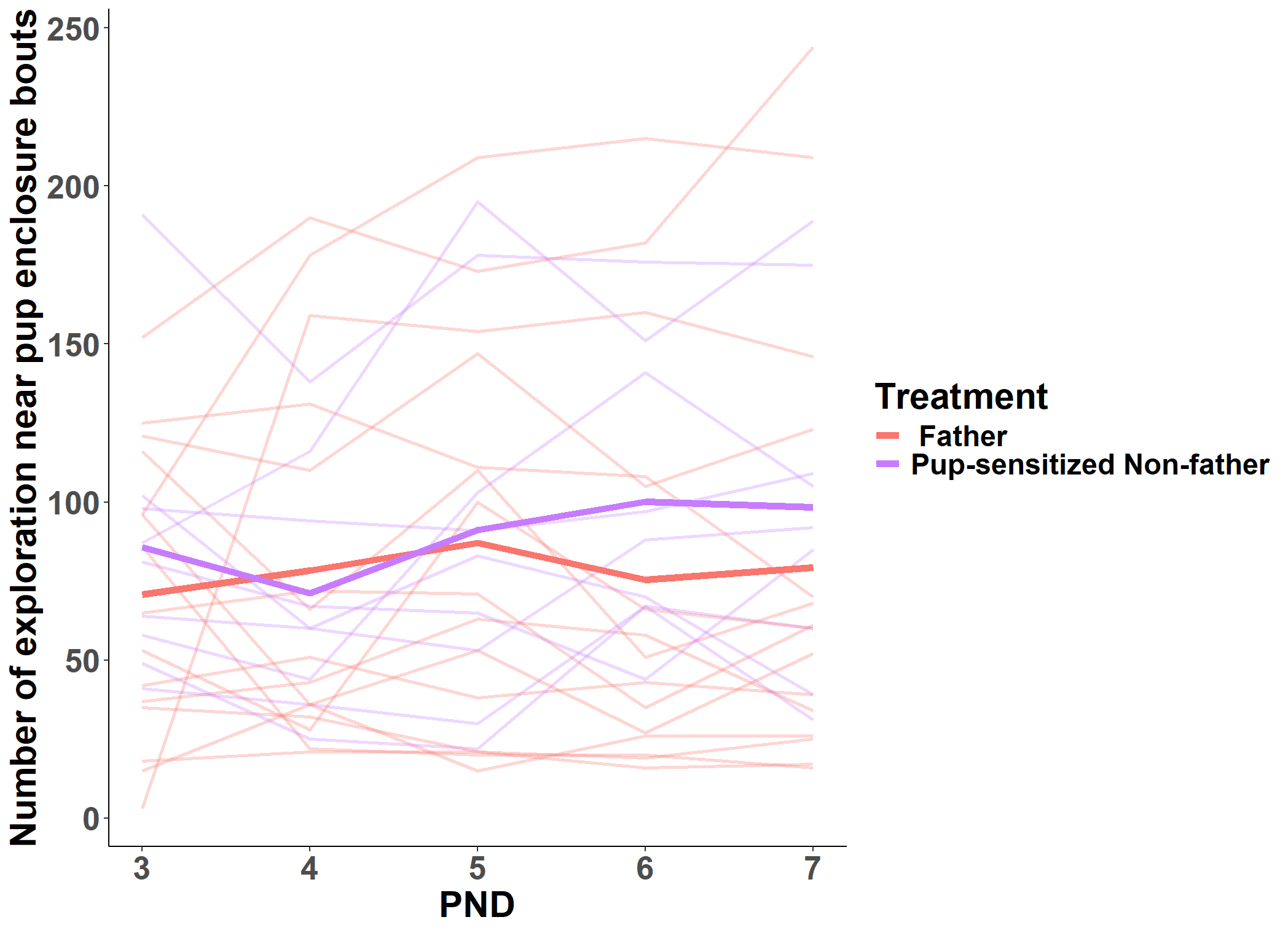

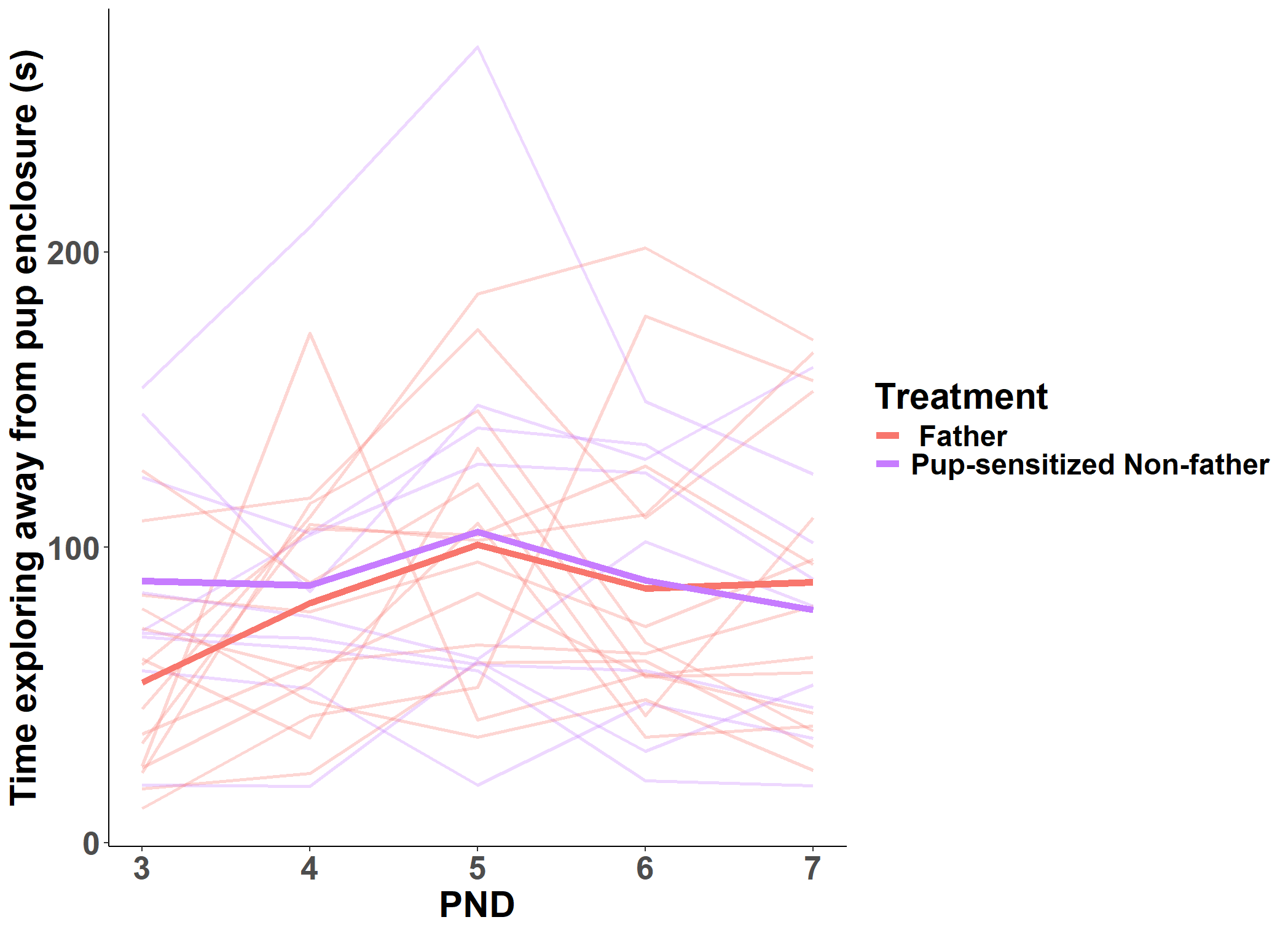

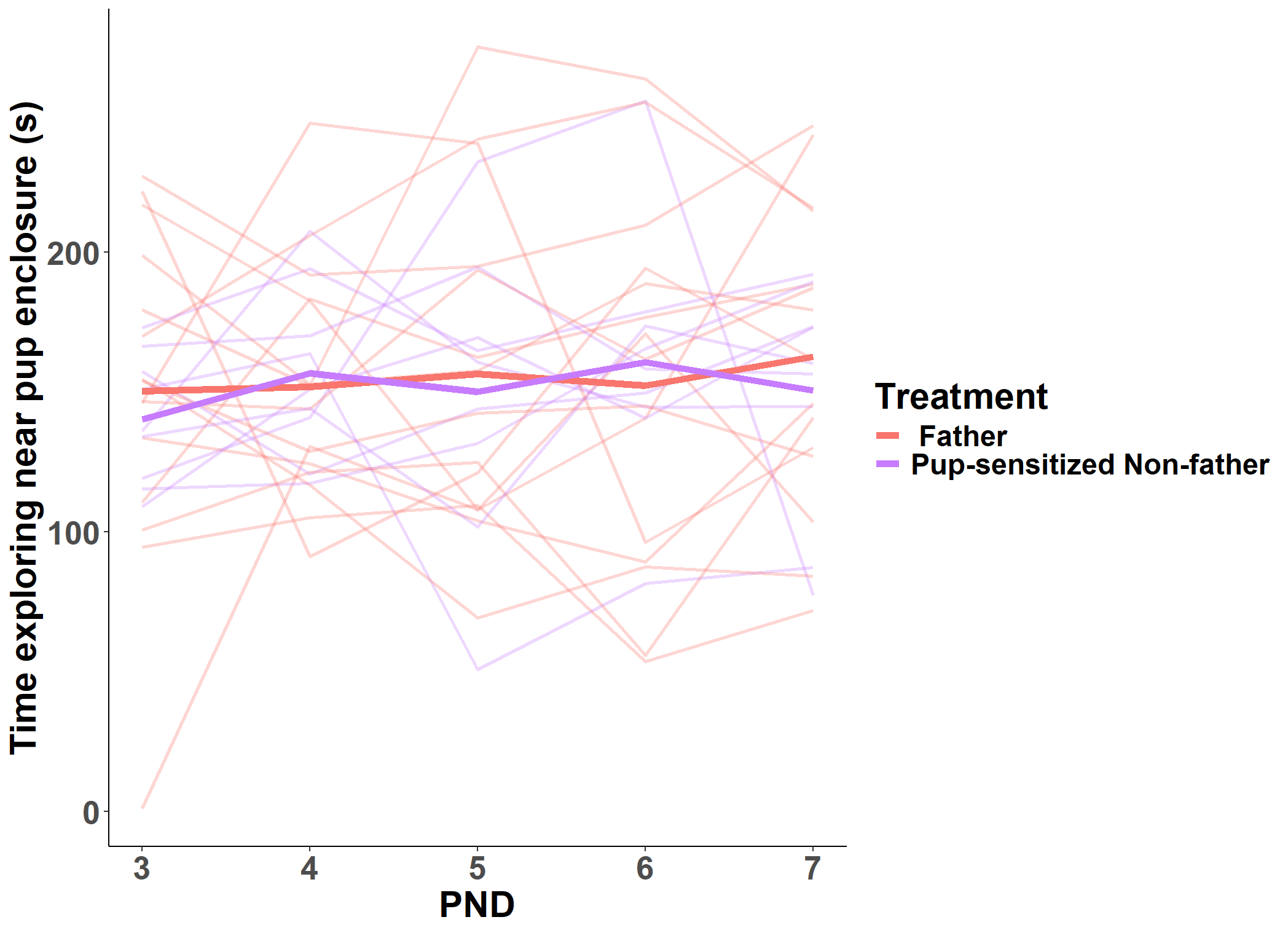
**Figure S14.** Exploration near the pup enclosure **(A)** duration and **(B)** frequency and exploration away from the pup enclosure **(C)** duration and **(D)** frequency. Thin lines represent individuals and bold lines represent experience means.

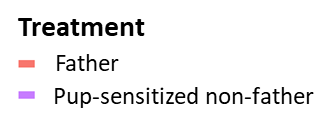

Pup-sensitized non-father

First-time father

**FREQUENCY**

**A**

**C**

**D**

**B**

**DURATION**

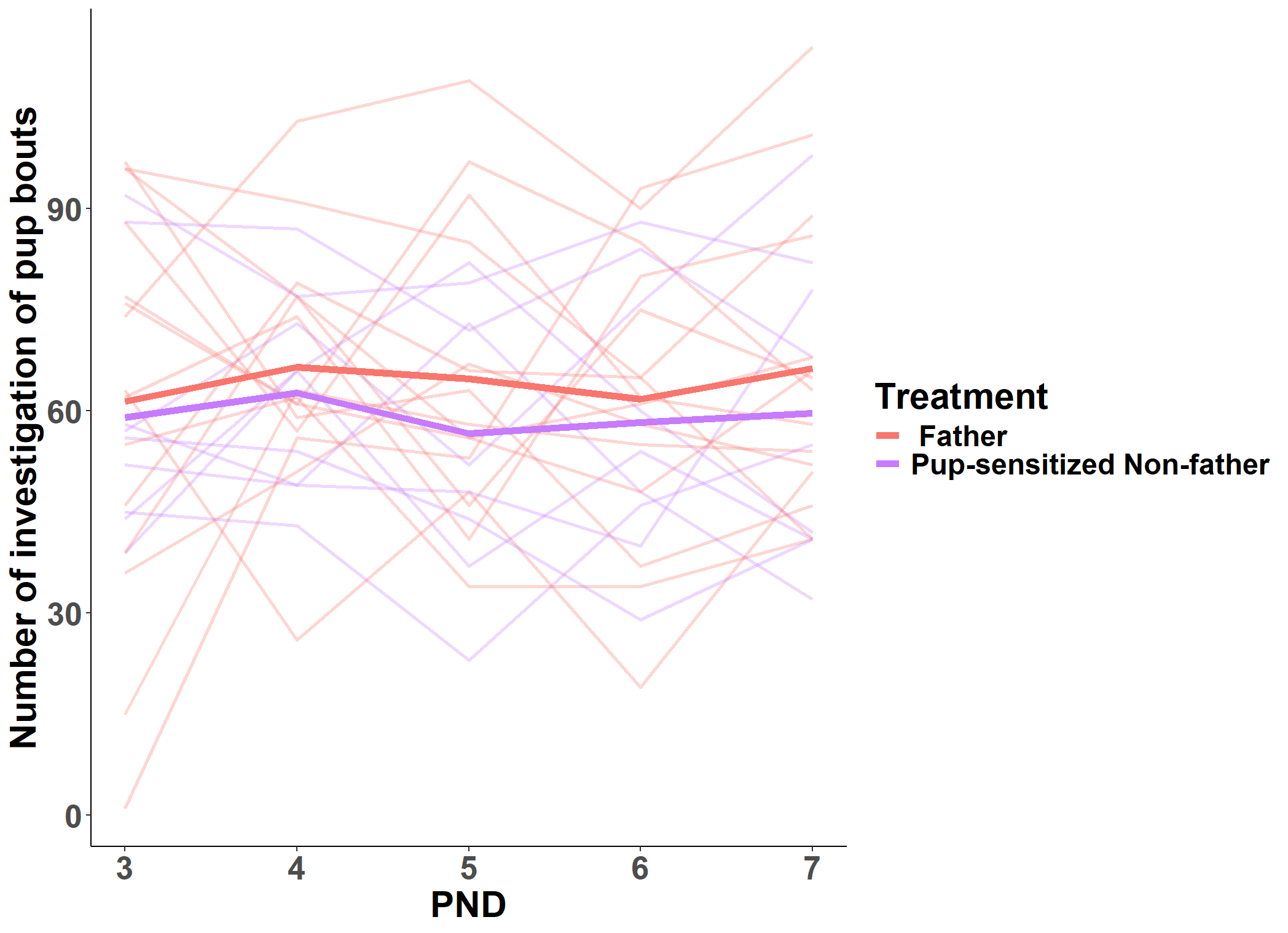

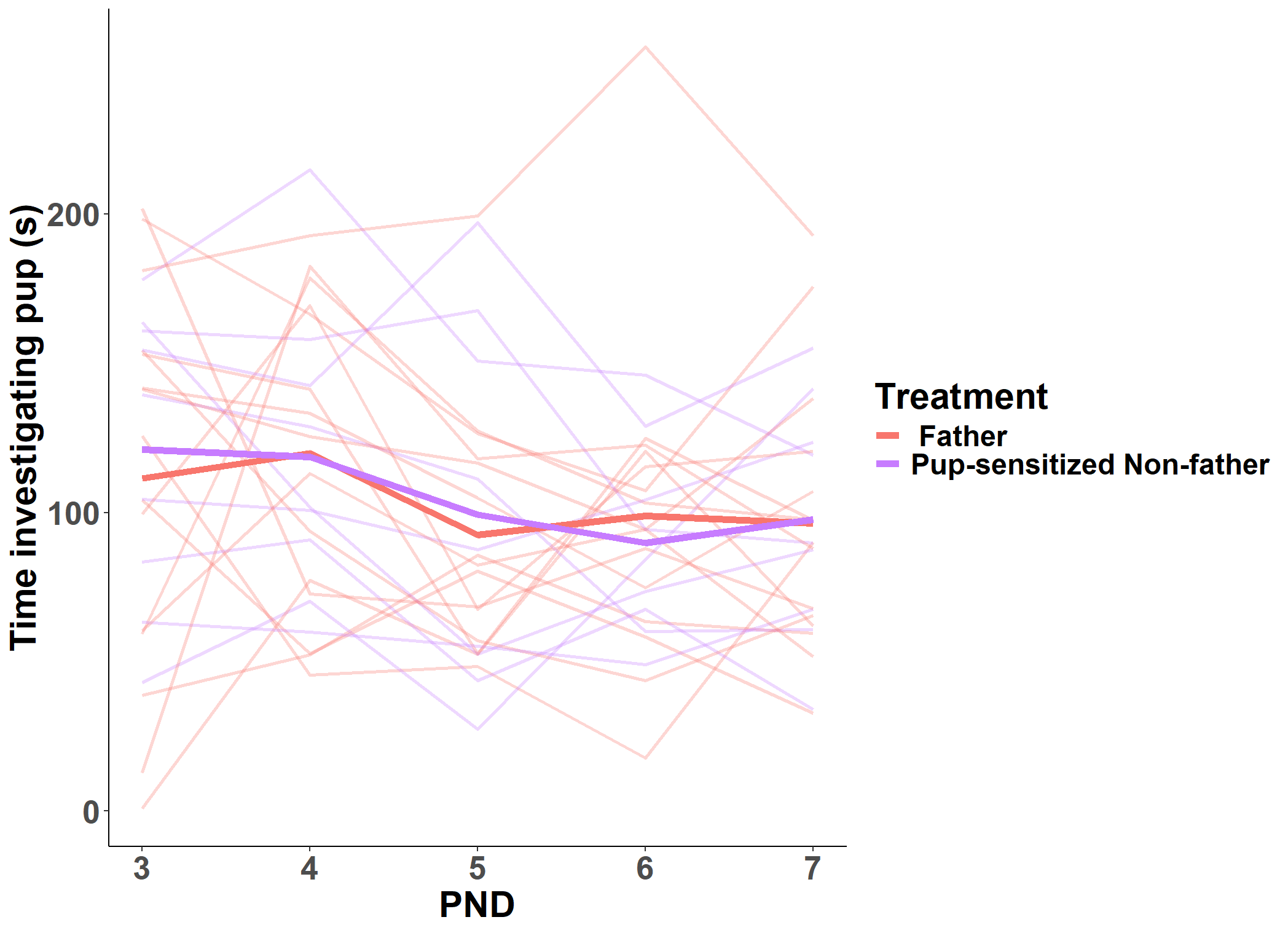

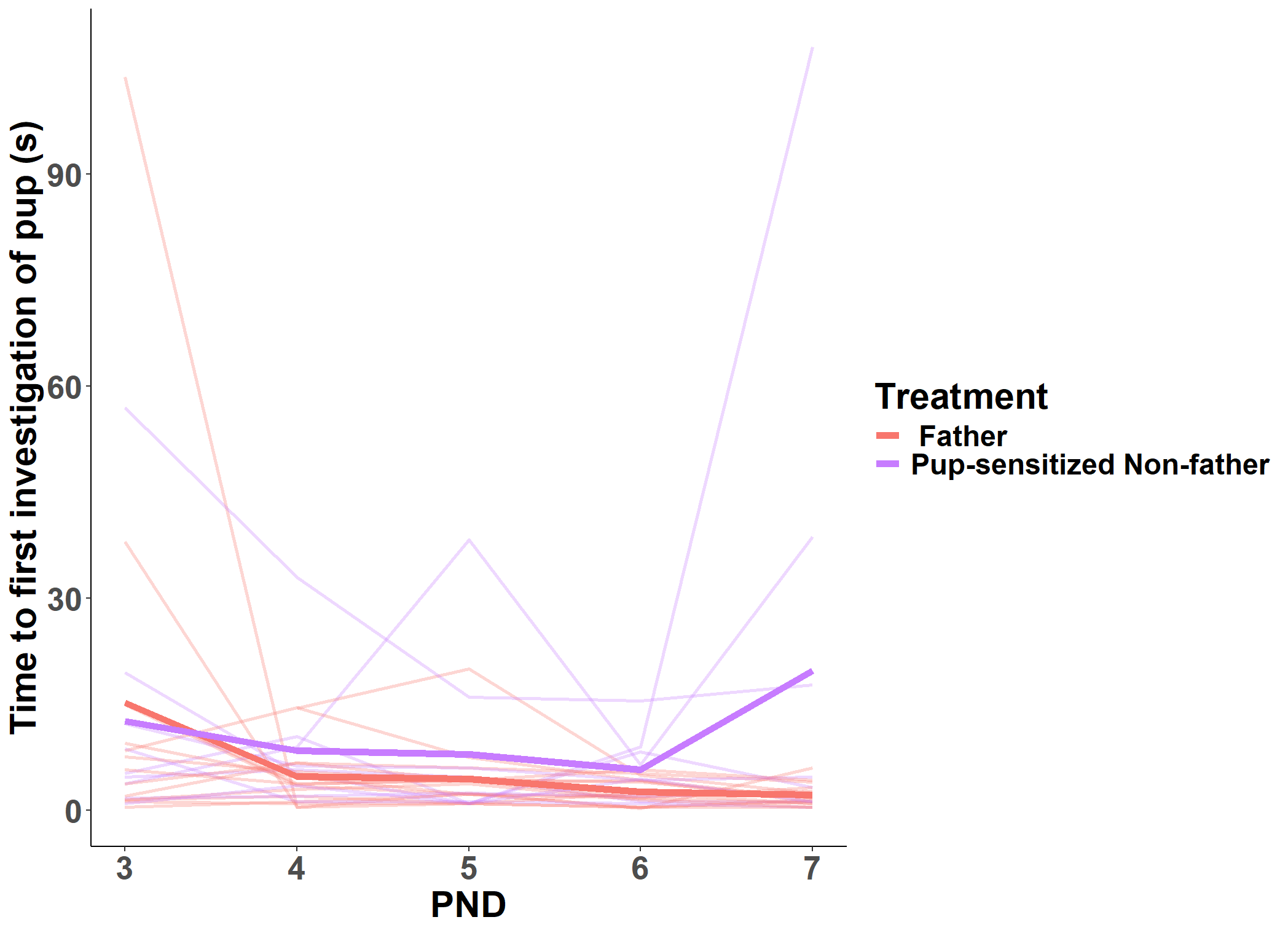

**FREQUENCY**

**LATENCY**

**B**

**C**

**A**

**DURATION**

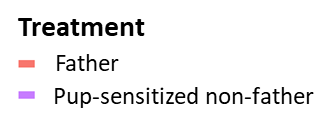

Pup-sensitized non-father

First-time father

**Figure S15.** Pup investigation behavior **(A)** latency, **(B)** duration, and **(C)** frequency of pup-sensitized non-fathers and first-time fathers during the pup-exposure assay. Thin lines represent individuals and bold lines represent experience means.

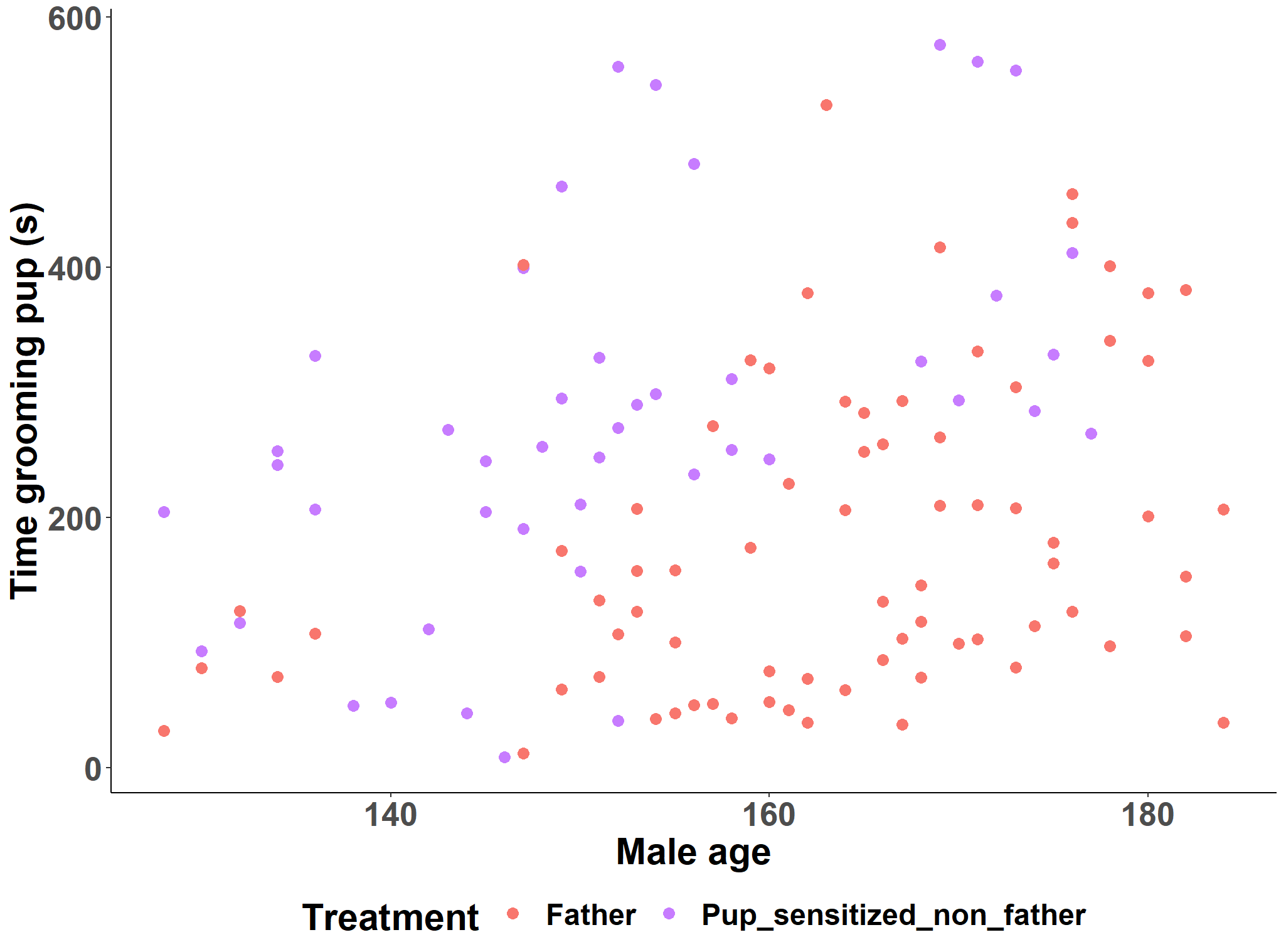

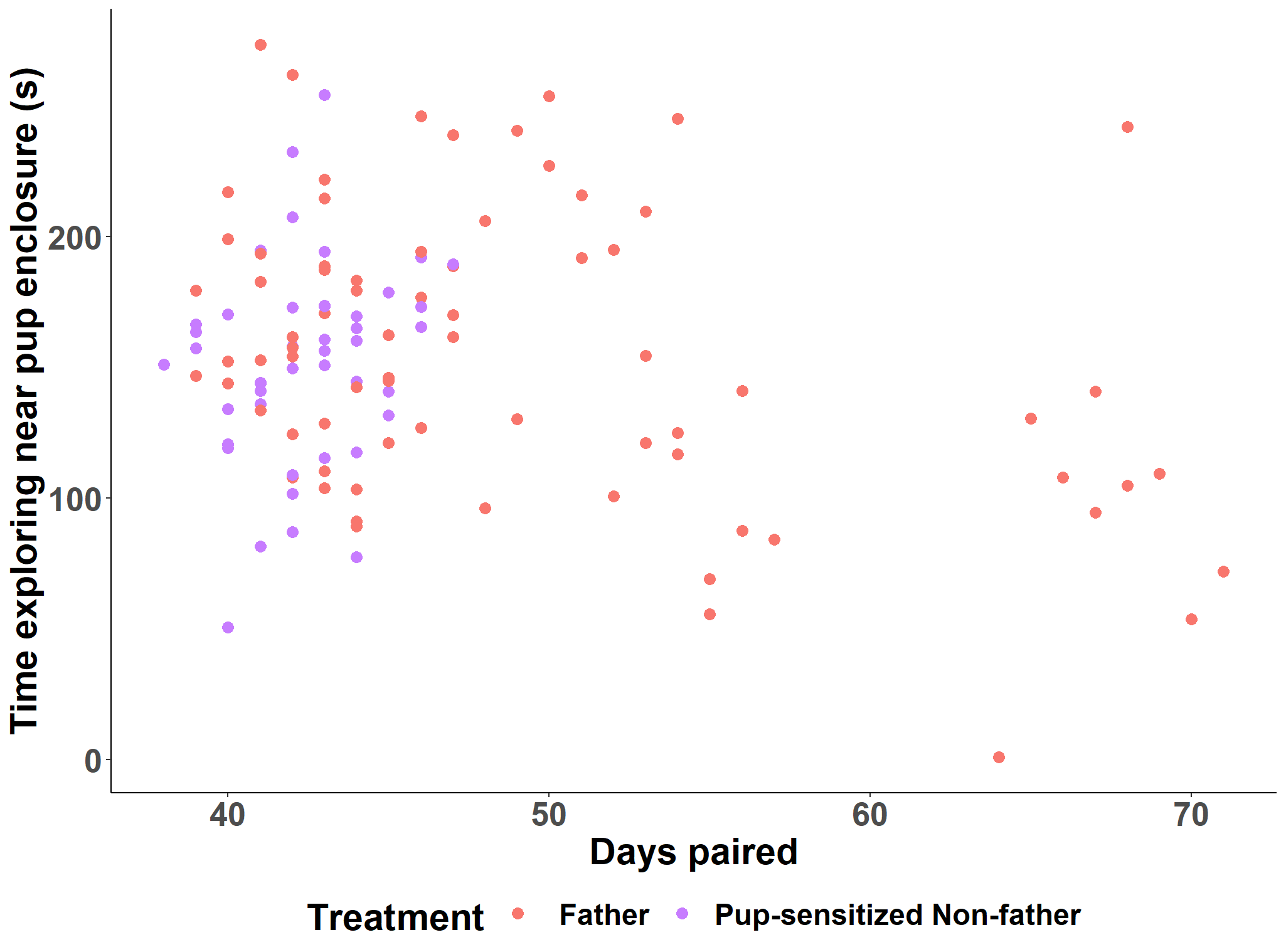

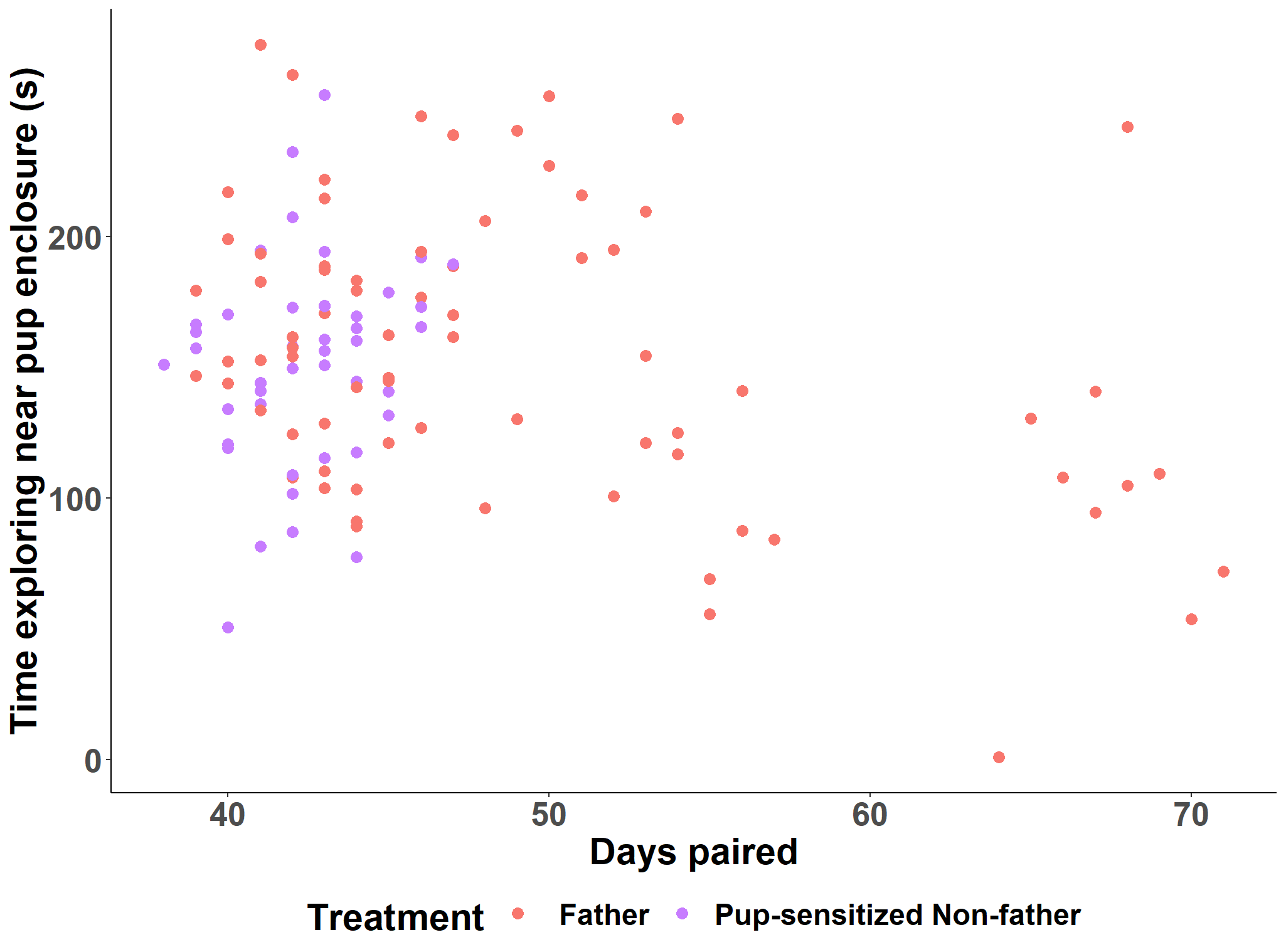

Pup-sensitized non-father

First-time father

**Figure S16.** Male age as a predictor of grooming behavior during the caregiving assay.

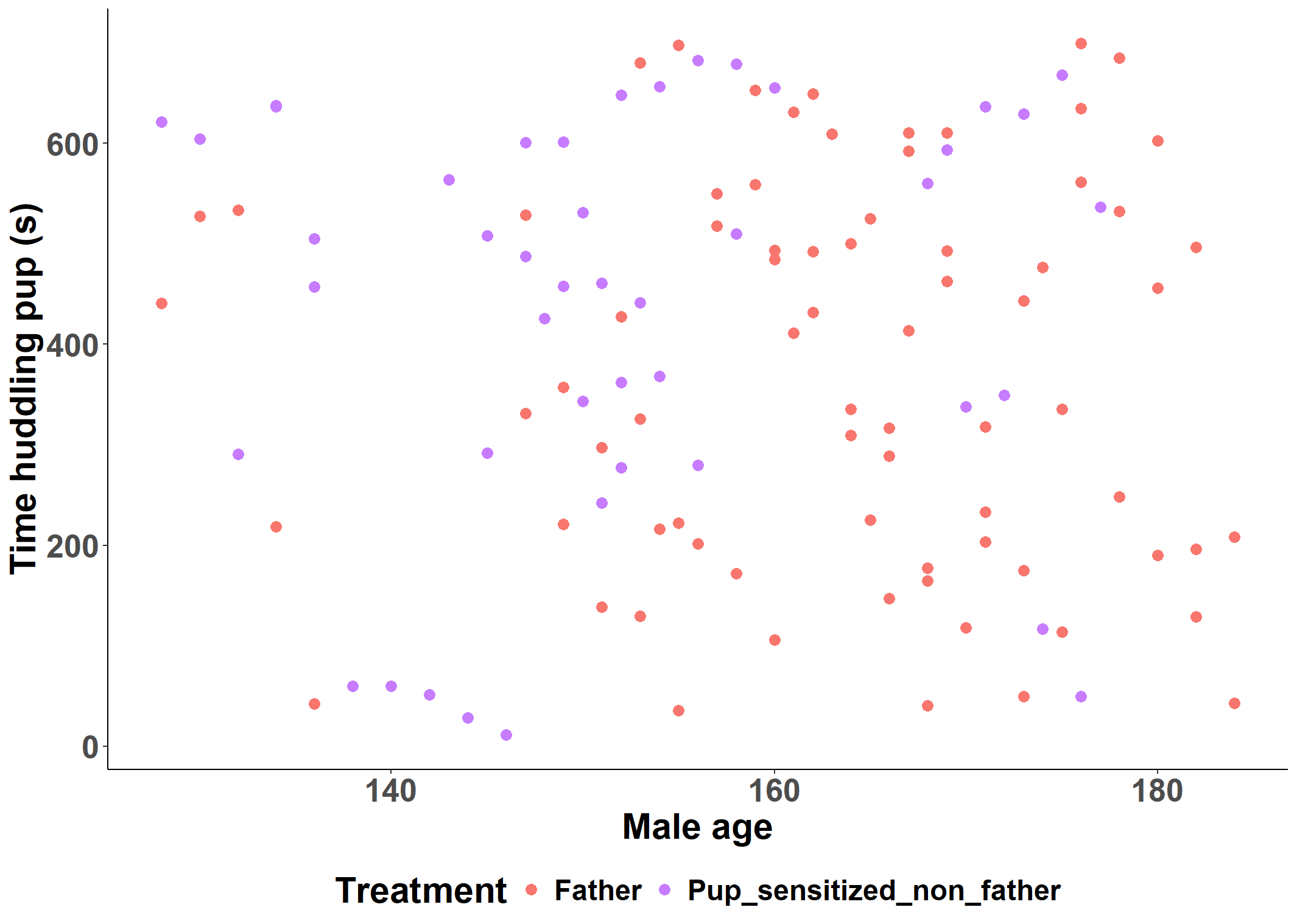

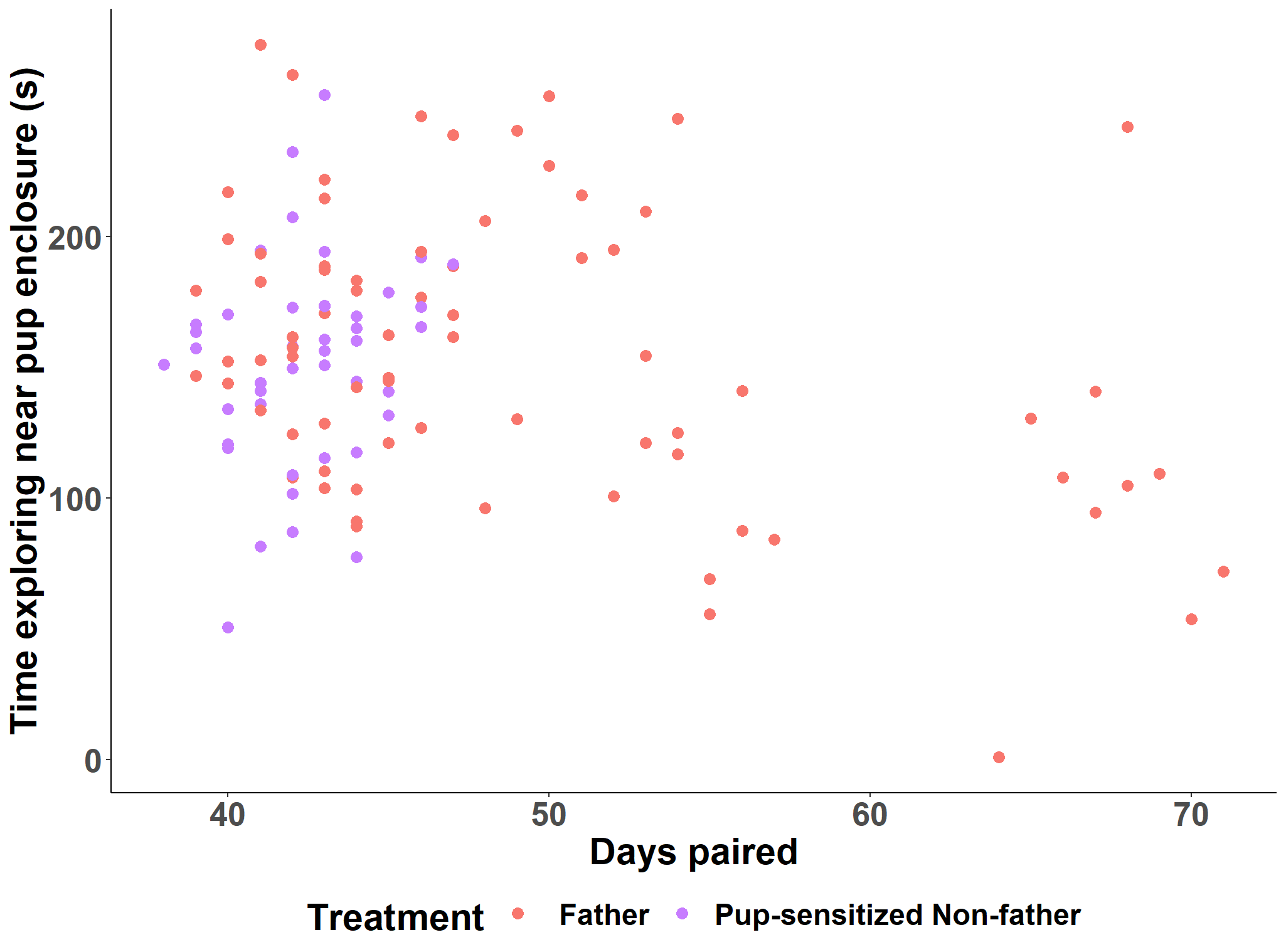

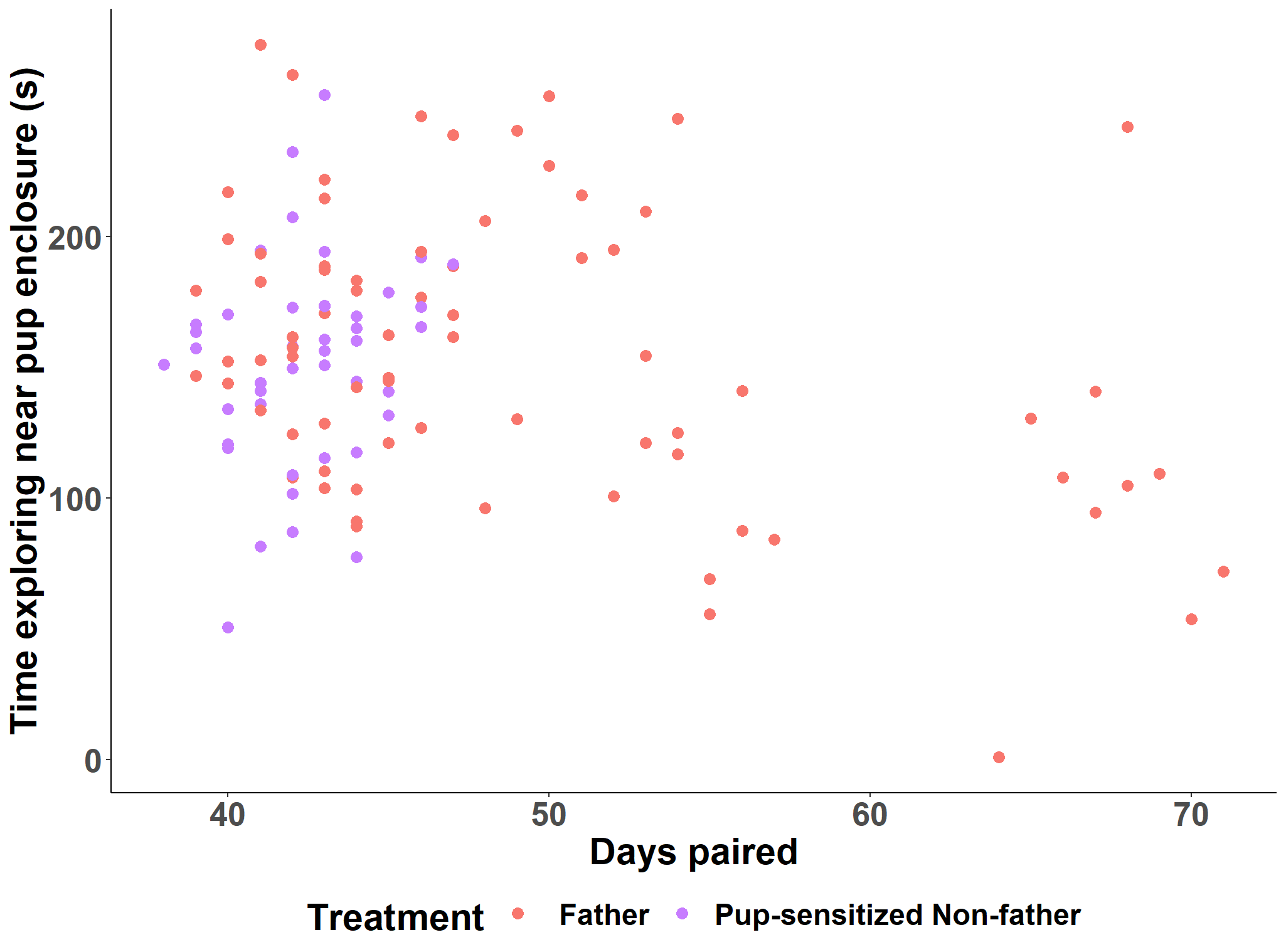

Pup-sensitized non-father

First-time father

**Figure S17.** Male age as a predictor of huddling behavior during the caregiving assay.

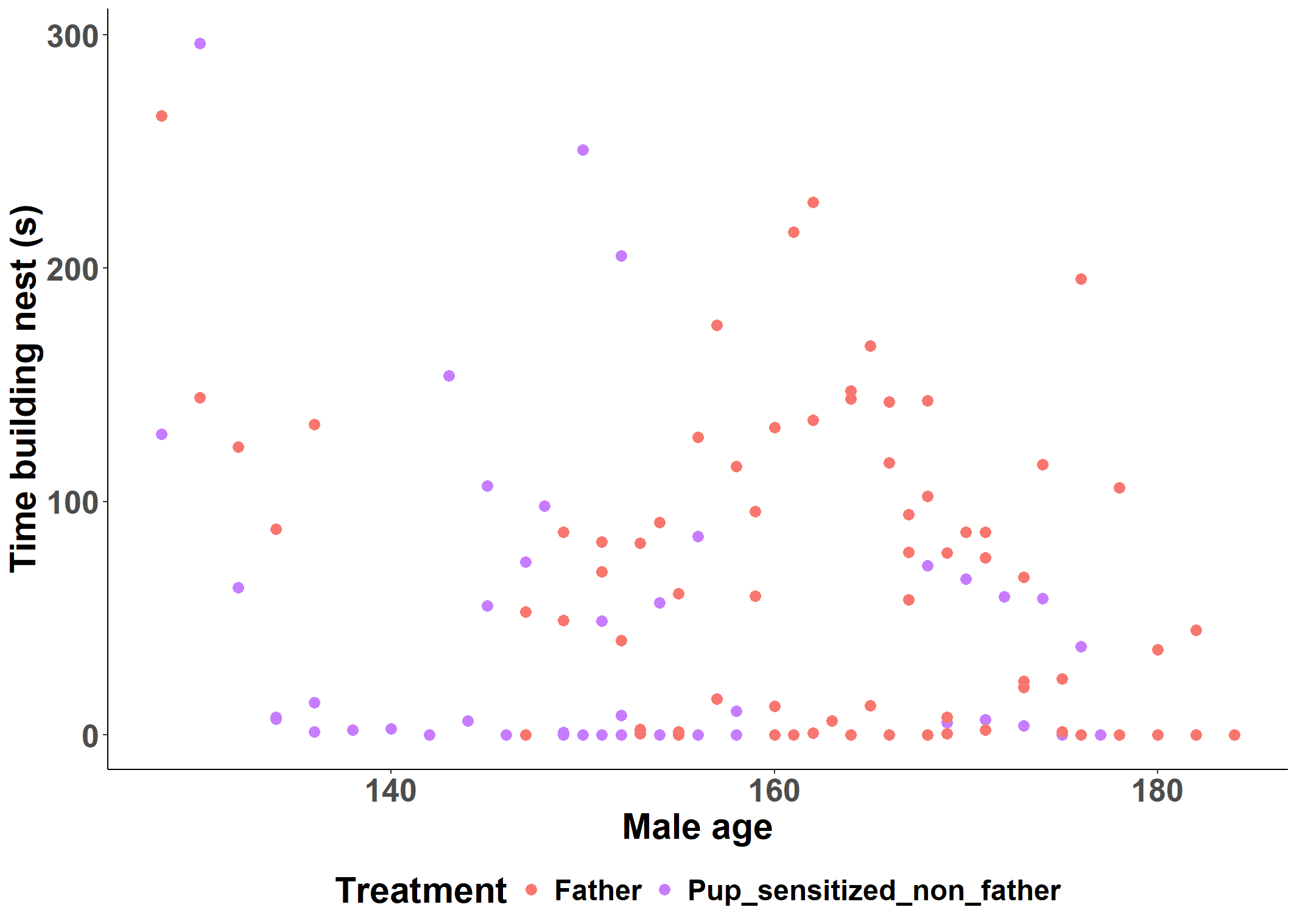

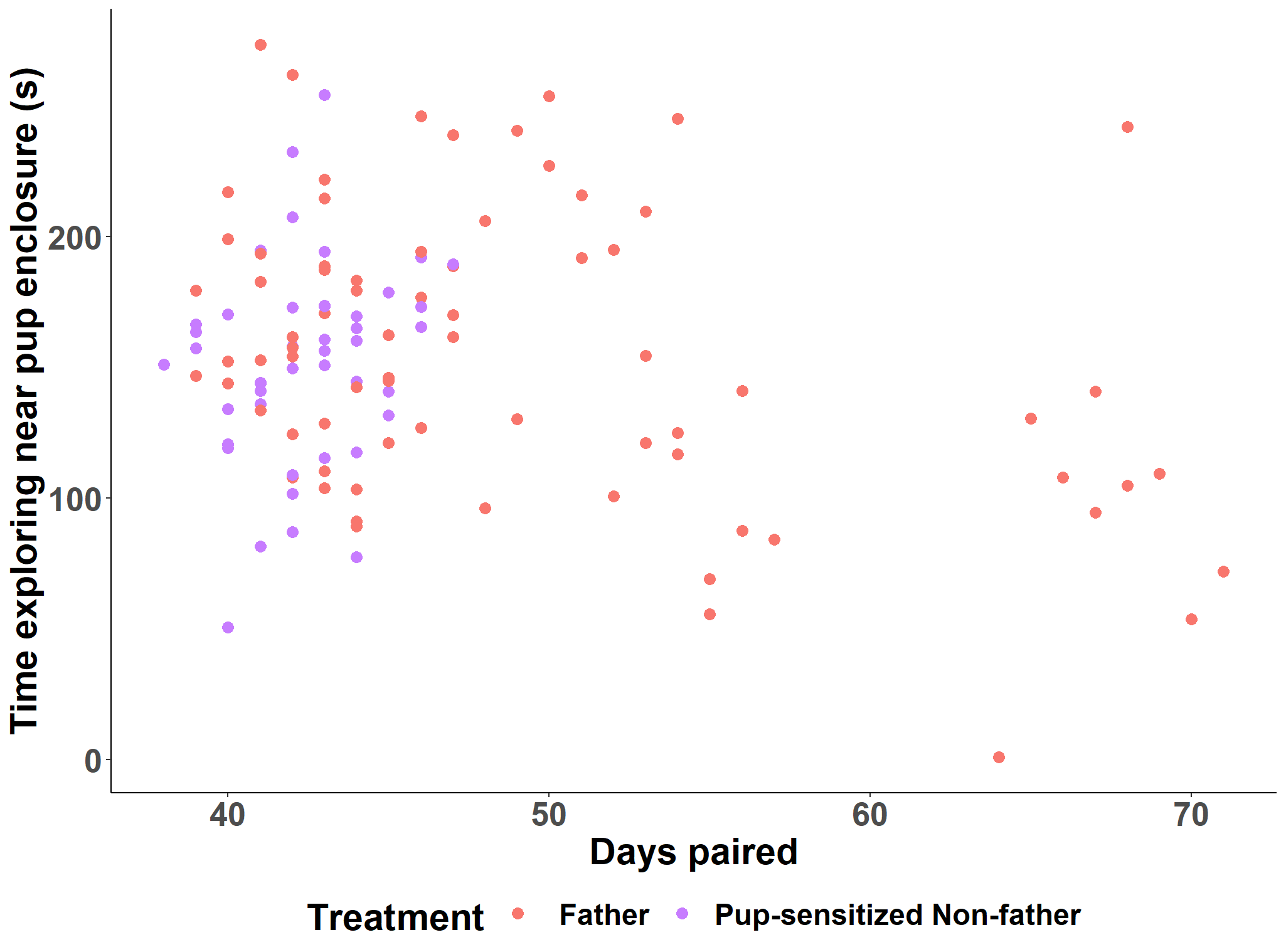

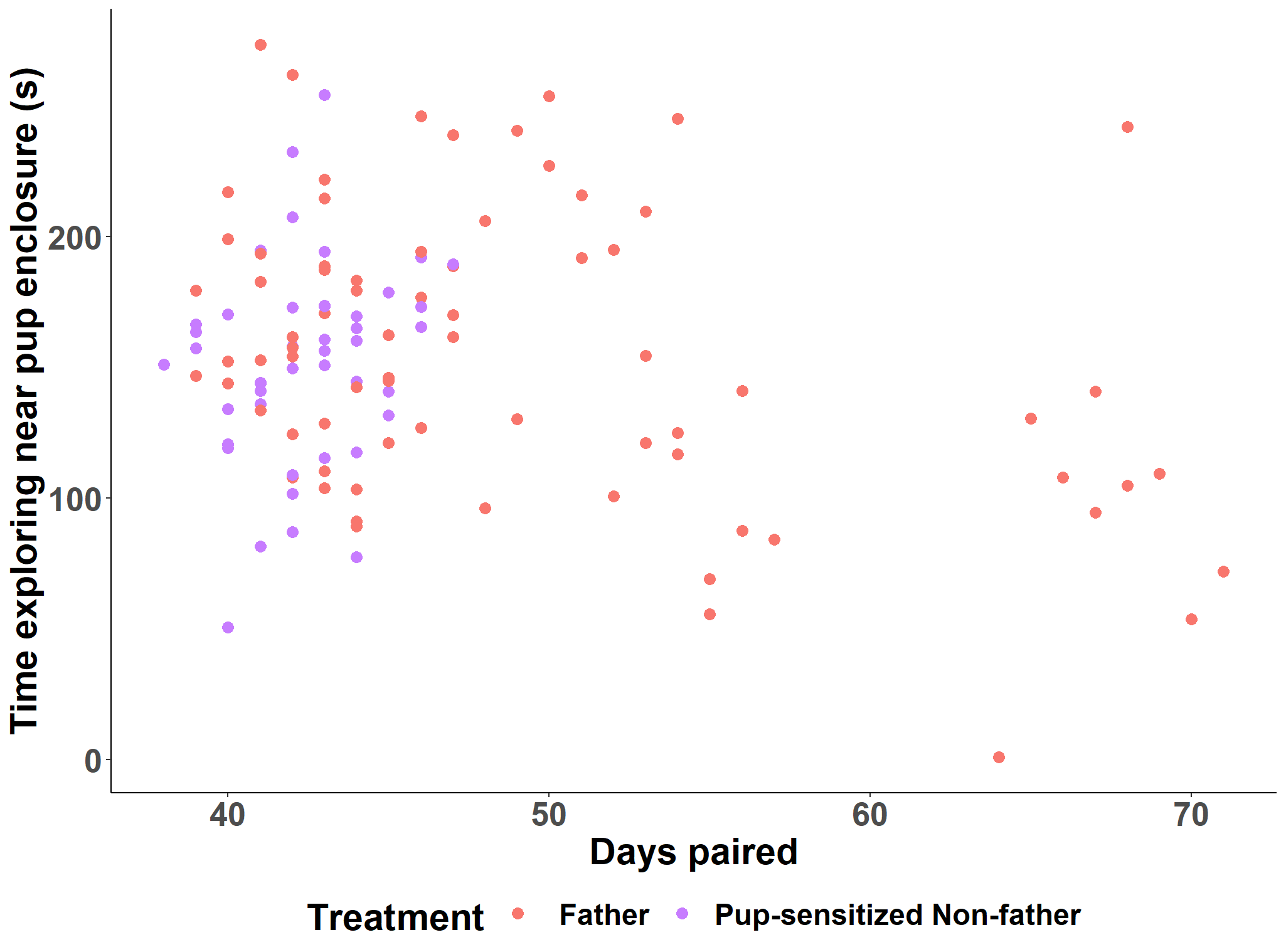

Pup-sensitized non-father

First-time father

**Figure S18.** Male age as a predictor of nest building behavior during the caregiving assay.

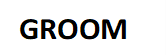

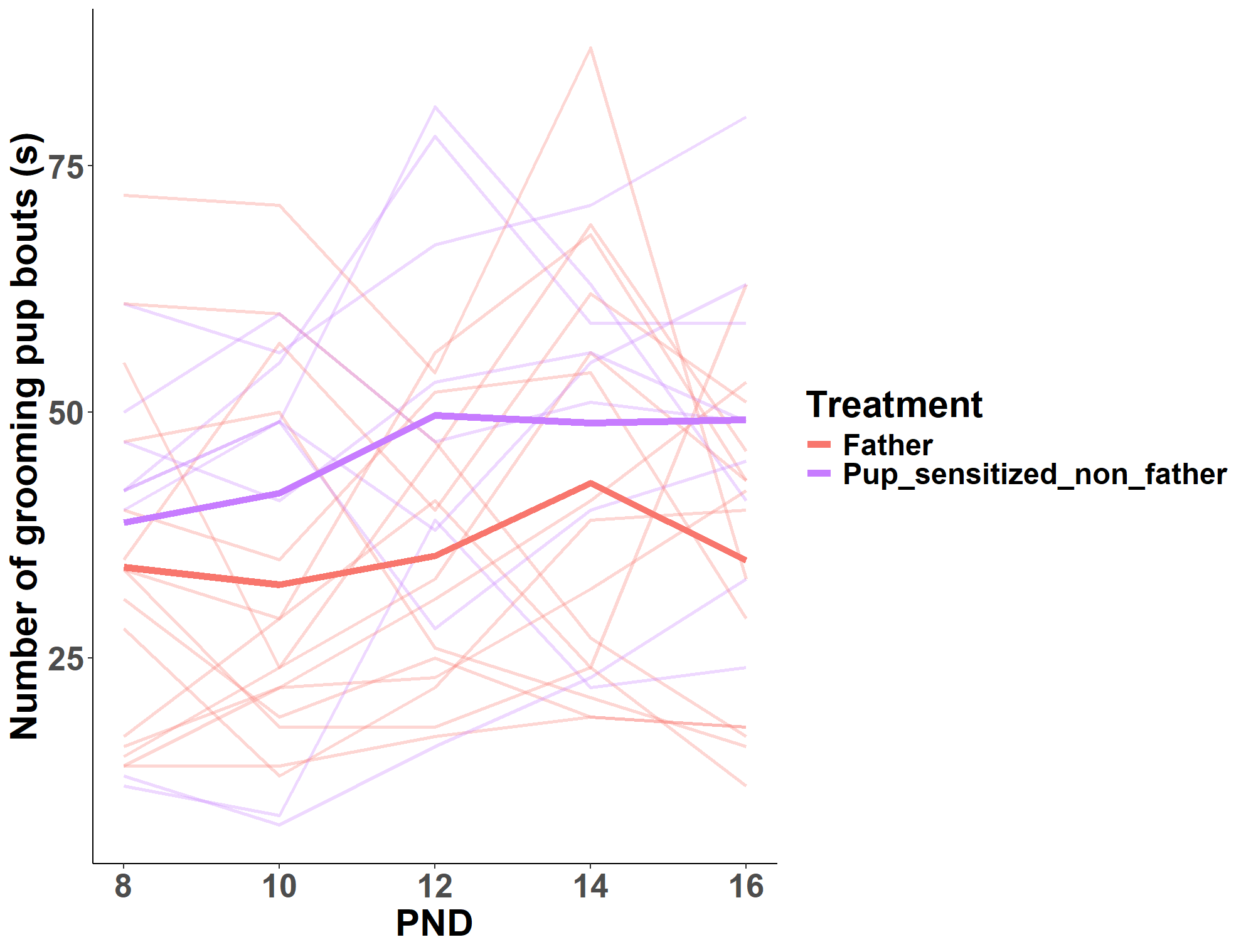

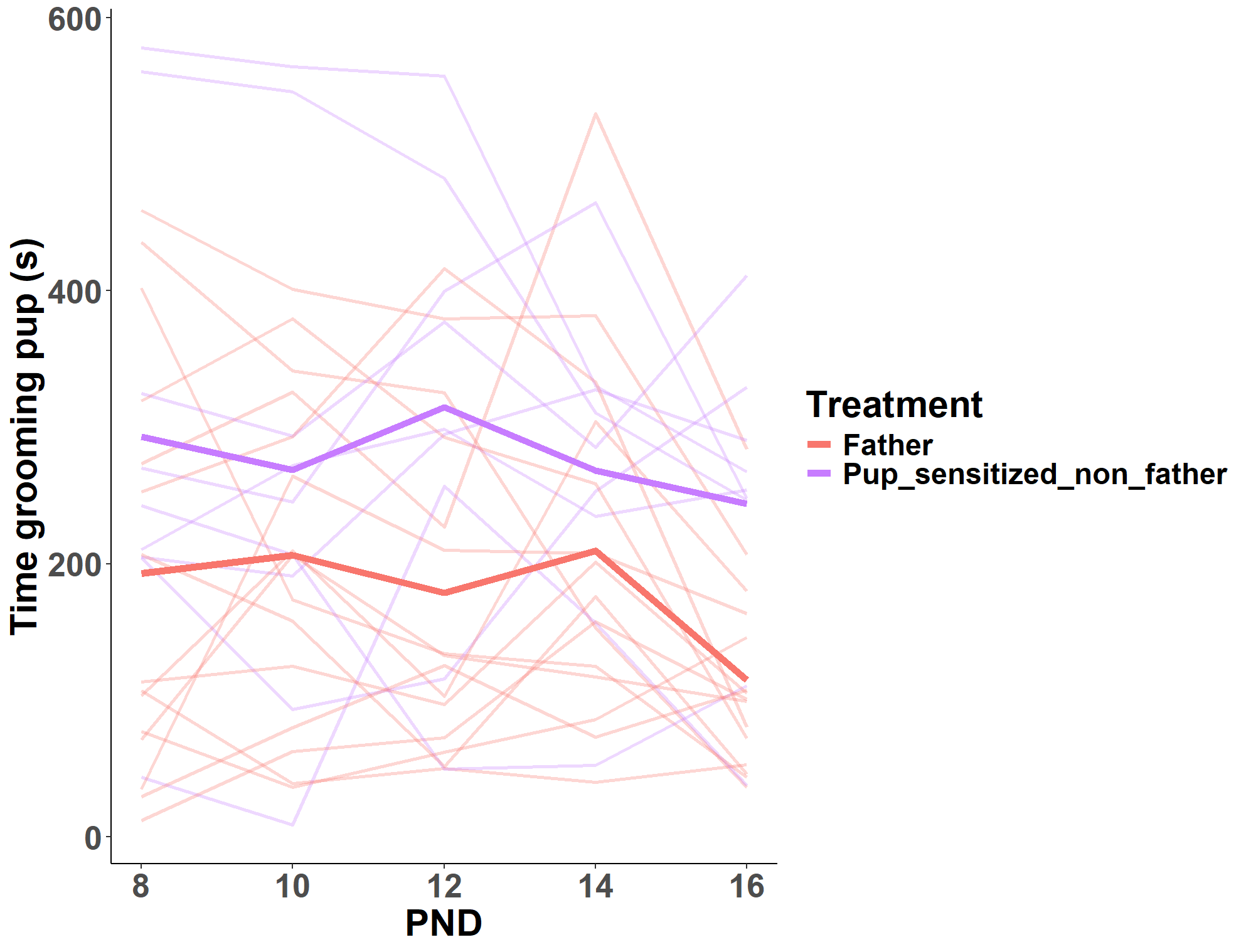

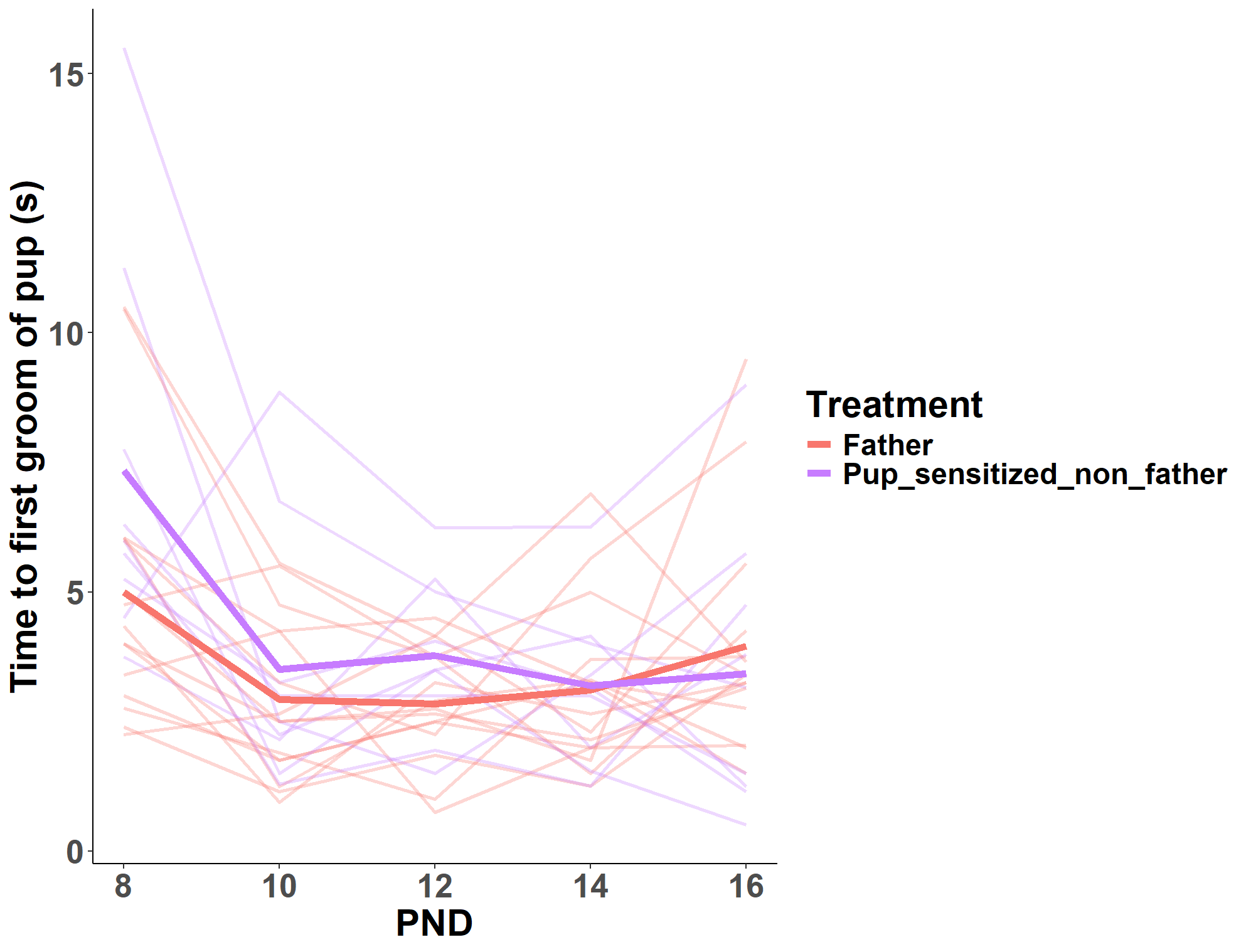

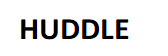

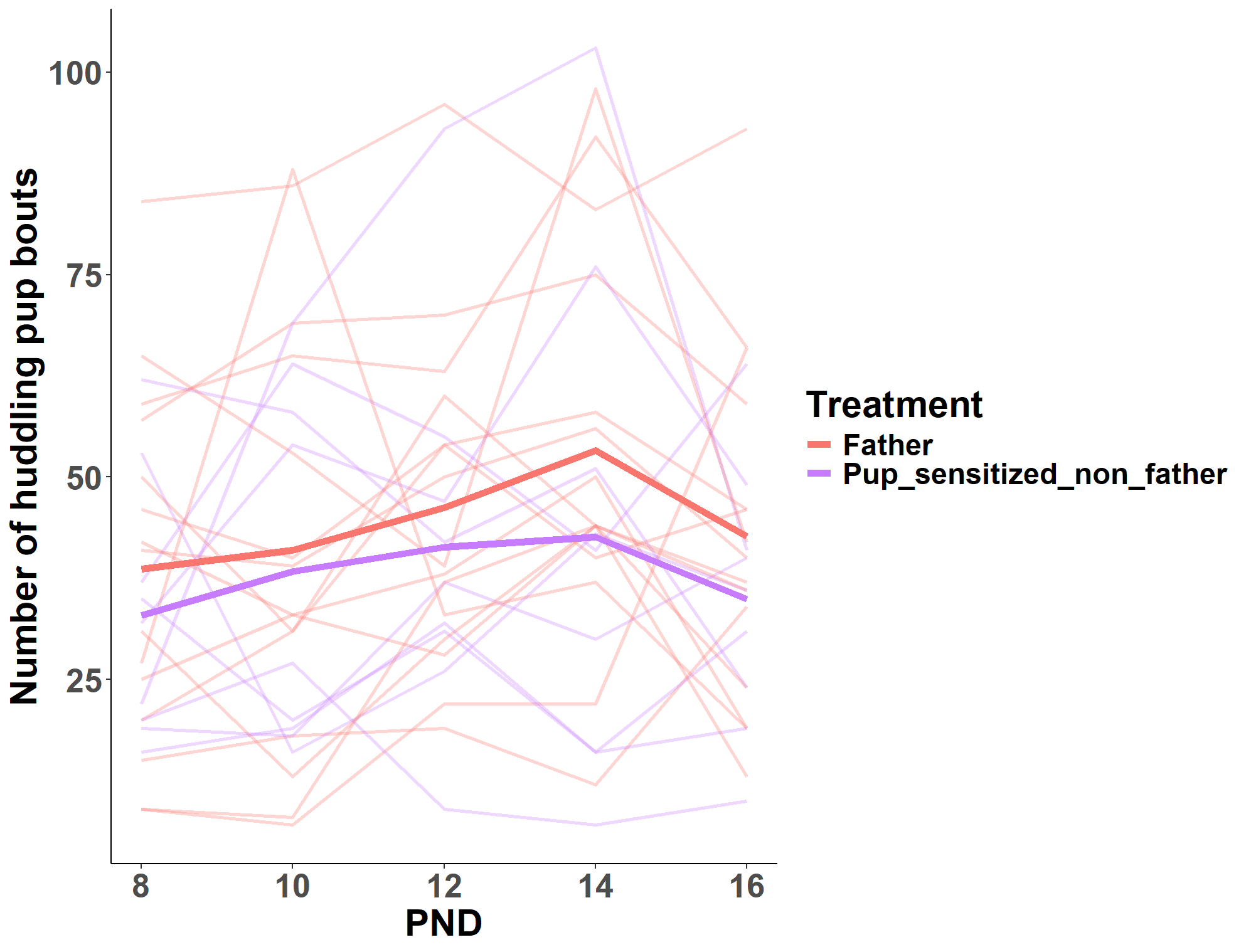

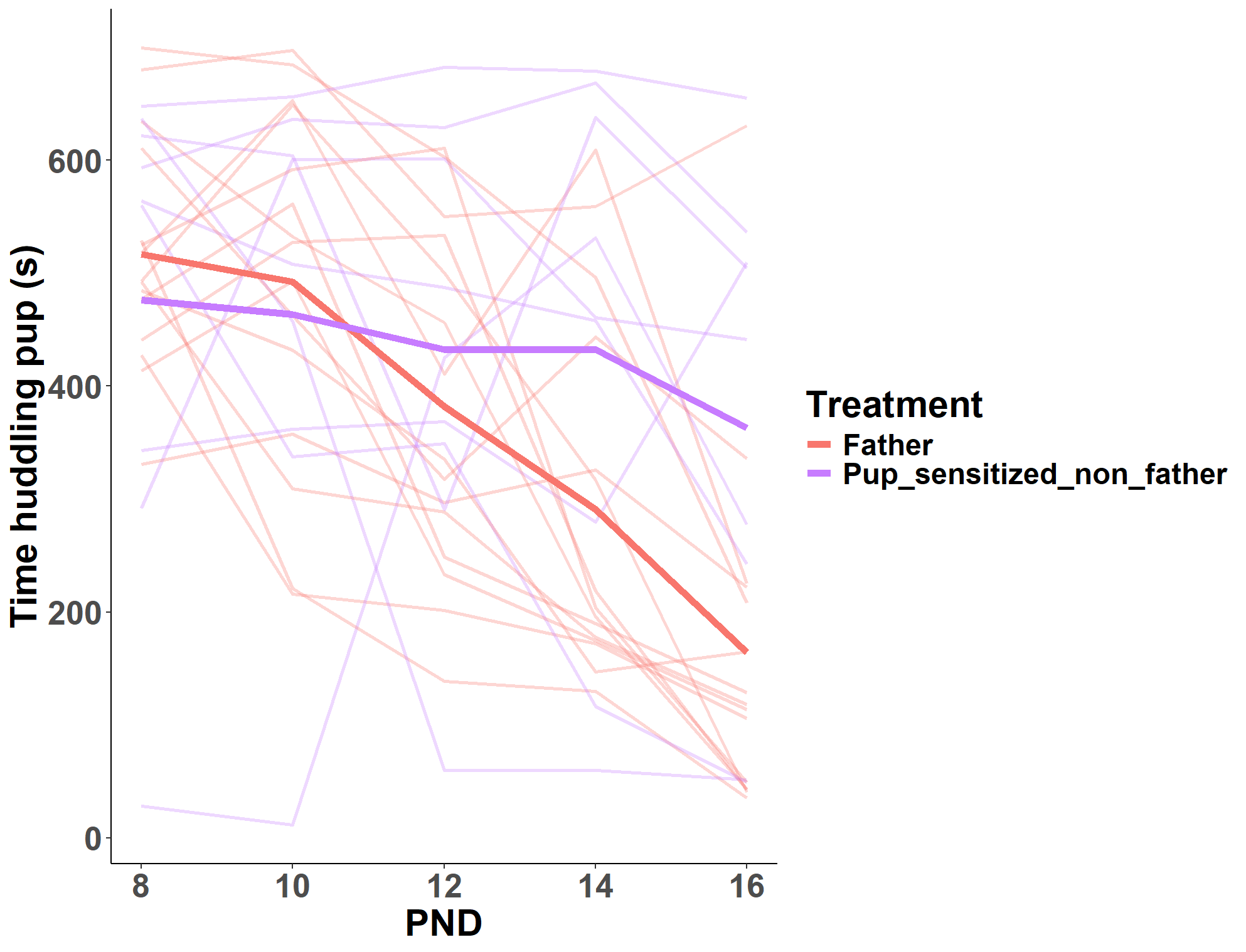

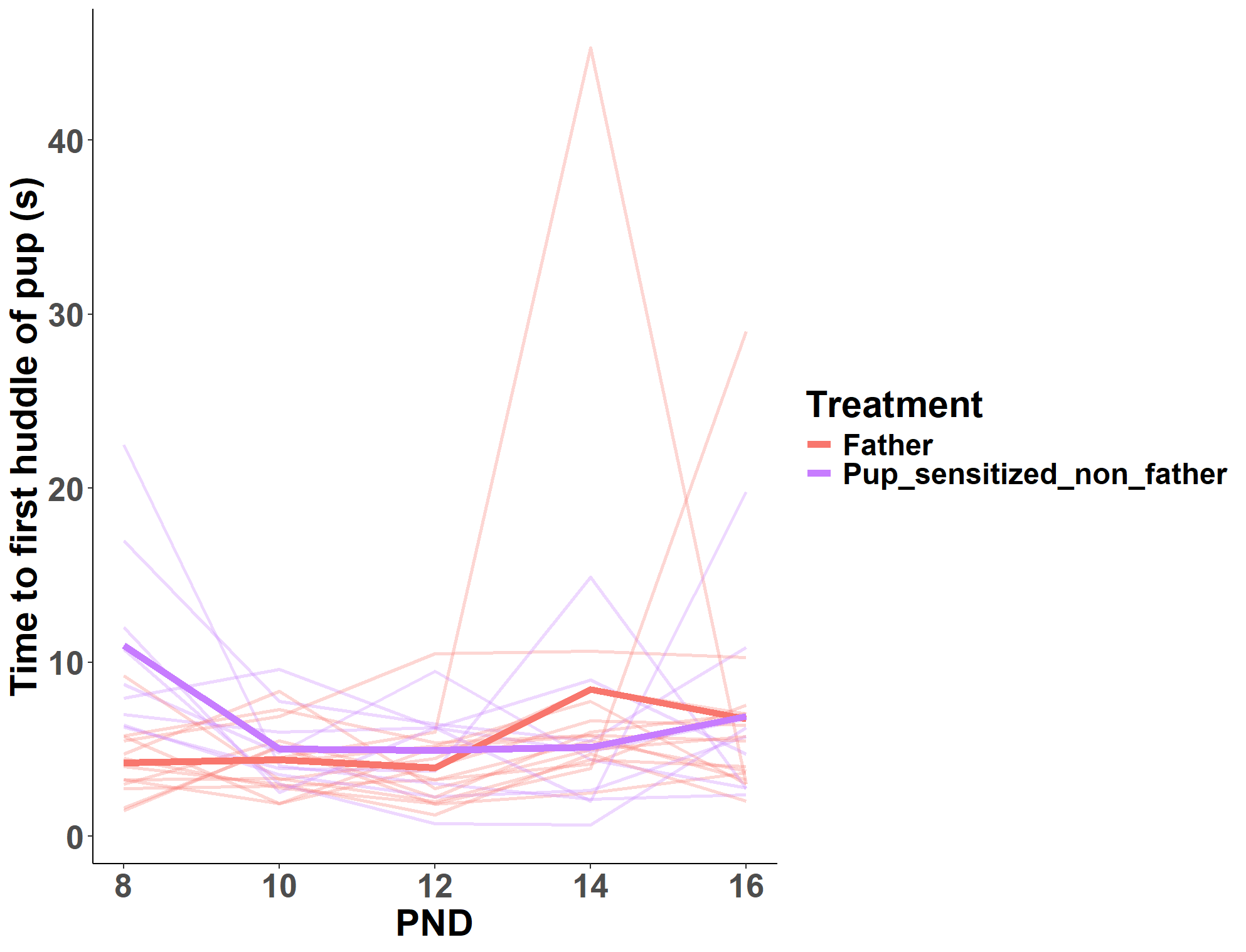

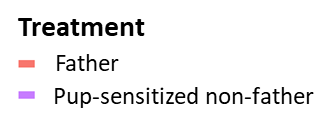

Pup-sensitized non-father

First-time father

**FREQUENCY**

**DURATION**

**LATENCY**

**F**

**E**

**C**

**B**

**D**

**A**

**Figure S19.** Huddling **(A)** latency, **(B)** duration, and **(C)** frequency and grooming **(D)** latency, **(E)** duration, and **(F)** frequency of pup-sensitized non-fathers and first-time fathers during the caregiving assay. Thin lines represent individuals and bold lines represent experience means.

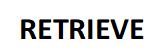

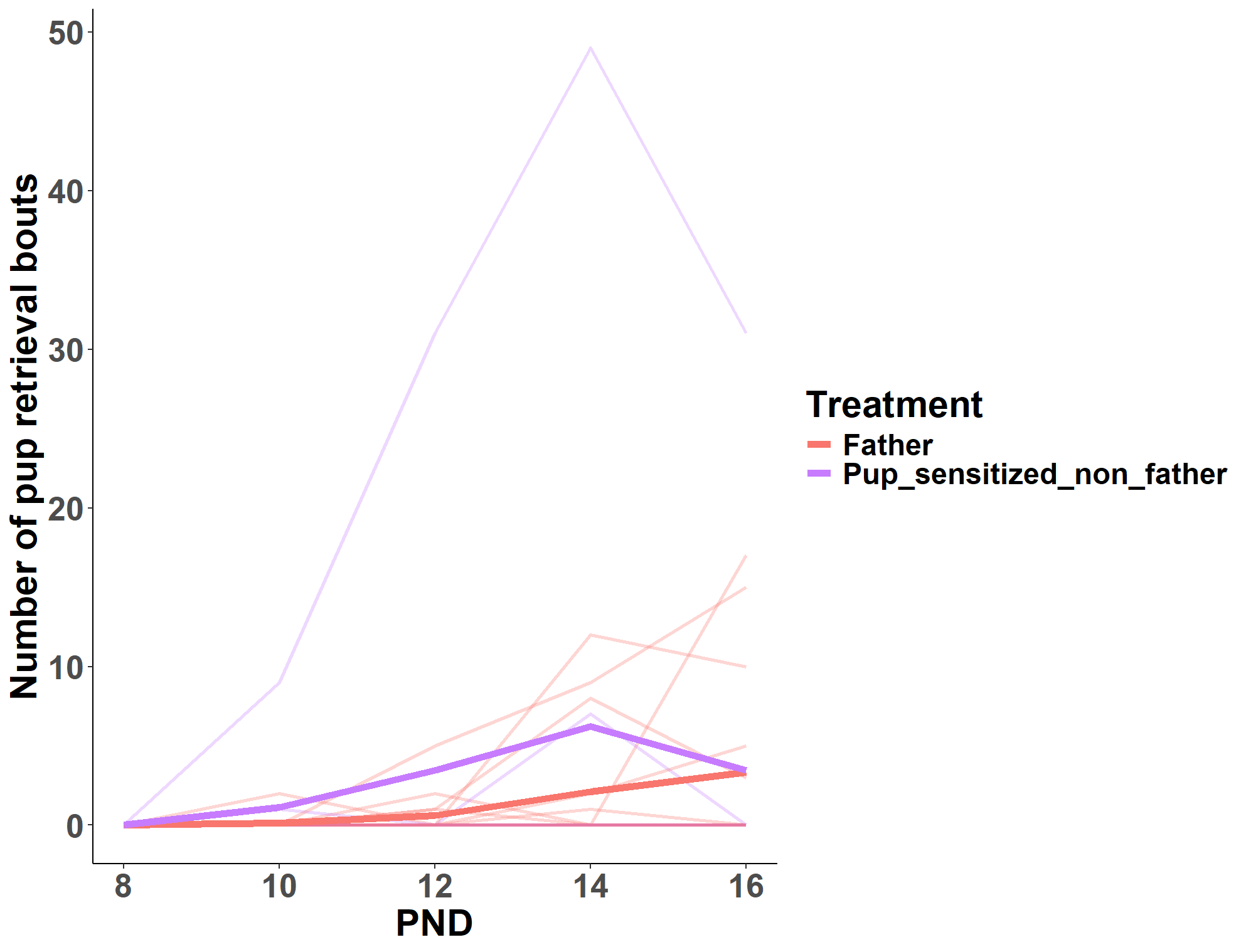

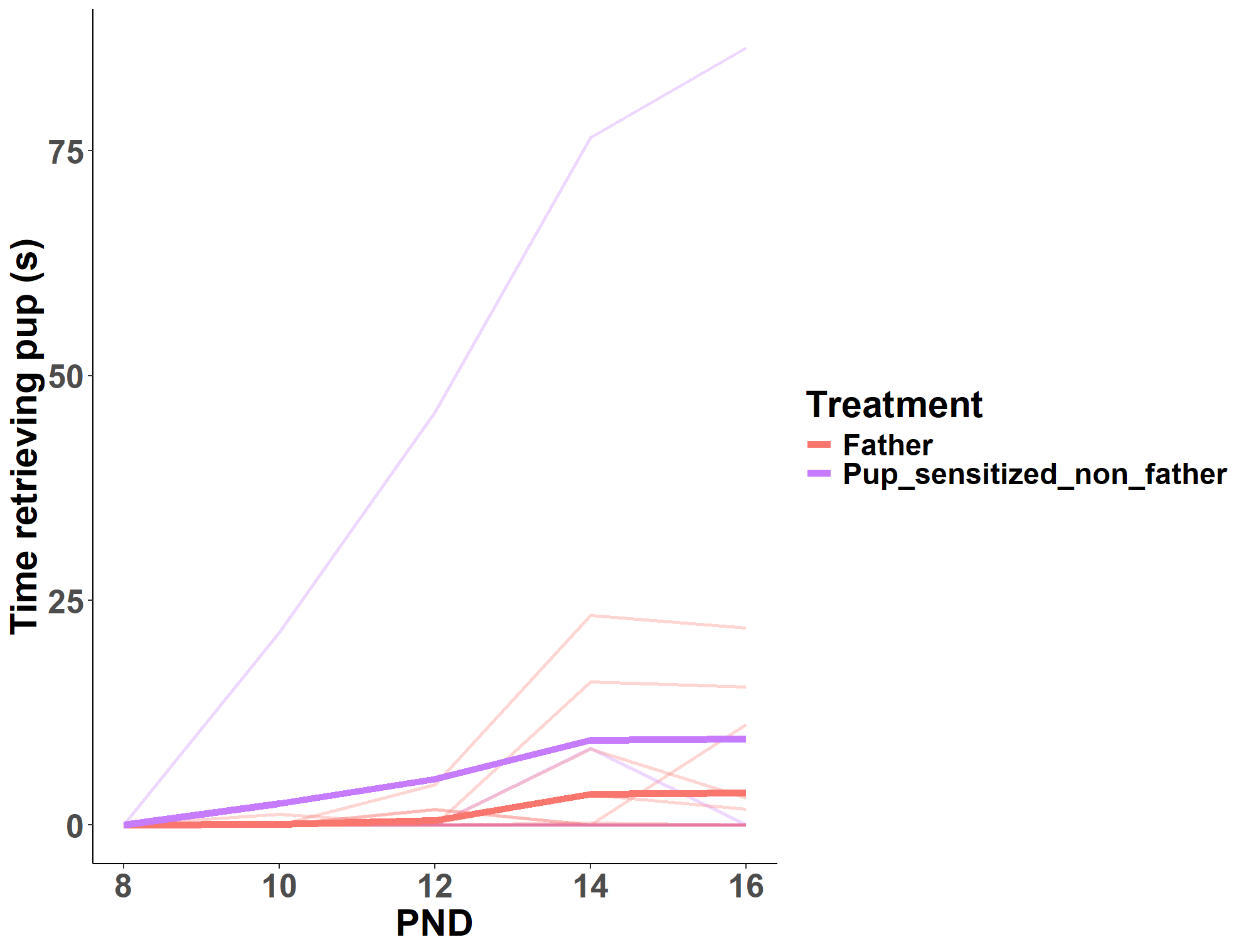

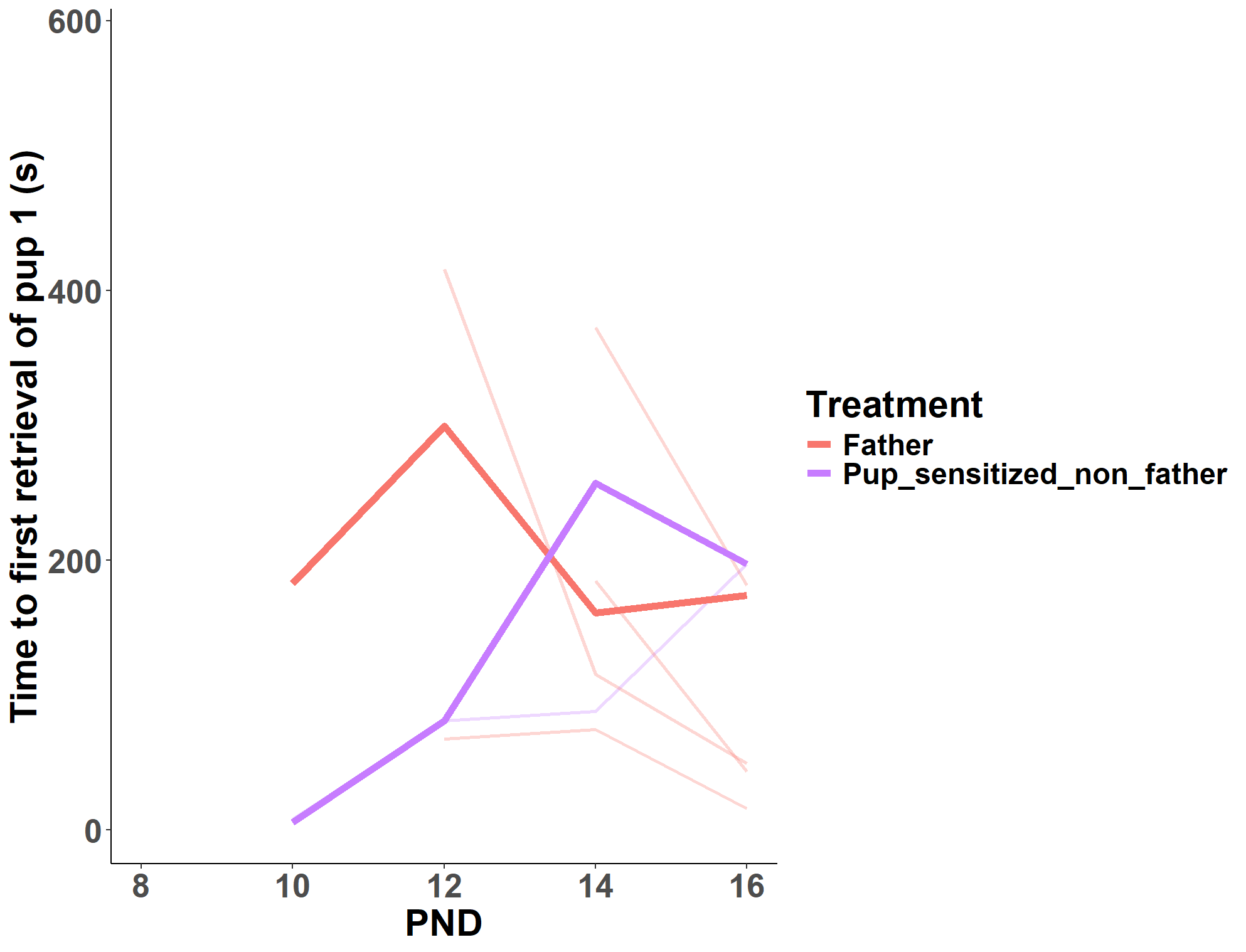

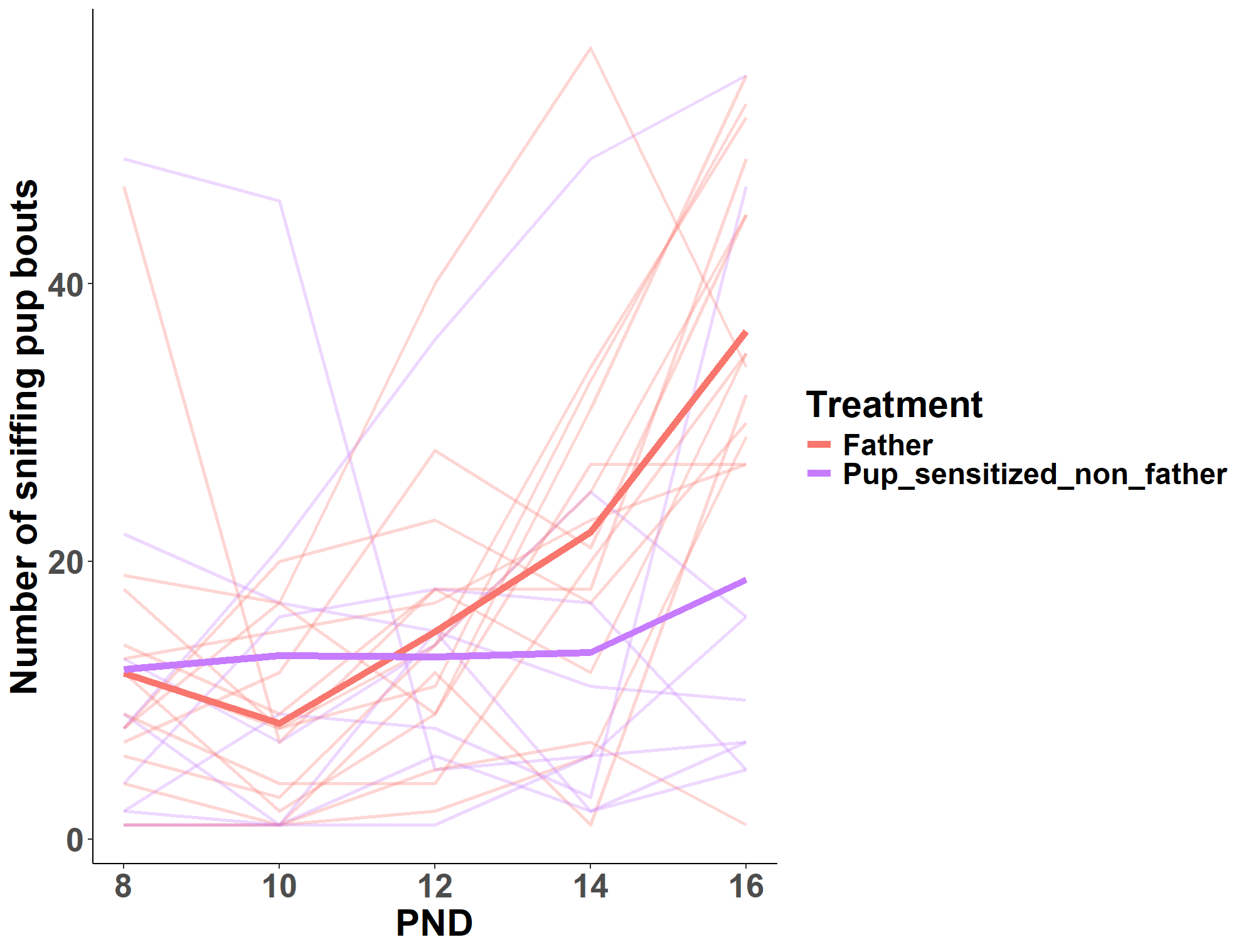

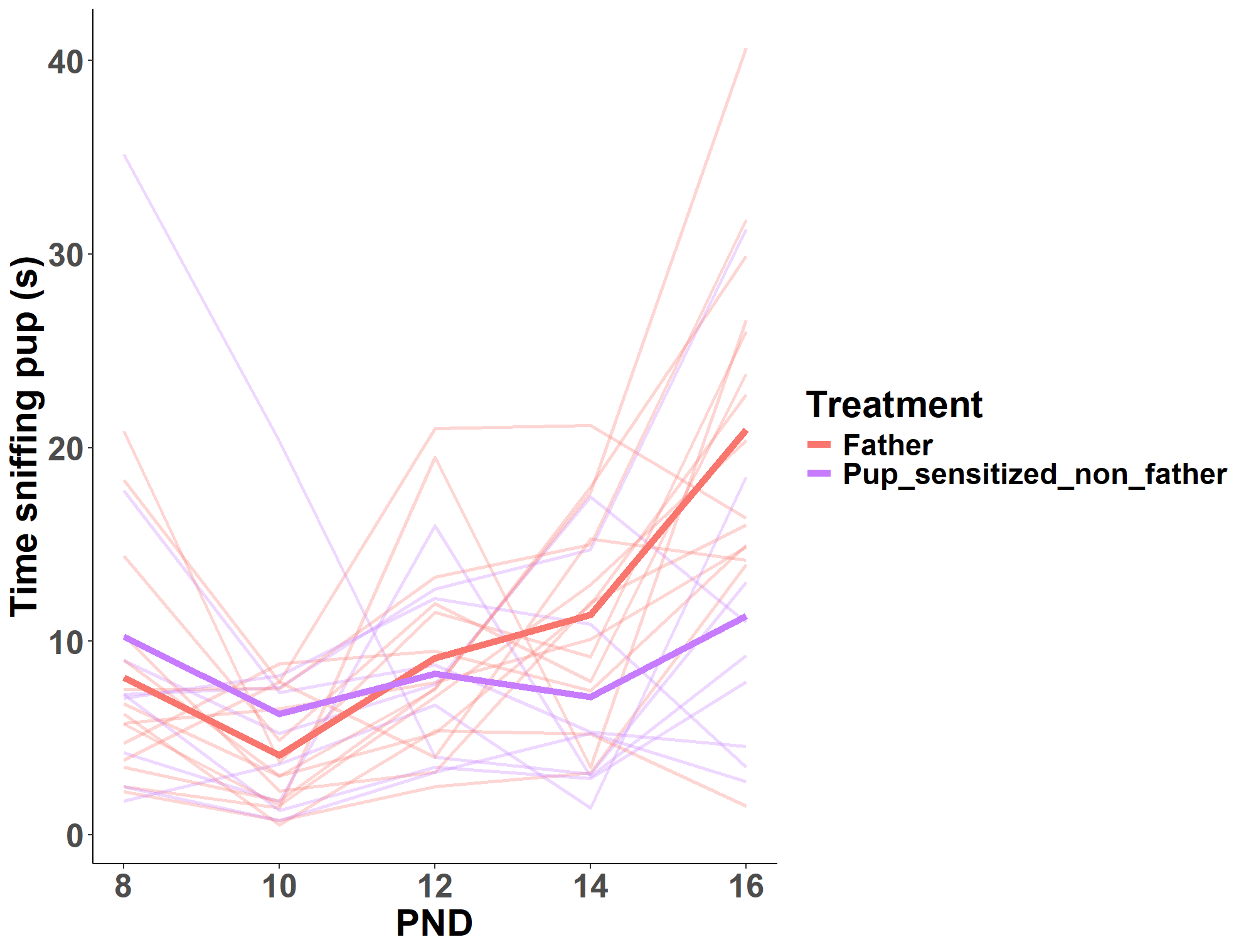

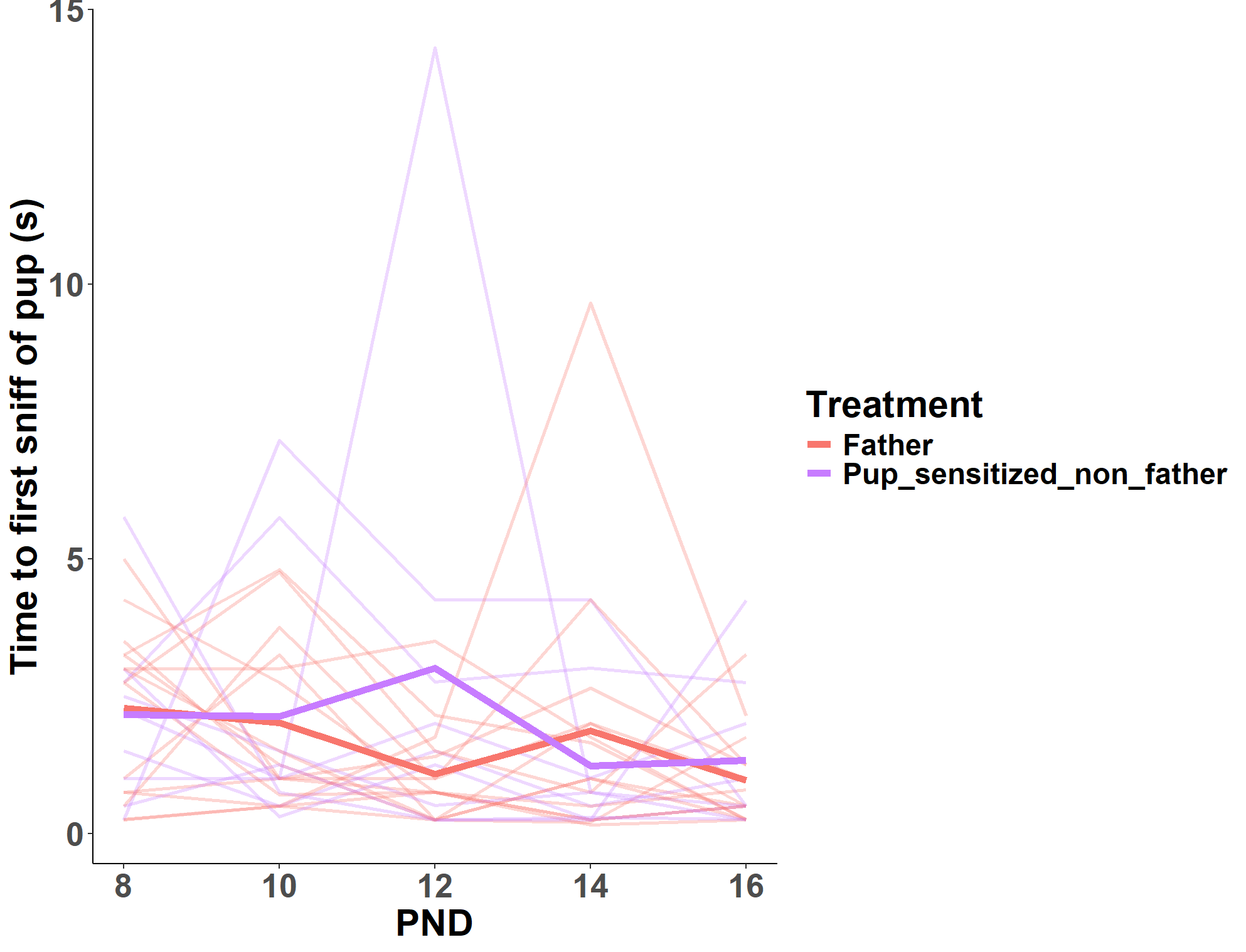

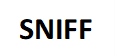
**Figure S20.** Sniffing **(A)** latency, **(B)** duration, and **(C)** frequency and retrieving **(D)** latency, **(E)** duration, and **(F)** frequency of pup-sensitized non-fathers and first-time fathers during the caregiving assay. Thin lines represent individuals and bold lines represent experience means.

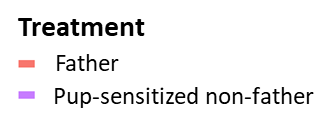

Pup-sensitized non-father

First-time father

**FREQUENCY**

**DURATION**

**LATENCY**

**F**
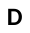

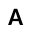

**E**
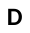

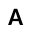

**D**
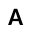

**C**

**B**

**A**

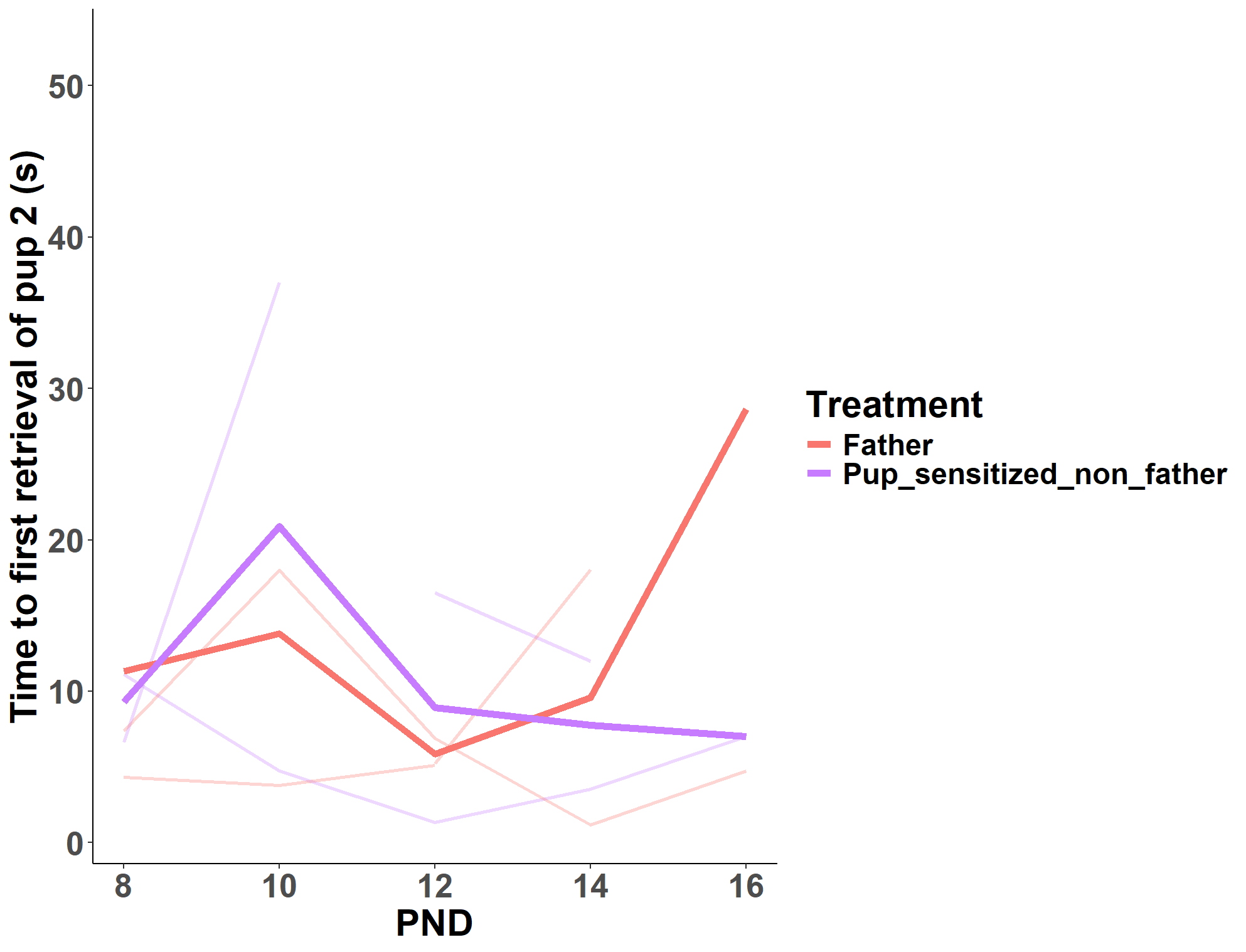
**Figure S21.** Latency to retrieval of second pup during the caregiving assay between pup-sensitized non-fathers and first-time fathers. Thin lines represent individuals and bold lines represent experience means.

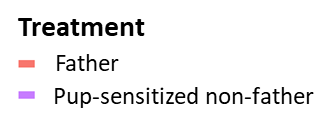

Pup-sensitized non-father

First-time father

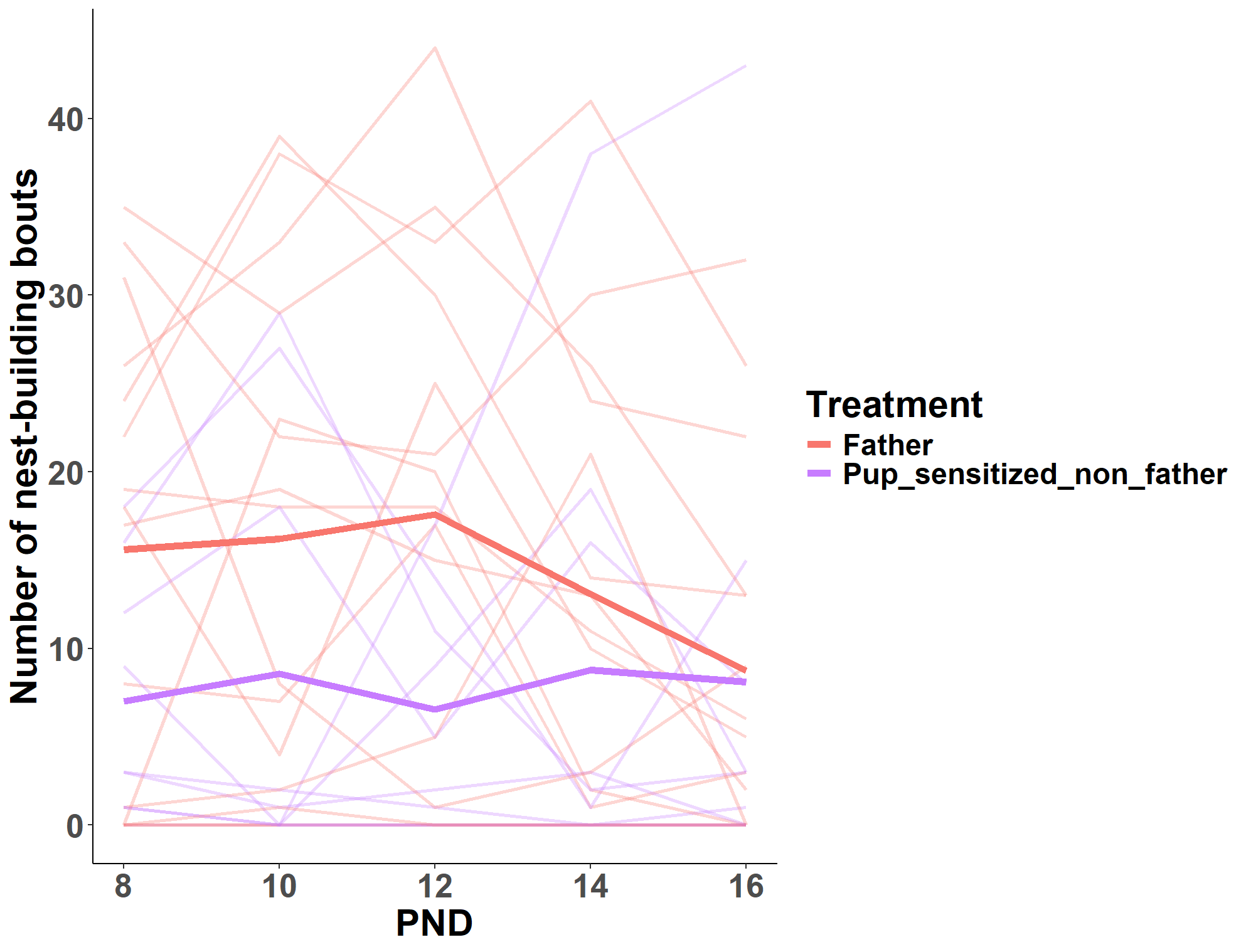

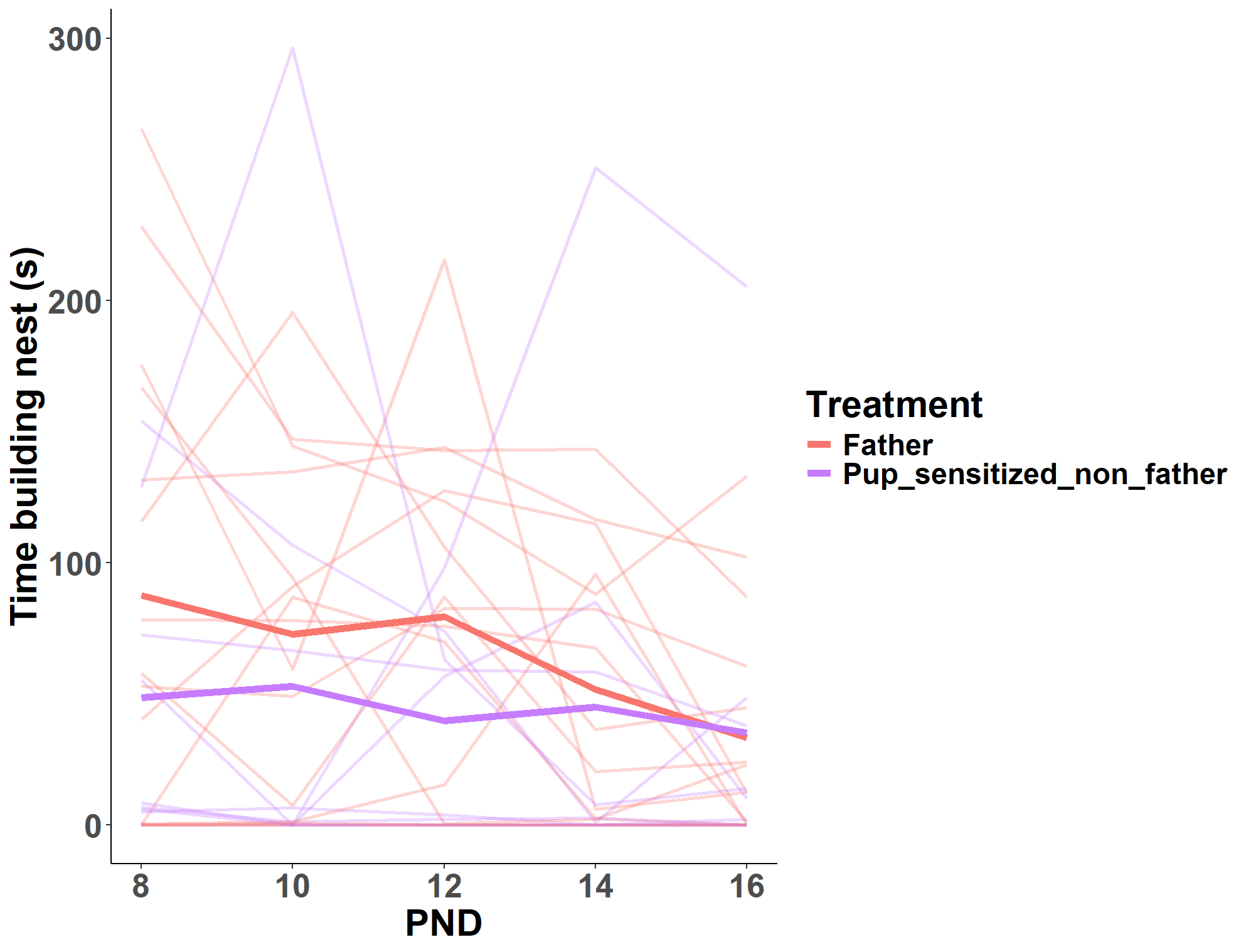
**Figure S22.** Nest building behavior (A) duration and (B) frequency of pup-sensitized non-fathers and first-time fathers during the caregiving assay. Thin lines represent individuals and bold lines represent experience means.

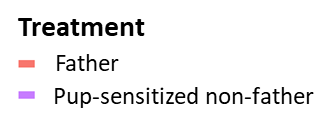

Pup-sensitized non-father

First-time father

**B**

**A**

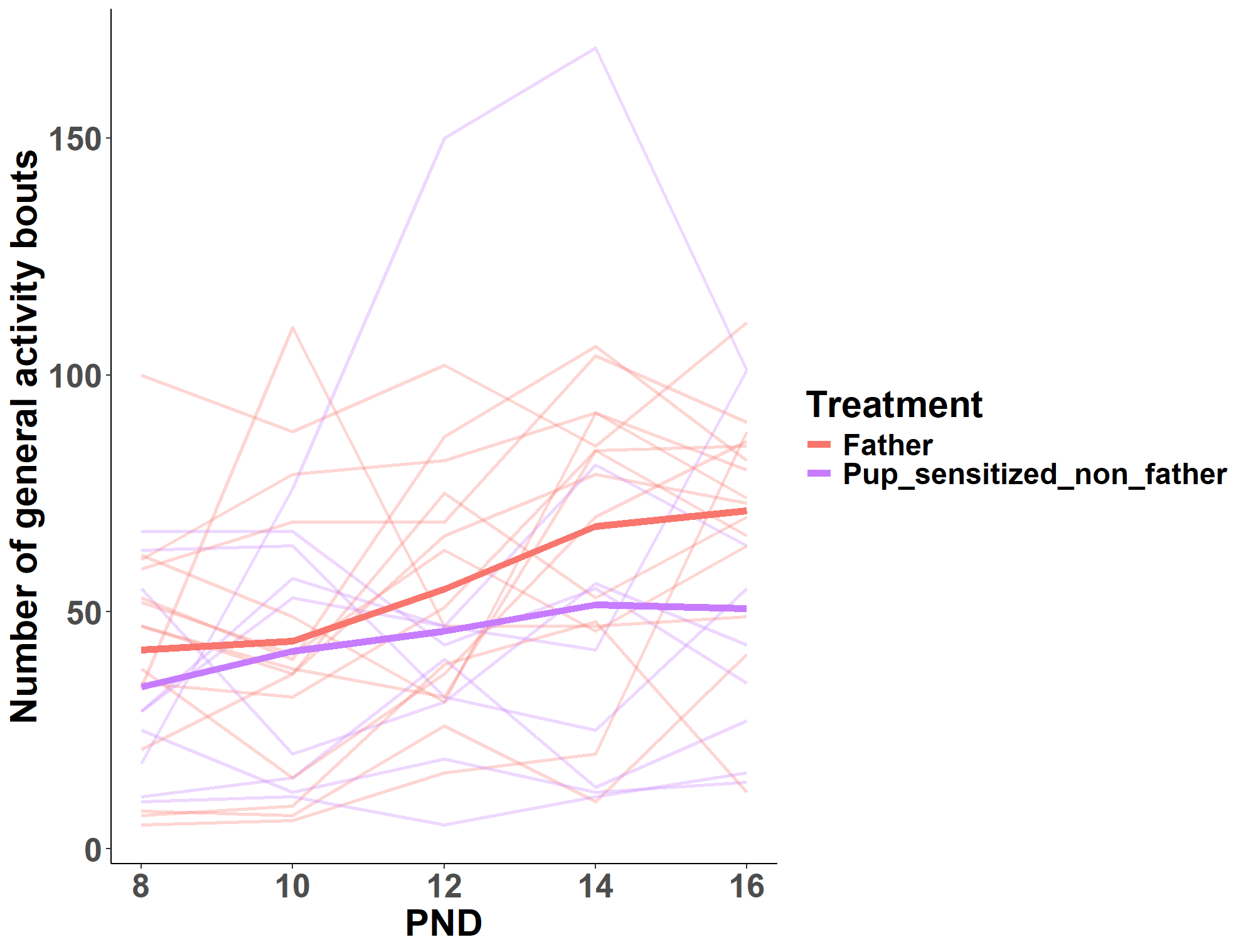

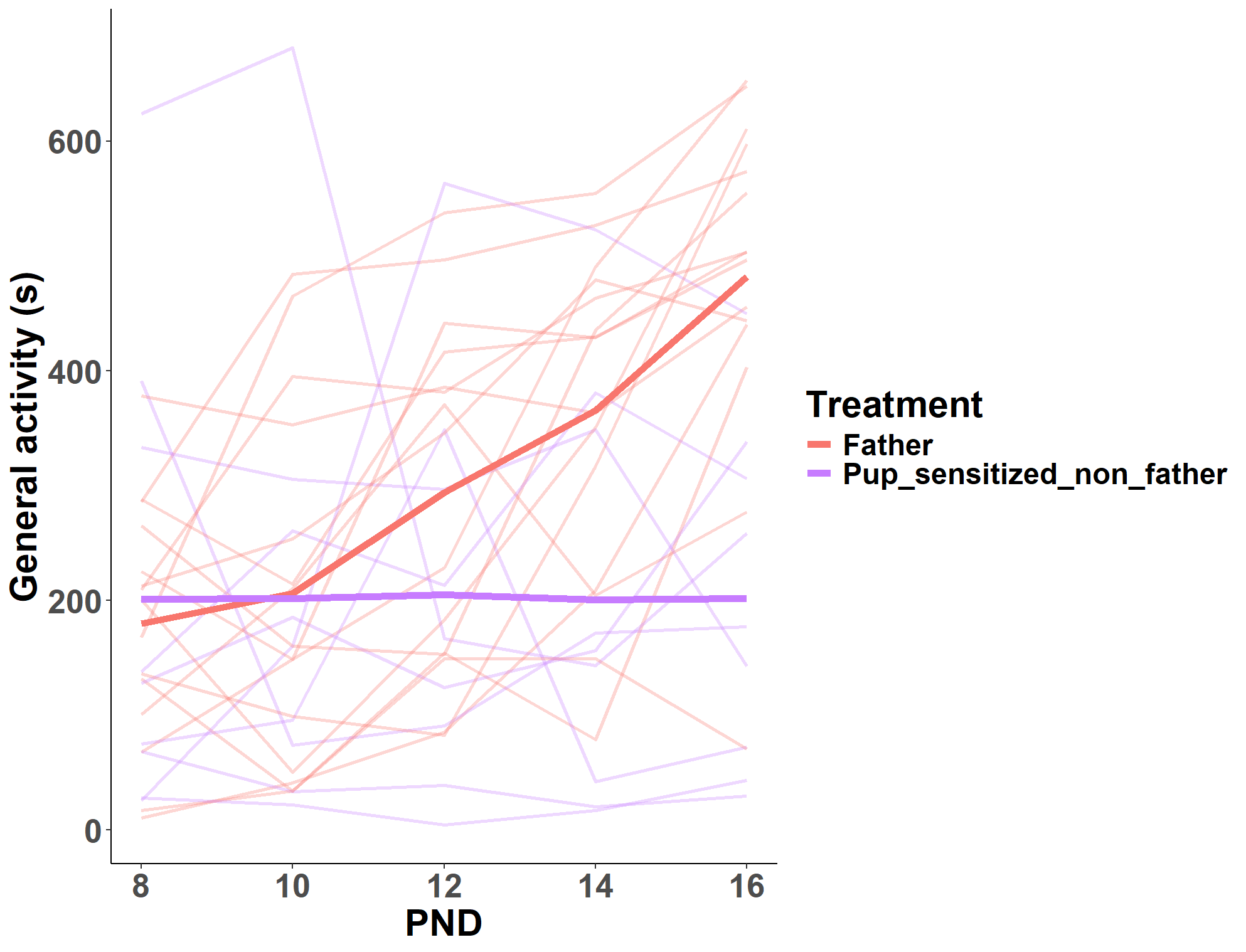
**Figure S23.** General activity **(A)** duration and **(B)** frequency behavior of pup-sensitized non-fathers and first-time fathers during the caregiving assay. Thin lines represent individuals and bold lines represent experience means.

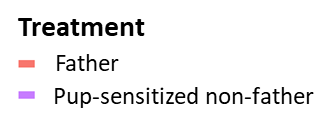

Pup-sensitized non-father

First-time father

**A**

**B**

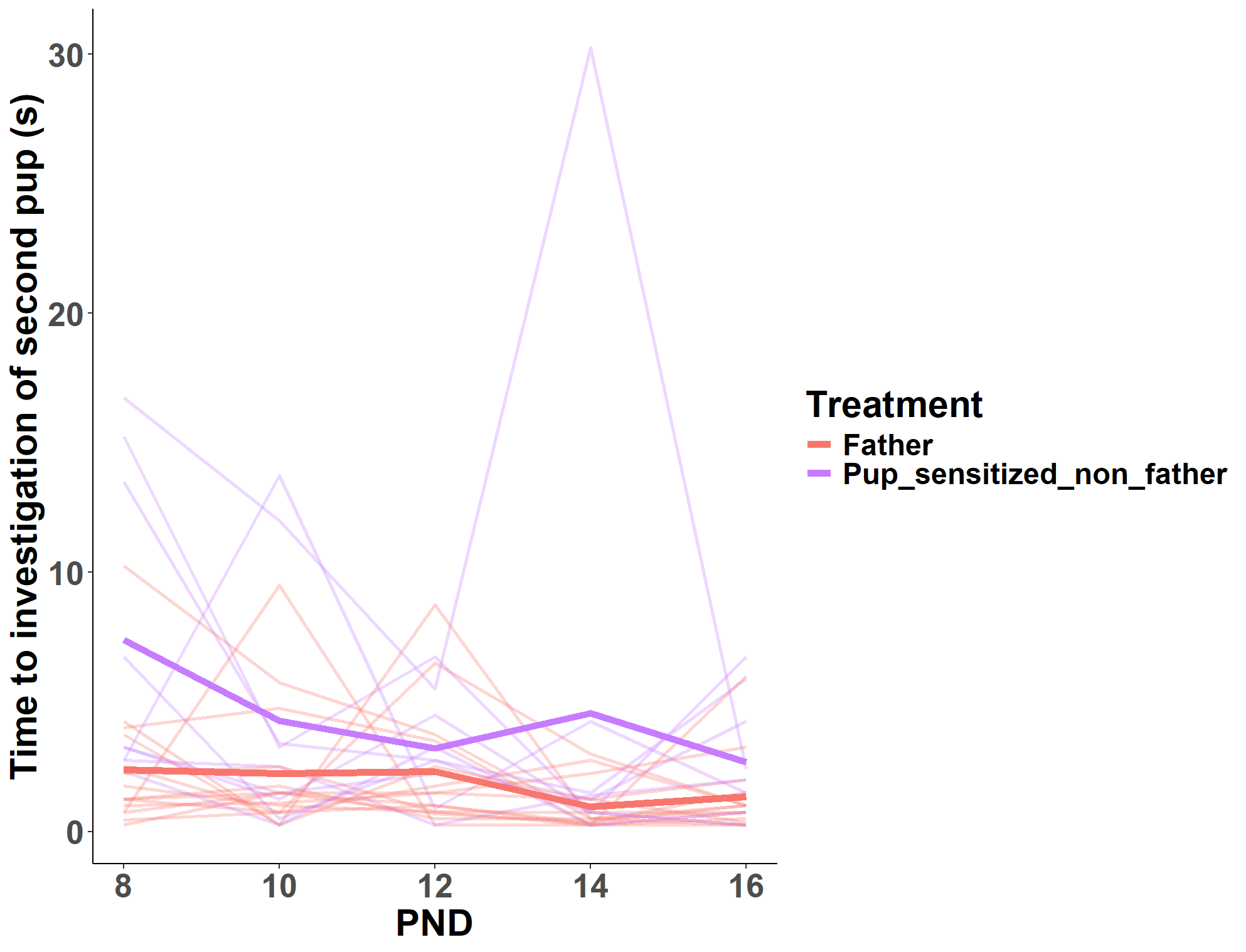

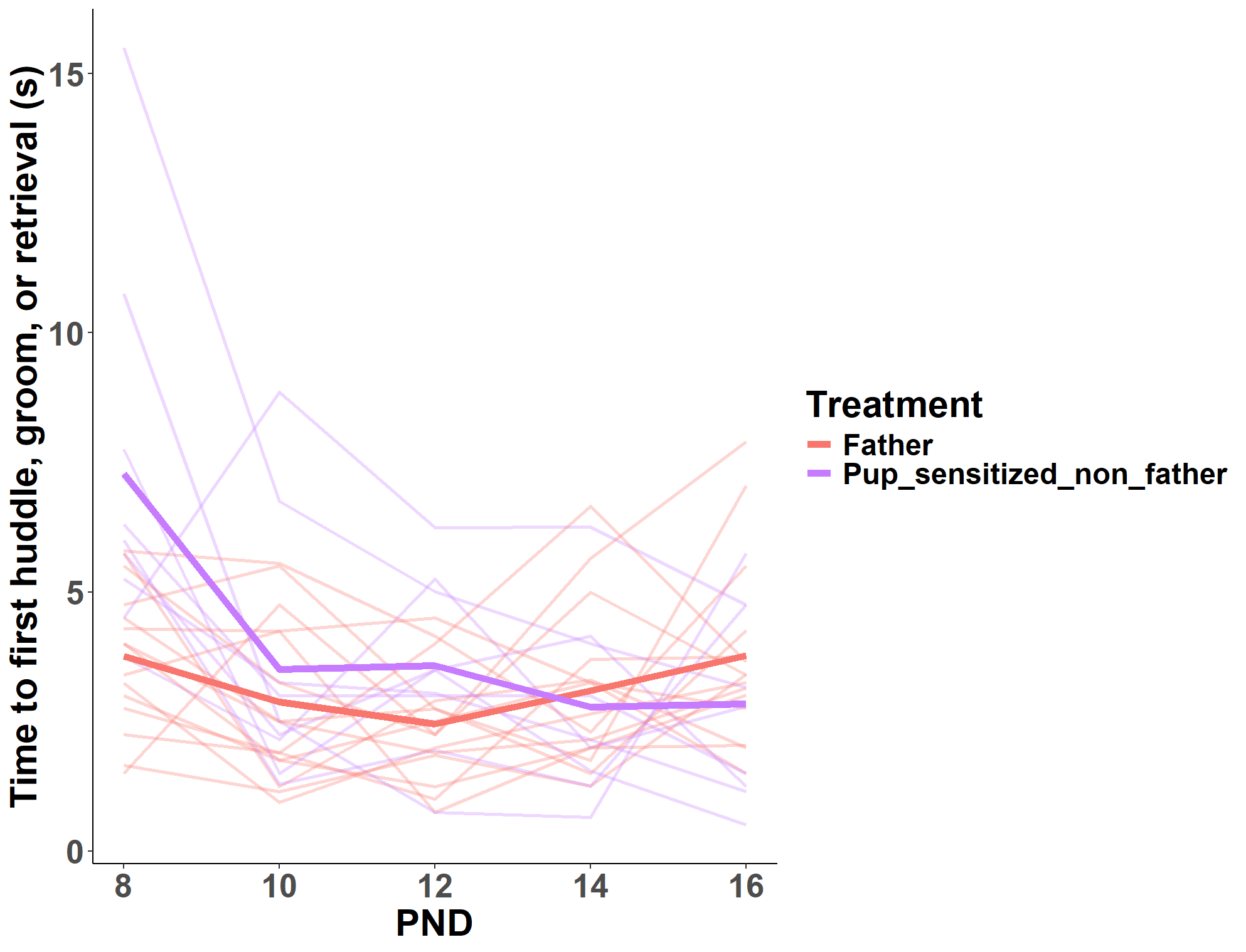
**Figure S24.** **(A)** Parental behavior latency and **(B)** pup investigation latency in pup-sensitized non-fathers and first-time fathers during the caregiving assay. Thin lines represent individuals and bold lines represent experience means.

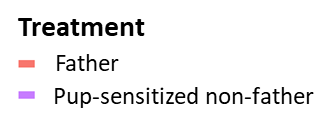

Pup-sensitized non-father

First-time father

**B**

**A**

**Supplemental Tables**

**Table S1.** Pup-exposure assay inter-rater reliability results. Intra-class correlations between scores of four independent coders on a subset of 21 videos and between the scores of the primary observer on a subset of five re-scored videos.

|  | Exploration near | Exploration away | Pup-directed activity | Pup-directed activity latency | Resting | Grooming | Jumping |
| --- | --- | --- | --- | --- | --- | --- | --- |
| Other coders | 0.78 | 0.98 | 0.93 | 1.00 | 0.37 | 0.90 | 0.98 |
| Self-check | 0.99 | 0.98 | 0.83 | 0.99 | NaN | 0.94 | 0.96 |

**Table S2.** Caregiving assay inter-rater reliability results. Intra-class correlations between scores of four independent coders on a subset of 21 videos and between the scores of the primary observer on a subset of five re-scored videos.

|  | Nest building | Parental behavior latency | Huddle latency | Groom latency | Sniff latency | Huddling | Grooming | Sniffing | Retrieving | General activity | Investigate latency | Retrieve latency 1 | Retrieve latency 2 |
| --- | --- | --- | --- | --- | --- | --- | --- | --- | --- | --- | --- | --- | --- |
| Other coders | 0.99 | 0.95 | 0.81 | 0.94 | 0.855 | 1.00 | 0.90 | 0.81 | 0.98 | 0.98 | 1.00 | 0.79 | 0.94 |
| Self-check | 1.00 | 0.99 | 1.00 | 0.99 | 0.939 | 1.00 | 0.98 | 0.89 | 1.00 | 1.00 | 1.00 | NA | NA |

**Table S3.** Novel object recognition test univariate linear models with experience as an effect in Experiment 1.

| **Response** | **Predictors** (P-value pairwise comparison) | **Estimated mean** | ***P* value** | **Adjusted R²** |
| --- | --- | --- | --- | --- |
| Difference score | Experiment: Virgin (v. Non-father) | 36.28 | 0.95 | 0.09 |
|  | Experiment: Non-father (v. Experienced father) | 38.89 | 0.12 |  |
|  | Experiment: Experienced father (v. Virgin) | 113.10 | 0.08 |  |
| Percent time with novel object | Experiment: Virgin (v. Non-father) | 66.09 | 0.16 | 0.25 |
|  | Experiment: Non-father (v. Experienced father) | 77.35 | 0.19 |  |
|  | Experiment: Experienced father (v. Virgin) | 89.82 | **0.01** |  |
| Time with novel object | Experiment: Virgin (v. Non-father) | 63.93 | 0.77 | 0.05 |
|  | Experiment: Non-father (v. Experienced father) | 52.54 | 0.12 |  |
|  | Experiment: Experienced father (v. Virgin) | 124.55 | 0.15 |  |
| Time with familiar object | Experiment: Virgin (v. Non-father) | 27.65 | 0.10 | 0.13 |
|  | Experiment: Non-father (v. Experienced father) | 13.66 | 0.82 |  |
|  | Experiment: Experienced father (v. Virgin) | 11.45 | 0.08 |  |

**Table S4.** Elevated plus maze test univariate linear models with experience as an effect in Experiment 1.

| **Response** | **Predictors** (P-value pairwise comparison) | **Estimated mean** | ***P* value** | **Adjusted R²** |
| --- | --- | --- | --- | --- |
| Number closed arm entries | Experiment: Virgin (v. Non-father) | 40.91 | **0.05** | 0.14 |
|  | Experiment: Non-father (v. Experienced father) | 60.80 | **0.05** |  |
|  | Experiment: Experienced father (v. Virgin) | 39.29 | 0.85 |  |
| Time in closed arms | Experiment: Virgin (v. Non-father) | 158.65 | 0.16 | 0.34 |
|  | Experiment: Non-father (v. Experienced father) | 199.28 | **0.002** |  |
|  | Experiment: Experienced father (v. Virgin) | 91.54 | **0.02** |  |
| Number open arm entries | Experiment: Virgin (v. Non-father) | 45.55 | 0.53 | 0.01 |
|  | Experiment: Non-father (v. Experienced father) | 36.40 | 0.17 |  |
|  | Experiment: Experienced father (v. Virgin) | 58.43 | 0.32 |  |
| Time in open arms | Experiment: Virgin (v. Non-father) | 115.55 | 0.23 | 0.32 |
|  | Experiment: Non-father (v. Experienced father) | 80.38 | **0.003** |  |
|  | Experiment: Experienced father (v. Virgin) | 183.85 | **0.02** |  |
| Distance moved | Experiment: Virgin (v. Non-father) | 4412.0 | 0.84 | -0.06 |
|  | Experiment: Non-father (v. Experienced father) | 4550.8 | 0.85 |  |
|  | Experiment: Experienced father (v. Virgin) | 4692.0 | 0.64 |  |
| Percent time in open arms | Experiment: Virgin (v. Non-father) | 42.13 | 0.20 | 0.33 |
|  | Experiment: Non-father (v. Experienced father) | 28.35 | **0.003** |  |
|  | Experiment: Experienced father (v. Virgin) | 66.80 | **0.02** |  |

**Table S5.** Novel object recognition univariate linear models with experience as an effect in Experiment 2.

| **Response** | **Predictors** (P-value pairwise comparison) | **Estimate mean** | ***P* value** | **Adjusted R²** |
| --- | --- | --- | --- | --- |
| Difference score | Experience: Non-father (v. First-time father) | 48.60 | 0.65 | -0.02 |
|  | Experience: Pup-sensitized non-father (v. Non-father) | 55.97 | 0.64 |  |
|  | Experience: First-time father (v. Pup-sensitized non-father) | 42.04 | 0.27 |  |
| Percent time with novel object | Experience: Non-father (v. First-time father) | 75.36 | 0.74 | -0.06 |
|  | Experience: Pup-sensitized non-father (v. Non-father) | 75.85 | 0.93 |  |
|  | Experience: First-time father (v. Pup-sensitized non-father) | 73.64 | 0.62 |  |
| Time with novel object | Experience: Non-father (v. First-time father) | 72.39 | 0.56 | 0.01 |
|  | Experience: Pup-sensitized non-father (v. Non-father) | 83.61 | 0.51 |  |
|  | Experience: First-time father (v. Pup-sensitized non-father) | 63.46 | 0.14 |  |
| Time with familiar object | Experience: Non-father (v. First-time father) | 23.79 | 0.75 | -0.04 |
|  | Experience: Pup-sensitized non-father (v. Non-father) | 27.631 | 0.63 |  |
|  | Experience: First-time father (v. Pup-sensitized non-father) | 21.413 | 0.33 |  |

**Table S6.** Elevated plus maze test univariate linear models with experience as an effect in Experiment 2.

| **Response** | **Predictors** (P-value pairwise comparison) | **Estimated mean** | ***P* value** | **Adjusted R²** |
| --- | --- | --- | --- | --- |
| Number closed arm entries | Experience: Non-father (v. First-time father) | 16.60 | 0.51 | -0.04 |
|  | Experience: Pup-sensitized non-father (v. Non-father) | 20.00 | 0.39 |  |
|  | Experience: First-time father (v. Pup-sensitized non-father) | 18.87 | 0.76 |  |
| Time in closed arms | Experience: Non-father (v. First-time father) | 75.38 | 0.23 | -0.01 |
|  | Experience: Pup-sensitized non-father (v. Non-father) | 86.54 | 0.68 |  |
|  | Experience: First-time father (v. Pup-sensitized non-father) | 103.14 | 0.50 |  |
| Latency to enter closed arms | Experience: Non-father (v. First-time father) | 51.43 | 0.54 | 0.002 |
|  | Experience: Pup-sensitized non-father (v. Non-father) | 25.51 | 0.16 |  |
|  | Experience: First-time father (v. Pup-sensitized non-father) | 41.80 | 0.34 |  |
| Number open arm entries | Experience: Non-father (v. First-time father) | 28.20 | 0.70 | 0.04 |
|  | Experience: Pup-sensitized non-father (v. Non-father) | 21.38 | 0.19 |  |
|  | Experience: First-time father (v. Pup-sensitized non-father) | 29.93 | 0.08 |  |
| Time in open arms | Experience: Non-father (v. First-time father) | 188.50 | 0.37 | -0.04 |
|  | Experience: Pup-sensitized non-father (v. Non-father) | 176.82 | 0.70 |  |
|  | Experience: First-time father (v. Pup-sensitized non-father) | 165.09 | 0.67 |  |
| Latency to enter open arms | Experience: Non-father (v. First-time father) | 6.96E-16 | 0.29 | -0.02 |
|  | Experience: Pup-sensitized non-father (v. Non-father) | 0.88 | 0.82 |  |
|  | Experience: First-time father (v. Pup-sensitized non-father) | 3.54 | 0.46 |  |
| Distance moved | Experience: Non-father (v. First-time father) | 4733.10 | 0.60 | 0.07 |
|  | Experience: Pup-sensitized non-father (v. Non-father) | 3782.10 | 0.15 |  |
|  | Experience: First-time father (v. Pup-sensitized non-father) | 5015.60 | **0.05** |  |
| Percent time in open arms | Experience: Non-father (v. First-time father) | 62.70 | 0.29 | -0.03 |
|  | Experience: Pup-sensitized non-father (v. Non-father) | 57.78 | 0.61 |  |
|  | Experience: First-time father (v. Pup-sensitized non-father) | 53.68 | 0.65 |  |

**Table S7.** Reproductive traits univariate linear models with experience as an effect in Experiment 1.

| **Response** | **Predictors** (P-value pairwise comparison) | **Estimated mean** | **P value** | **Adjusted R²** |
| --- | --- | --- | --- | --- |
| Average testis weight | Experience: Virgin (v. Non-father) | 168.04 | 0.12 | 0.09 |
|  | Experience: Non-father (v. Experienced father) | 204.86 | **0.04** |  |
|  | Experience: Experienced father (v. Virgin) | 147.86 | 0.36 |  |
| Sperm count | Experience: Virgin (v. Non-father) | 8993006 | **0.01** | 0.22 |
|  | Experience: Non-father (v. Experienced father) | 23539928 | 0.12 |  |
|  | Experience: Experienced father (v. Virgin) | 14812499 | 0.16 |  |

**Table S8.** Reproductive traits univariate linear models with experience as an effect in Experiment 2.

| **Response** | **Predictors** (P-value pairwise comparison) | **Estimated mean** | **P-value** | **Adjusted R²** |
| --- | --- | --- | --- | --- |
| Average testis weight | Experience: **Non-father** (v. First-time father) | 171.11 | 0.14 | 0.08 |
|  | Experience: **Pup-sensitized non-father** (v. Non-father) | 157.85 | 0.58 |  |
|  | Experience: **First-time father** (v. Pup-sensitized non-father) | 202.77 | 0.04 |  |
| Sperm count | Experience: **Non-father** (v. First-time father) | 5021296 | 0.38 | -0.02 |
|  | Experience: **Pup-sensitized non-father** (v. Non-father) | 4950617 | 0.96 |  |
|  | Experience: **First-time father** (v. Pup-sensitized non-father) | 6008889 | 0.36 |  |

**Table S9.** Pup-exposure assay best fitting linear mixed effects models with experience and male age as fixed effects.

| **Response** | **Fixed effects** | **Estimate** | **SE** | **t-value** | ***P* value** | **Random effect** | **Variance** | **SD** |
| --- | --- | --- | --- | --- | --- | --- | --- | --- |
| Exploration near pup enclosure duration | Intercept | 154.79 | 9.14 | 16.94 | 4.14E-14 | Subject | 927.8 | 30.46 |
|  | Experience | -3.11 | 14.92 | -0.21 | 0.84 | Residual | 1621.2 | 40.26 |
| Number of bouts of exploration near pup enclosure | Intercept | 233.23 | 110.74 | 2.11 | 0.04 | Subject | 2123.6 | 46.08 |
|  | Experience | 0.04 | 21.64 | 0.002 | 1.00 | Residual | 812.8 | 28.51 |
|  | Male age | -0.99 | 0.70 | -1.41 | 0.17 |  | | |
| Exploration away from pup enclosure duration | Intercept | 82.13 | 9.77 | 8.41 | 2.55E-08 | Subject | 1171 | 34.22 |
|  | Experience | 7.66 | 15.95 | 0.48 | 0.64 | Residual | 1297 | 36.02 |
| Number of bouts of exploration away from pup enclosure | Intercept | 51.11 | 8.52 | 6.00 | 4.85E-06 | Subject | 960.8 | 31 |
|  | Experience | 4.34 | 13.91 | 0.31 | 0.76 | Residual | 633.8 | 25.18 |
| Investigating pup duration | Intercept | 103.83 | 9.93 | 10.45 | 5.36E-10 | Subject | 1213 | 34.83 |
|  | Experience | 1.47 | 16.22 | 0.09 | 0.93 | Residual | 1335 | 36.54 |
| Number of investigation of pup bouts | Intercept | 64.15 | 3.92 | 16.37 | 8.32E-14 | Subject | 173.6 | 13.17 |
|  | Experience | -4.88 | 6.40 | -0.76 | 0.45 | Residual | 283.3 | 16.83 |
| Latency to investigate pup | Intercept | 9.10 | 3.50 | 2.60 | 0.02 | Subject | 38.81 | 6.23 |
|  | Experience | 1.85 | 5.72 | 0.32 | 0.75 | Residual | 726.52 | 26.95 |

Note: The model with the lowest AICc was selected as the best fitting model. When intercept-only, null model is the best fit model, only the estimate of the intercept is reported. ‘Experience’ compares pup-sensitized non-fathers and first-time fathers.

**Table S10.** Caregiving assay best fitting linear mixed effects models with experience and male age as fixed effects.

| **Response** | **Fixed effects** | **Estimate** | **SE** | **t-value** | ***P* value** | **Random effect** | **Variance** | **SD** |
| --- | --- | --- | --- | --- | --- | --- | --- | --- |
| Earliest latency | Intercept | 1.65 |  | | | | | |
| Parental behavior latency | Intercept | 3.19 | 0.29 | 10.91 | 2.41E-10 | Subject | 0.44 | 0.67 |
|  | Experience | 0.81 | 0.48 | 1.69 | 0.11 | Residual | 4.20 | 2.05 |
| Huddle latency | Intercept | 5.56 | 0.63 | 8.83 | 1.17E-14 |  | | |
|  | Experience | 1.04 | 1.03 | 1.01 | 0.32 |  |  |  |
| Huddling duration | Intercept | 1175.14 | 378.97 | 3.10 | 0.01 | Subject | 19863 | 140.9 |
|  | Experience | 8.79 | 71.77 | 0.12 | 0.91 | Residual | 26482 | 162.7 |
|  | Age | -4.93 | 2.31 | -2.14 | **0.05** |  | | |
| Number of huddling bouts | Intercept | 44.35 | 4.77 | 9.29 | 4.52E-09 | Subject | 295.3 | 17.19 |
|  | Experience | -6.35 | 7.80 | -0.81 | 0.42 | Residual | 232.0 | 15.23 |
| Groom latency | Intercept | 3.57 | 0.35 | 10.36 | 6.37E-10 | Subject | 0.79 | 0.89 |
|  | Experience | 0.68 | 0.56 | 1.21 | 0.24 | Residual | 4.94 | 2.22 |
| Grooming duration | Intercept | -285.68 | 219.16 | -1.30 | 0.20 | Subject | 6402 | 80.01 |
|  | Experience | 129.57 | 41.24 | 3.14 | **0.01** | Residual | 9449 | 97.21 |
|  | Age | 2.85 | 1.33 | 2.14 | **0.04** |  | | |
| Number of grooming bouts | Intercept | -50.83 | 30.76 | -1.65 | 0.11 | Subject | 136 | 11.66 |
|  | Experience | 15.70 | 5.87 | 2.68 | **0.01** | Residual | 162.7 | 12.76 |
|  | Age | 0.53 | 0.19 | 2.84 | **0.01** |  | | |
| Sniff latency | Intercept | 1.77 |  | | | | | |
| Sniffing duration | Intercept | 10.73 | 1.01 | 10.67 | 3.64E-10 | Subject | 3.19 | 1.79 |
|  | Experience | -2.09 | 1.64 | -1.27 | 0.22 | Residual | 59.87 | 7.74 |
| Number of sniffing bouts | Intercept | 18.79 | 2.22 | 8.46 | 2.29E-08 | Subject | 36.75 | 6.06 |
|  | Experience | -4.65 | 3.63 | -1.28 | 0.21 | Residual | 185.75 | 13.63 |
| Retrieving duration | Intercept | 1.52 | 2.45 | 0.62 | 0.54 | Subject | 77.07 | 8.78 |
|  | Experience | 3.79 | 4.01 | 0.95 | 0.35 | Residual | 66.13 | 8.13 |
| Number of retrieval bouts | Intercept | 1.24 | 1.30 | 0.96 | 0.35 | Subject | 20.74 | 4.56 |
|  | Experience | 1.60 | 2.12 | 0.76 | 0.46 | Residual | 22.62 | 4.76 |
| Interaction with pup duration | Intercept | 1106.99 | 547.73 | 2.02 | 0.06 | Subject | 44889 | 211.9 |
|  | Experience | 125.88 | 105.37 | 1.20 | 0.26 | Residual | 48041 | 219.2 |
|  | Age | -3.34 | 3.33 | -1.00 | 0.33 |  | | |
| Nest building duration | Intercept | 392.14 | 112.36 | 3.49 | 0.002 | Subject | 1618 | 40.23 |
|  | Experience | -43.20 | 21.02 | -2.06 | **0.05** | Residual | 2654 | 51.52 |
|  | Age | -2.00 | 0.68 | -2.93 | **0.01** |  | | |
| Number of nest building bouts | Intercept | 65.46 | 21.97 | 2.98 | 0.01 | Subject | 78.21 | 8.84 |
|  | Experience | -9.98 | 4.31 | -2.32 | **0.03** | Residual | 67.01 | 8.19 |
|  | Age | -0.31 | 0.13 | -2.35 | **0.02** |  | | |
| General activity duration | Intercept | -33.02 | 315.78 | -0.11 | 0.92 | Subject | 12584 | 112.2 |
|  | Experience | -80.01 | 58.93 | -1.36 | 0.20 | Residual | 21526 | 146.7 |
|  | Age | 2.07 | 1.92 | 1.08 | 0.29 |  | | |
| Number of general activity bouts | Intercept | 56.00 | 6.20 | 9.03 | 7.53E-09 | Subject | 468.1 | 21.64 |
|  | Experience | -11.20 | 10.13 | -1.11 | 0.28 | Residual | 546.4 | 23.37 |
| Resting duration | Intercept | 0.56 | 2.63 | 0.21 | 0.83 | Subject | 25.29 | 5.028 |
|  | Experience | 11.28 | 4.29 | 2.63 | **0.02** | Residual | 391.74 | 19.79 |
| Investigate latency | Intercept | 1.85 | 0.62 | 2.98 | 0.01 | Subject | 3.41 | 1.85 |
|  | Experience | 2.57 | 1.02 | 2.54 | **0.02** | Residual | 11.90 | 3.45 |

Note: The model with the lowest AICc was selected as the best fitting model. When intercept-only, null model is the best fit model, only the estimate of the intercept is reported. ‘Experience’ compares pup-sensitized non-fathers and first-time fathers.
